# Exploring the substrate specificity and catalytic mechanism in a PAPS-independent ASST from *Salmonella typhimurium*

**DOI:** 10.64898/2026.10.08.757629

**Authors:** Nina M. Nimbona, Haya Obeid, Dilara Balci, Aurélie Joublin-Delavat, Anne Gladis, Lionel Cladière, Mehdi D. Davari, Cyrille Grandjean, Franck Daligault, Mirjam Czjzek

**Affiliations:** Sorbonne Université, CNRS, UMR8227, Laboratory of Integrative Biology of Marine Models, Station Biologique de Roscoff, Roscoff, 29680 France; Nantes Université, CNRS, US2B, UMR 6286, F-44000 Nantes, France; Department of Bioorganic Chemistry, Leibniz Institute of Plant Biochemistry, Weinberg 3, 06120 Halle, Germany

**Keywords:** Enzyme mechanism, substrate specificity, bacterial arylsulfate sulfotransferase, PAPS-independent sulfation, protein structure, molecular dynamics

## Abstract

Sulfation plays an important role in many metabolic processes such as substrate detoxification, solubilization or activation. The enzymes responsible for these reactions, sulfotransferases, occur in all domains of life and can be classified in two major, evolutionary independent groups, PAPS-dependent and PAPS independent or arysulfate sulfotransferases (ASST) enzymes. Besides the fact that they play key roles in biological functions, sulfotransferases can be interesting enzymatic tools to selectively sulfate specific positions of various metabolites. To this end we need to understand the relation between sequence, structure and function. Actually, eukaryotic PAPS dependent sulfotransferases have extensively been studied, while we largely lack knowledge about the diversity of bacterial PAPS-independent ASSTs. We have conducted a multidisciplinary study of a bacterial sulfotransferase from *Salmonella typhimurium,* revealing the importance of active site residues for activity and guiding substrate specificity. Based on the crystal structure in complex with substrate, we used molecular docking combined with Quantum Mechanics/Molecular Mechanics (QM/MM) free energy calculations to investigate the reaction catalytic mechanism with a focus on the role of specific active site residues in substrate specificity. Our results show that the catalytic mechanism involving three conserved histidines is common with previously determined ASSTs, and highlight a specific pair of aromatic residues, Phe174 and Tyr557, that play an important role in defining the substrate specificity in ASST clade 5. These findings enhance our general understanding of how substrate specificity is attained and has the potential to guide further enzyme engineering to expand the acceptor scope of ASSTs.

**TOC GRAPHIC:** 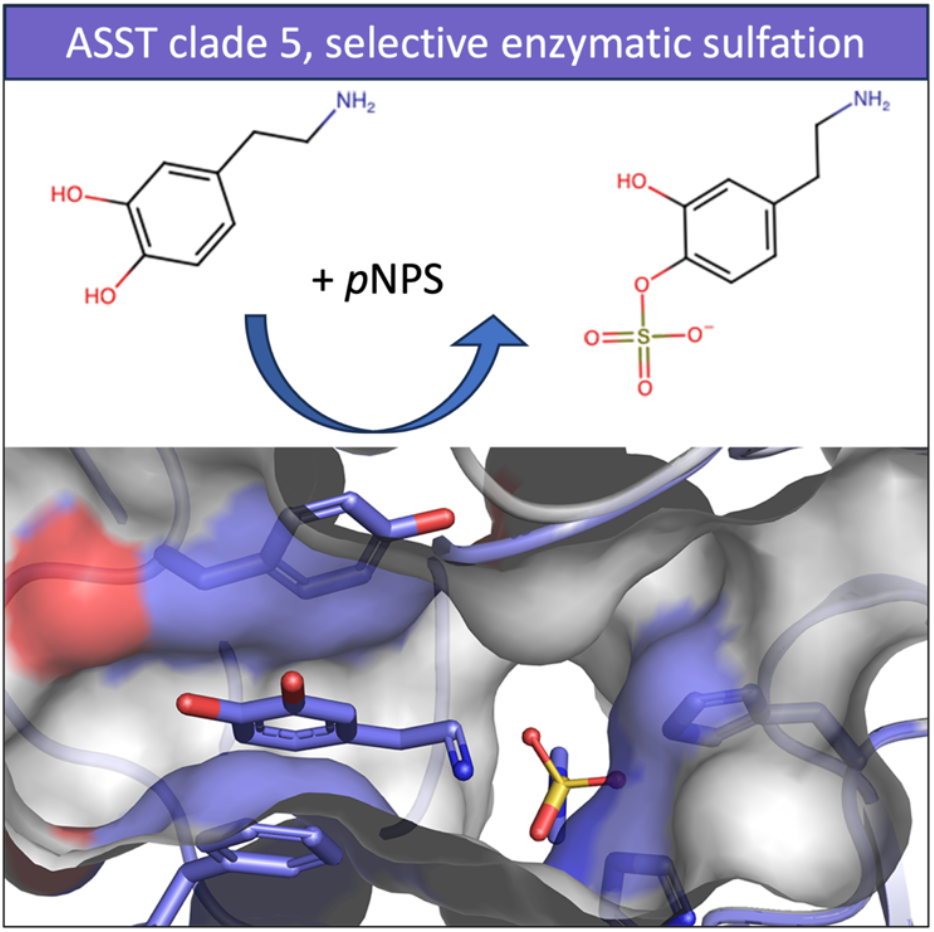

## INTRODUCTION

Sulfate plays a fundamental role in the structure and function of many essential biological molecules. In particular, it participates in the activation/inactivation and detoxification of many compounds (1), whether endogenous such as hormones (2) or exogenous such as xenobiotics (3–5). The transport of sulfate across the cell membrane is carried out by an active, energy-dependent mechanism and facilitated by proteins called sulfate permeases (6). Once assimilated, sulfate can be incorporated into essential cellular molecules through the action of sulfotransferases (7).

Sulfotransferases are a family of enzymes capable of transferring a sulfonate group to substrates with hydroxyl or amine groups. Initially discovered in 1980, when two sulfotransferases were isolated in eukaryotic cells, they were subsequently identified in prokaryotic organisms (8,9). Based on their sulfate donor specificity, sulfotransferases are classified into two major superfamilies: PAPS-dependent sulfotransferases and PAPS-independent sulfotransferases, also referred to as arylsulfate sulfotransferases (ASSTs) (10).

PAPS-dependent sulfotransferases are predominantly found in eukaryotic cells and are structurally characterized by a fold composed mainly of α-helices and few β-sheets (11), while ASSTs have an architecture consisting mainly of β-sheets, accompanied by a few α-helices (12) (Figure 1). Owing to their presence in humans, PAPS-dependent sulfotransferases enzymes have been extensively studied. In contrast, ASSTs remain poorly studied, mainly because they are primarily found in bacteria.

**Figure 1:**
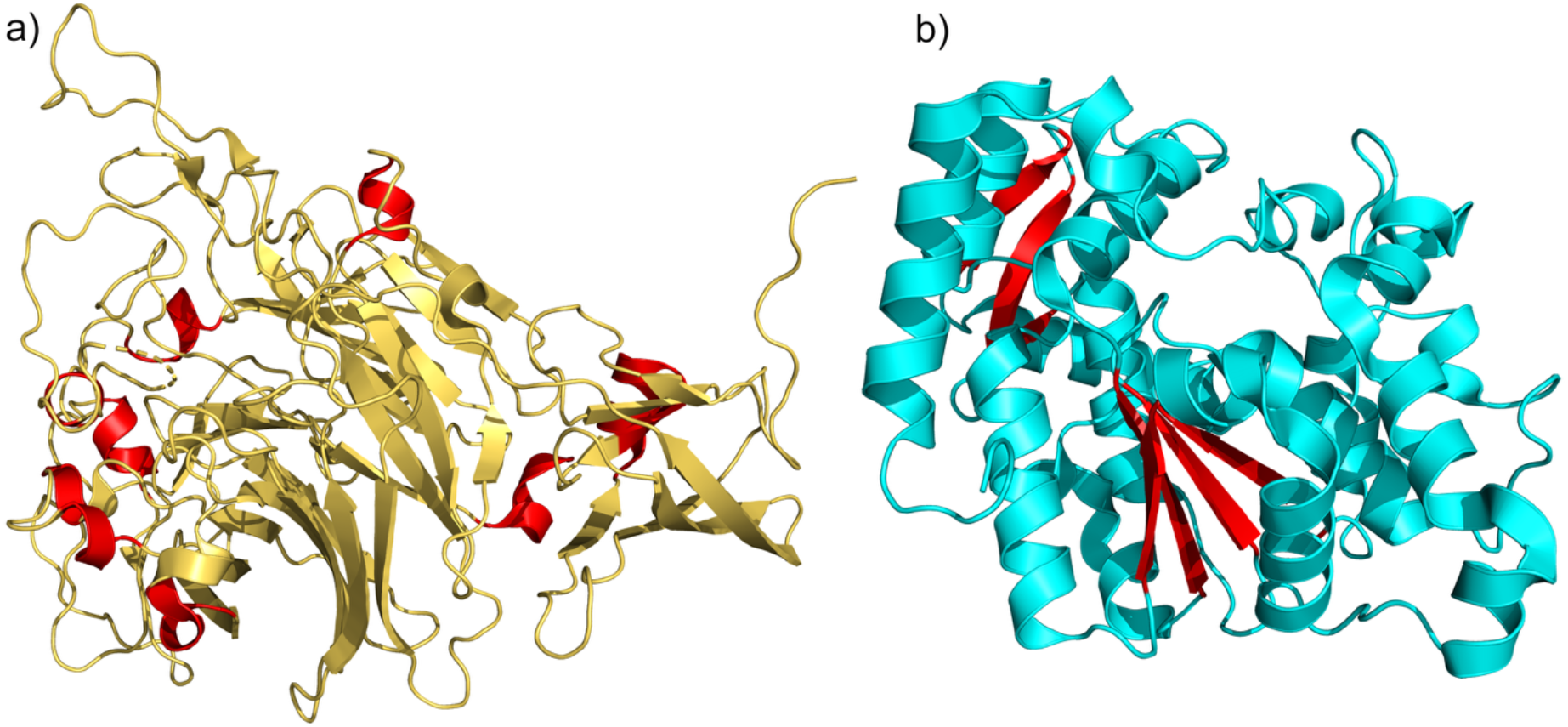
Overall structures of two sulfotransferases. (**a**) PAPS-independent arylsulfate sulfotransferase (ASST) from *Escherichia coli* (PDB ID: 3ELQ), shown in yellow, with α-helices highlighted in red. (**b**) PAPS-dependent sulfotransferase from *Arabidopsis thaliana* (PDB ID: 5MEX), shown in blue, with β-sheets highlighted in red. The Figure was drawn using PYMOL (13).

ASST catalyze the transfer of a sulfonate group from a sulfate donor distinct from PAPS to an acceptor, and in most cases both the donor and acceptor metabolites remain largely unknown. A catalytic mechanism was proposed following the resolution of the first crystallographic structure of an ASST from *Escherichia coli* (strain CFT073) (12). The proposed mechanism is of bi-bi ping-pong type, in which the sulfonate group, derived from a donor such as *p*-nitrophenyl sulfate (*p*NPS), is first transferred to the catalytic histidine His436 (N^ε2^), and subsequently transferred to an acceptor (12,14). The first step of catalysis was confirmed by observing, in the crystallographic structure, a sulfo-group on His436 in the presence of *p*NPS. Debate about the second step and the residues involved in the transfer to the acceptor is ongoing. The mechanism has been recently revised by Balci *et al*. (15) based on molecular docking combined with Quantum Mechanics/Molecular Mechanics (QM/MM) free energy calculations, proposing a mechanism involving two of the three conserved histidines.

Interest in the fundamental study of PAPS-independent arylsulfate sulfotransferases has grown in recent years, driven by the need to better understand the diversity and catalytic mechanisms of this large enzyme family (12,16–18,7,19,20). From a physiological point of view, arylsulfate sulfotransferases are attracting growing interest due to their presence in the human microbiome, particularly in the gut microbiota, where they could influence many aspects of host physiology. The gut microbiome plays a central role in human health, intervening in various processes such as digestion, immune regulation, and even behavior (20–23). In line with this, metabolite sulfation by the gut microbiome was exemplified by a study showing that *p*-cresol was sulfated by an ASST from a gut associated bacterium, becoming *p*-cresyl sulfate (PCS), and its increased levels are correlated with worse outcome of chronic kidney disease patients (24).

From a biotechnological perspective, enzymatic sulfation of metabolites offers an attractive alternative to chemical methods, which often lack regioselectivity and require harsh conditions (25). ASSTs could therefore provide valuable tools for the selective modification of bioactive compounds (19). To this end we need to understand the sequence-structure-function relationship.

In this study, we present complementary structural and biochemical analyses of an ASST from *Salmonella typhimurium*, further on named St_ASST_5, shedding light on an ASST from a so far uncharacterized clade, based on an extensive phylogeny provided by Barbeyron *et al*. (Barbeyron *et al.*, in preparation). The X-ray crystal structure reveals conserved architectural features shared with *E. coli* ASST, but also highlights specific features, unique to this clade of ASSTs. Combined with biochemical activity assays, QM/MM free energy simulations and site directed mutagenesis we investigated the catalytic mechanism of this enzyme with a focus on the understanding of substrate specificity. Together, these approaches provide new insights into the molecular basis of catalysis and substrate recognition within the ASST family.

## EXPERIMENTAL PROCEDURES

### Protein expression, production and purification

ASSTs from *Escherichia coli* (12) belonging to clade 1 (Ec_ASST_1), *Salmonella Typhimurium* from clade 5 (St_ASST_5) (26), and *Desulfitobacterium hafniense* from clade 6 (Dh_ASST_6) (16) were heterologously expressed in *E. coli*. The corresponding genes (Table S1) were cloned into pET15 and pET100/D-TOPO expression vectors. Protein production was carried out in 300 mL of auto-inducible ZYP-5052 medium supplemented with 100 µg/mL ampicillin. The cultures were grown at 20°C with 150 rpm for 72 hours. Cells were harvested by centrifugation at 5000×g, 4°C for 45 minutes. The supernatants were discarded and cell pellets were frozen at − 20°C before use. Pellets were resuspended in 20 mM Hepes pH 7.5, 300 mM NaCl and disrupted using ultrasonicator (Branson Sonifier SFX 250). The cell lysates were centrifuged at 20,000×g, 4°C and for 45 minutes. The soluble fractions were then purified by affinity chromatography (ÄKTA™ chromatography system) using a 5 mL Histrap FF (Cytiva) column loaded with Ni^2+^ and equilibrated with Hepes buffer 20 mM pH 7.5 containing 300 mM NaCl and 20 mM imidazole. The recombinant ASSTs were eluted with the same buffer containing imidazole (500 mM). The eluted fractions were then analyzed on 12% SDS-PAGE gel. The fractions containing the protein were pooled together and then concentrated using a 30 kDa Amicon to a final volume of 5 mL. This was followed by size exclusion chromatography using a Hiload 16/600 Superdex 200pg column (Cytiva), equilibrated with MES buffer 20 mM pH 6.5 containing 200 mM NaCl. The fractions were then analyzed on a 12% SDS-PAGE gel (Figure S1) and the purified protein was concentrated using a 30 kDa Amicon to a concentration greater than 10 mg·mL^-1^ and then stored at 4°C. The experimental molecular mass of ST_ASST_5, determined by UPLC-HRMS (Figure S2), was consistent with the theoretical mass. The concentration was calculated from the absorption at A280 and the extinction coefficient calculated using the ProtParam tool from ExPASy (“0.1%” of 1.462 for Ec_ASST_1, 1.612 for Dh_ASST_6 and 1.41 for St_ASST_5). The final concentrations of Ec_ASST_1, Dh_ASST_6 and St_ASST_5 were 2.3, 16.9 and 14.5 mg·mL^-1^, respectively.

#### Determination of Optimal pH and Temperature

The activity of the purified St_ASST_5 was determined in triplicates in 96-well microtiter plates, using a modified protocol based on Herlet *et al*. (27), at various pH values ranging from 5 to 10, obtained by adjusting the pH of a universal Britton-Robinson buffer (50 mM), and at various temperatures ranging from 30 to 60°C (Figure S3). Each well contained 3 µL of purified enzyme (2 mg·mL^-1^ in MES buffer 20 mM, pH 6.5, 200 mM NaCl) and 67 µL of reaction mixture containing phenol and *p*NPS at final concentrations of 20 mM and 10mM respectively. After 15 min incubation in a PCR thermal cycler, 140 µL of a 1M Na_2_CO_3_ solution were added to stop the reactions and raise the pH above 10. Subsequently, 100 µL of each reaction were transferred to flat-bottom 96-well plates (Greiner Bio-One, cat. No. 655101), and absorbance at 405 nm was measured using a microplate reader (TECAN Infinite M1000, Männedorf, Switzerland). The activity was determined by comparing absorbance to a *p*NP standard curve obtained under the same conditions, with increasing concentration of *p*NP (0.0 to 0.25 mM). Data were processed using Origin (OriginLab) to generate contour plots of enzyme activity as a function of temperature and pH.

Thermal stability of St_ASST_5 was additionally assessed by nano differential scanning fluorimetry (nanoDSF; Prometheus NT.48). Protein samples were loaded into nanoDSF capillaries and subjected to a controlled temperature ramp from 20°C to 90°C at a rate of 1°C/min. Intrinsic fluorescence emission ratios (F_350_/F_330_) were continuously recorded to determine the apparent melting temperature (T_m_). Simultaneously, static light scattering was monitored to detect protein aggregation during unfolding. Experiments were conducted in a buffer composed of 100 mM Tris-HCl and 50 mM NaCl (pH 8.0), using three protein concentrations (1.0, 0.5 and 0.1 mg·mL^-1^ to evaluate potential concentration-dependent effects on thermal stability (Figure S4a).

#### Thermal Stability Assay

The thermal stability of St_ASST_5 was evaluated by monitoring residual catalytic activity following controlled thermal pre-incubation. Purified St_ASST_5 enzyme (35.9 nM) was incubated at 58°C in 50 mM MES buffer (pH 5.5) in the absence of substrates for defined time intervals (0-20 min). After each pre-incubation period, the enzymatic reaction was initiated by the addition of a pre-equilibrated substrate solution containing 20 mM phenol and 10 mM *p*NPS in the same buffer, and allowed to proceed for 5 min at 58°C. The reactions were terminated by the addition of 1 M Na₂CO₃, and residual activity quantified spectrophotometrically as described before. Residual activities were normalized to a non-heated control, defined as 100 % activity (Figure S4b).

#### Substrate screening assay

ASST activity was determined by measuring the initial rate of *p*NP release from *p*NPS by spectrophotometry at 405 nm for 30 min (Tecan Infinite M1000). All reactions were performed in triplicate in 96 well microplates (Greiner Bio-One, Kremsmünster, Austria; cat. no. 655101) at 37 °C in 100 mM Tris–HCl buffer, pH 8.0, in a final reaction volume of 200 µL containing *p*NPS (10 mM), acceptor substrate (20 mM) and purified enzyme (final concentrations: 0.025 mg·mL⁻¹ for St_ASST_5, 0.02 mg·mL⁻¹ for Ec_ASST_1 and 0.0013 mg·mL^-1^ Dh_ASST_6). The following acceptor substrates, phenol, resorcinol (Merck, Darmstadt, Germany), and dopamine (TCI Chemicals, Tokyo, Japan) were dissolved in Tris-HCl buffer, whilst less soluble acceptor substrates including tyramine (Merck, Darmstadt, Germany) and epicatechin (TCI Chemicals, Tokyo, Japan) were used in Tris-HCl buffer containing 10% DMSO (v/v). Control experiments without acceptor substrate or without enzyme were conducted to monitor the formation of *p*NP resulting from enzyme catalyzed hydrolysis or chemical hydrolysis of *p*NPS respectively. This background hydrolysis was systematically subtracted from the activities towards the different acceptors. Protein purity was assessed using Agilent microfluidic protein analysis chips (Agilent technologies, Santa Clara, CA, USA cat. no. 5067-1517). Purity was taken into account in the enzyme activity calculations.

#### Reaction product characterization by LC-MS

Products formation was analyzed by liquid chromatography coupled to tandem mass spectrometry (LC-MS/MS). For the sulfation by St_ASST_5, dopamine was added to a final concentration of 1.6 mM in 100 mM Tris-HCl buffer pH 8.0 containing 1.2 mM *p*NPS and 0.025 µg protein. The reactions were stopped after 24 hours by the addition of 500 µL of methanol before LC-MS analyses were performed using a Vanquish Flex UHPLC system coupled to an Exploris 240 Orbitrap mass spectrometer (Thermo Scientific). Separation was achieved on a Hypersil Gold Aq C18 column (150 × 2.1 mm, 2.6 µm) at 30°C with a flow rate of 0.5 mL/min, using a water/acetonitrile gradient (0.1% formic acid) over 9 min. The injection volume was 1 µL and the autosampler was maintained at 10°C.

Mass spectrometry was carried out using a heated electrospray ionization (H-ESI) source. Full MS scans were acquired in positive and negative modes at a resolution of 120,000 over a mass range of m/z 67–1000. Data-dependent MS/MS analyses were performed in negative mode at a resolution of 60,000, using targeted inclusion lists and HCD fragmentation (10, 35, and 60% collision energies). Additional collision energies were tested without conclusive results. Data acquisition was performed using Xcalibur 4.7 (28), and data were processed using FreeStyle, MZmine (29), and Microsoft Excel.

#### NMR Analysis of reaction product

One-dimensional (^1^H) and two-dimensional (^1^H COSY) NMR spectra (Figure S5) were recorded on a BRUKER AVANCE III HD 500 spectrometer equipped with a 5 mm BBFO indirect detection probe. Samples were analyzed at 25°C in D₂O (10 mg·mL⁻¹). Sulfated products were obtained by subjecting the samples to the same reaction conditions as those used for the spectrophotometric assays, with an extended incubation time of 24 h. Spectra were acquired using standard BRUKER pulse programs with a relaxation delay of 2 s, a 30° pulse, and 64 scans. Chemical shifts were referenced to TMS as an external standard and are reported in ppm (30,31).

#### Steady-State Kinetic Assays of St_ASST_5

To determine the kinetic mechanism, inhibition type, and catalytic constants of St_ASST_5, steady-state enzymatic assays were performed at 37°C in 50 mM MES buffer (pH 5.5). Reactions were carried out in 96 well microplates, with each well containing a final volume of 40 µL. The reaction mixture comprised varying concentrations of *p*-nitrophenyl sulfate (1-10 µM) at fixed phenol concentrations (8-180 mM), and vice versa, to generate a matrix of substrate combinations. Each reaction was initiated by the addition of 10 µL of purified St_ASST_5 enzyme at a final concentration of 35.9 nM to wells pre-equilibrated with substrate solutions. After a 7 min incubation at 37°C, previously confirmed as the linear phase of product formation, the enzymatic reaction was quenched by the addition of 140 µL of 1 M Na₂CO₃ allowing spectrophotometric detection at 405 nm. Initial velocities were determined from the absorbance values and plotted as a function of substrate concentration. All assays were performed in triplicate, and reported values represent the mean of independent experiments. Double-reciprocal (Lineweaver-Burk) plots and secondary plots were constructed to determine the enzyme’s kinetic mechanism and inhibition type. A global nonlinear regression analysis (GraphPad Prism 8.0.1) was used to fit the data to appropriate kinetic models (e.g., ping-pong, or sequential), yielding true kinetic constants (*K*_m_, V_max_, *k*_cat_) and, where applicable, inhibition constants (*K*_i_).

#### Crystal structure determination

Crystallization trials were carried out by sitting drop vapor diffusion using a Crystal Gryphon multi-distribution nanodrop robot (Art Robbins Instruments) using commercial screens: PACT Premier Eco Screen, PEGsuite, PEGsuite II and JCSG-plus Eco Screen. The initial crystallization conditions of St_ASST_5 was manually optimized and single crystals were obtained as follows: 2 µL of enzyme at 14.5 mg·mL^-1^ were mixed with 2 µL of reservoir solution containing 1.4 M Ammonium Sulfate and 1 M Na acetate pH 4.6 in hanging drops at 20°C. Single crystals of St_ASST_5 in complex with *p*NPS (St_ASST_5_*p*NPS) were obtained by soaking pre-formed native crystals with substrate powder directly in the crystallization drop. For the St_ASST_5 complex with dopamine (St_ASST_5_dopamine), the crystals used were formed from a 1:1 mixture of the enzyme and *p*NPS. This was followed by a desalting step using PD10 columns (Cytiva) to remove excess of *p*NPS and *p*NP product. New crystals formed by this protein solution were subsequently soaked with dopamine powder. Crystals were flash-cooled in liquid nitrogen using a cryoprotectant of 20% of glycerol or ethylene glycol.

Diffraction data for the St_ASST_5 native and in complex with *p*NPS were collected on the PROXIMA2 and St_ASST_5_dopamine was collected on PROXIMA1 beamline at SOLEIL synchrotron at Gif-sur-Yvette (France). The data were processed using XDS (32) and scaled with Aimless 440 from the CCP4 program package (33). The structures were solved by molecular replacement with the CCP4 suite program Phaser (34) using an AlphaFold model of St_ASST_5 as the starting model (UniParc: UPI00000CCE18). For all the structures, the initial molecular replacement solutions were further refined with the CCP4 suite program REFMAC5 (35). Iterative rounds of model building and refinement were carried out using COOT (36). Water molecules were automatically added with WARP implemented in REFMAC5 and verified manually. The validation of the crystal structures was performed with MolProbity (37).

#### Site-directed Mutagenesis

Site-directed mutagenesis of St_ASST_5 was performed to investigate the roles of residues involved in substrate binding and catalysis. Residues Phe174 and Tyr557, identified as potentially involved in substrate binding, were individually replaced by alanine (F174A and Y557A), and a double mutant (F174A/Y557A) was also generated. In addition, the catalytic residue His401 was substituted by glutamine (H401Q) or asparagine (H401N) to assess its role in catalysis. Mutagenesis was carried out using the QuikChange Site-Directed Mutagenesis Kit for single mutations and the QuikChange Multi Site-Directed Mutagenesis Kit (200518 and 200515, Agilent Technologies) for double mutations, using the primers listed in Table S2. All mutated constructs were verified by complete gene sequencing (Eurofins Genomics, France). Expression and purification of the mutant proteins were performed as described in the “Protein expression, production and purification” section.

#### Small angle X-ray scattering (SAXS) analyses

SAXS measurements were conducted at the French Synchrotron SOLEIL (St. Aubin, France) on the SWING beamline and performed at 15°C. The SAXS data were measured using the same buffer as for crystallization trials (MES buffer 20 mM, pH 6.5, NaCl 200 mM). The scattering vector is defined as q = 4 π/λ sinθ, where 2θ is the scattering angle. Data were collected covering a q range of 0.005-0.5 Å^-1^. Data were recorded using an AVIEX170170 CCD detector at a distance of 1.807 m (λ = 1.033 Å). The protein solution of St_ASST_5 was prepared at final concentrations as given in Table 2. SAXS data were collected on samples eluting from an online size exclusion high-performance liquid chromatography (SEHPLCBio-SEC3Agilent) column, directly connected to the SAXS measuring cell. A volume of 60 µl of protein sample at 14.5 mg·mL^-1^ was injected and eluted directly into the SAXS capillary at a flow rate of 0.2 mL·min^-1^. The elution buffer consisted of 20 mM MES buffer at pH 6.5 and 200 mM NaCl, identical to the one used for the size exclusion chromatography purification step. Two hundred fifty SAXS frames were collected continuously during the elution at a frame duration of 1.5 s and a dead time between frames of 0.5 s. One hundred frames accounting for buffer scattering were collected before the void volume. SAXS data were normalized to the intensity of the incident beam and background (i.e. the elution buffer) subtracted using the program FoxTrot (38). The experimental curves were analyzed using Primus as part of the ATSAS suite (39) to calculate the Rg value in the Guinier region, and GNOM (40) to calculate the P(r) and D_max_ values. The experimental curve was compared with the theoretical diffusion curve of the crystal structure using CRYSOL as implemented in the ATSAS suite (39). DAMMIF (41) was used in subsequent steps for data interpretation and *ab inito* modelling to find the best shapes that fit the experimental data. The quality of the 3D modeling was determined using the discrepancy χ^2^, defined according to (42). Values lying in the range of 0.9−1.1 are accepted to indicate a good fit between the models and the data. However, the calculation of χ^2^ is inversely proportional to the measurement error. Using the low-error detector at SOLEIL, higher χ^2^ values were obtained (43,44).

#### Analyses of the genomic context

Comparative genomics was performed using the tools available on the Genoscope’s MaGe (Magnifying Genomes) website (45). A window of 20,000 base pairs around the ASST genes was set up to identify any clusters (Figure S6). The genes in the clusters identified around ASSTs were listed and analyzed by multiple alignment using the NCBI website, which has a database of manually annotated protein sequences, UniProt (46). For unannotated sequences, an alignment was performed, again on NCBI, on nr (non-redundant), a database of automatically annotated protein sequences. Functional analyses were then performed using the CDD Search (Conserved Domain Database) and InterProScan tools to predict conserved protein domains and potential catalytic sites (47,48). Finally, structural prediction was performed by AlphaFold2 (49) and the structures obtained were compared to functionally similar enzymes present in the Protein Data Bank (PDB) (50) using DALI (51).

#### Molecular Docking

Molecular docking calculations were carried out using the Molecular Operating Environment (MOE) v2024.06 (52). The crystal structure of St_ASST_5_*p*NPS, in which the catalytic His401 residue is present in its sulfonated state, was used as the receptor structure. Prior to docking, all ligands, solvent molecules, and non-relevant small molecules were removed from the structure. Initial protonation states were assigned using the H++server (53). The protonation states of the three conserved active-site histidine residues (His242, His317, His401) were adjusted according to the constant pH molecular dynamics calculations reports for the homologous ASST system in Balci *et al.* (15).

Phenol and dopamine were docked separately into the active site of St_ASST_5 using the same defines docking setup. The docking pocket was defined around the sulfonated catalytic His401 residue and selected active site residues, including His317, Arg335, His242, Phe562, Asn319, Cys472, and Val473. In both cases, the receptor was kept rigid, while ligand flexibility was treated by rotation around rotatable bonds. Docking was performed using the Triangle Matcher placement method with London dG scoring, followed by rigid-receptor refinement and rescoring using GBVI/WSA dG (53,54). For each substrate an independent docking calculation was carried out with a maximum of 3000 generated poses, using the same active-site definition, receptor preparation, placement algorithm, scoring function, refinement procedure, and pose-generation settings.

The resulting poses were evaluated based on their docking scores and mechanism-based geometric criteria derived from previously proposed ASST sulfo-transfer mechanism (15). Catalytically plausible poses were required to orient the acceptor hydroxyl group toward the sulfur atom of the His-401-bound sulfo-group and to place the group in proximity to the conserved histidine reside involved in acceptor activation and proton transfer, namely His-317. For dopamine, the poses in which one of the catechol hydroxl groups fulfilled these criteria were considered productive. The final poses were selected based on a combination of favorable docking score, productive acceptor-group orientation, and compatibility with the proton coupled sulfo-transfer mechanism. The selected docking poses were used as starting structures for subsequent tunnel analysis and molecular dynamics simulation.

#### Tunnel Analysis

Tunnel analysis was performed for the separately for the selected docking-derived St_ASST_5-phenol and St_ASST_5-dopamine complexes using the CAVER v3.0 (55) plugin in PyMOL (13). Tunnel detection was initiated from the active-site cavity containing the sulfonated catalytic His401 residue, corresponding to the covalent sulfo-enzyme intermediate and the acceptor-binding region identified by docking. In CAVER, the starting point was defined by coordinates located in this active-site region. The detected pathways were therefore interpreted as putative substrate-access and product-release tunnel connecting the catalytic pocket with the protein surface.

A probe radius of 0.6 Å was used for tunnel detection. The shell radius and shell depth were set to 3.0 and 4.0 Å, respectively. Tunnels were clustered using average-link hierarchical clustering with a clustering threshold of 3.5, and the lowest-cost tunnel per cluster was retained for further analysis. Residues located within 3.0 Å of the identified tunnels were defined as tunnel-lining residues and were used to characterize the chemical environment of each pathway. For each detected tunnel cluster, the throughput, tunnel length, curvature, and bottleneck radius were extracted. These parameters were used to compare the geometry and accessibility of the active-site tunnels in the phenol- and dopamine-docked St_ASST_5 models.

#### Molecular dynamics (MD) simulations

Classical molecular dynamics simulations were initiated from the selected docking-derived St_ASST_5-phenol and St_ASST_6-dopamine complexes based on the St_ASST_5_pNPS crystal structure. In this structure, the catalytic His401 residue is present as a sulfonated histidine and therefore represents the covalent sulfo-enzyme intermediate.

The protonation states of the catalytic histidine residues (His242, His317, His401) were assigned based on previous constant-pH simulations of the homologous enzyme Dh_ASST_6 (15). In the St_ASST_5 model, the non-sulfonated nitrogen of His401 was protonated at the Nδ position, His242 was protonated at the Nε position, and His317 was modeled in the singly protonated Nδ state. Although the corresponding histidine was predicted to be doubly protonated in the absence of acceptor substrate in the Dh_ASST_6 constant pH simulation (15), the singly protonated Nδ state was chosen here to represent the reactant configuration of the second catalytic step, in which His317 acts as the acceptor-activating base. The protonation states of all remaining titratable residues were determined using the H++ server at pH 6.5 (56).

All classical molecular dynamics simulations were performed with Amber22 (57)(Case *et al.*, 2023) using the ff19SB (58)(Tian *et al.*, 2020) force field for the protein. The acceptor substrates phenol and dopamine, as well as the sulfated His401 residue were described using AMBER/GAFF atom types. Missing partial charged for the non-standard atoms of the acceptor ligand and the sulfonated His401 residue, including the sulfo-group and atoms involved in the His401-sulfo linkage, were obtained from quantum-mechanical (QM) optimizations at the B3LYP/6-31+G* (59–63) level using Gaussian16 (64) and were parametrized using a RESP (65) model (RHF/6-31G**) (59,60,66) on the optimized geometry with Antechamber (67). The bond length between the His401 nitrogen atom and the sulfur atom of the sulfo group was set to 2.2 Å based on previous observations for the homologous Dh_ASST_6 (15).

The systems were solvated under periodic boundary conditions in cubic simulation boxes using OPC (68) water molecules and neutralized by adding Na⁺ counterions. Each system was subjected to a three-step energy minimization protocol, in which first hydrogen atoms, then solvent molecules, and finally the complete system were relaxed. The systems were subsequently heated in two stages under harmonic restraints applied to the protein backbone. In the first step, the temperature was increased from 0 to 200 K, followed by a second heating step from 200 to 300 K over 1 ns using the Langevin thermostat in the NVT ensemble. Equilibration was carried out in six consecutive 20 ps steps while gradually reducing the applied restraints. The first four equilibration steps were performed in the NVT ensemble, whereas the final two steps were carried out in the NPT ensemble without restraints on the protein backbone atoms.

For production sampling three simulations per system were performed for 200 ns using a 2 fs integration time step. Periodic boundary conditions were applied throughout, and a cutoff of 10 Å was used for non-bonded interactions. Bonds involving hydrogen atoms in water molecules were constrained using the SHAKE (69) algorithm. Trajectory analysis was performed using cpptraj (70) and the Python package MDAnalysis (71). Molecular visualizations were prepared using PyMOL (13).

The complete simulation protocol is provided in Supplemental Information section 7 and is not discussed further in the main text.

#### QM/MM Umbrella Sampling

Quantum Mechanics/Molecular Mechanics (QM/MM) umbrella sampling simulations were performed to investigate the second catalytic sulfo-transfer step for phenol and for both catechol hydroxyl positions of dopamine. The reaction was described by proton transfer from the acceptor hydroxyl group to His317 and formation of the acceptor O-S bond to the sulfo-group bound to His401. Starting structures for the QM/MM umbrella sampling were selected from the equilibrated classical MD trajectories based on near-attack conformation criteria derived from the previously proposed sulfo-transfer mechanism of the homologous enzyme Dh_ASST_6 reported by Balci *et al.* (15).

Two distance-based collective variables (CVs) were used. CV1 describes the distance between the transferring hydroxyl hydrogen and the catalytic nitrogen of His317, whereas CV2 describes formation of the bond between the acceptor oxygen and the sulfur atom of the His401-bound sulfo-group. For phenol, the H-N distance was sampled from 1.8266 to 1.0000 Å and the O-S distance from 3.6647 to 1.6000 Å. For the favored dopamine OH position, the corresponding ranges were 1.8483 to 1.1000 Å and 3.8528 to 1.6000 Å. For the alternative dopamine OH position, the ranges were 1.9660 to 1.1000 Å and 4.9860 to 1.6000 Å.

Umbrella windows were generated by linearly interpolating both CVs between reactant and product configurations. Phenol was sampled with 25 windows. The favored dopamine pathway initially used the same windowing strategy, with three additional windows introduced in the transition-state region to improve neighboring-window overlap, resulting in 28 windows. For the alternative dopamine pathway, one additional refinement window was introduced, resulting in 26 windows. Harmonic restraints of 100 kcal mol−1 Å−2 were applied to both CVs, consistent with the restraint files used in the umbrella calculations.

All QM/MM umbrella sampling simulations were carried out in Amber22 (57) using the DFTB3 (72) semiempirical QM method. The QM region was chosen based on the active-site architecture of St_ASST_5. The QM region included all atoms directly involved in the chemical transformation, namely the acceptor substrate, the sulfonated His401 residue, and His317, which acts as catalytic base in the second step. In addition, Arg335, His242, Phe562, Asn319, Cys472, and Val473 were included because they form the immediate catalytic pocket and contribute to positioning and stabilization of the sulfo-group and acceptor substrate. This QM region therefore captures proton transfer, O-S bond formation, sulfo-group stabilization, and the local active-site environment required for the reaction.

Covalent boundaries between the QM and MM regions were treated using hydrogen link atoms (73). Amino-acid residues were truncated at the Cα–Cβ bond, whereas the acceptor substrate moieties were retained intact. A total QM charge of +1 was assigned, and a QM cutoff of 10 Å was used. Simulations were performed at 300 K using a Langevin thermostat under periodic boundary conditions. A non-bonded cutoff of 10 Å was applied. Harmonic restraints were applied to both CVs in each umbrella window using a force constant of 600 kcal·mol⁻¹·Å⁻² for both phenol and dopamine. Each umbrella window was simulated for 10000 steps with a time step of 0.0002 ps, corresponding to 2 ps per window. The windows were simulated sequentially, with the final restart structure of each window used as the starting structure for the following window.

Umbrella trajectories were analyzed using cpptraj (70). One- and two-dimensional potential-of-mean-force profiles were reconstructed using WHAM (74) and pyMBAR-based analysis (75). The favored and alternative dopamine profiles were analyzed independently and each was referenced to its own reactant basin at 0 kcal mol−1. Consequently, comparisons of activation barriers are direct within the sampled pathways, whereas product-state offsets should be interpreted as pathway-referenced reaction free energies rather than absolute binding free-energy differences between dopamine orientations.

## RESULTS and DISCUSSION

### Phylogenetic selection of genes

Phylogenetic analyses (Barbeyron *et al.,* in preparation), when applying a cut-off of 35% sequence identity, divide the sequences of ASSTs present in public databases into 22 clades. Despite this common cut-off, some of these clades contain many members (i.e. clade 1 includes 652 sequences, and clade 2 1099 sequences) while other are rather small (i.e. clade 11 has 5 sequences or clade 9 only 3 sequences). In the aim to examine whether the separation into different clades relates to differences in functional aspects, we have selected ASSTs from different clades for further investigation. In the present study, we compare two representative and characterized ASSTs, Ec_ASST_1 from *Escherichia coli* (clade 1) (12) and Dh_ASST_6 from *Desulfitobacterium hafniense* (clade 6) (15,16) to a third one, focus of the here described biochemical and structural characterization, the St_ASST_5 from *Salmonella typhimurium* (clade 5).

### Selection of Catalytic Conditions Balancing Activity and Stability

Optimal catalytic conditions have been reported earlier for Ec_ASST_1 (12) and Dh_ASST_6 (16) but not for St_ASST_5, we have thus screened activity as a function of pH and temperature for the latter. The search for optimal pH and temperature revealed an apparent optimum for St_ASST_5 at pH 5 and 58 °C (Figure S3). NanoDSF measurements, however, showed that this temperature exceeds the thermal melting point of the enzyme (Tₘ = 54.11 °C; Figure S4a), indicating limited structural stability under these conditions. Consistent with this observation, incubation at 58 °C resulted in rapid time-dependent inactivation, with approximately 50% loss of enzymatic activity after 5 min and 100% after 15min (Figure S4b). These findings demonstrate that the temperature optimum derived from activity profiling reflects short-term catalytic performance rather than sustained enzyme stability. To prevent time-dependent inactivation during kinetic measurements, subsequent experiments were conducted at 37 °C and pH 5.5, conditions that preserved catalytic activity over the activity assay timeframe.

### Screening of activity against a selection of substrates

Phenolic compounds are known to be good acceptor substrates for ASSTs. Consequently, we selected phenol, tyramine, epicatechin, dopamine, and resorcinol as acceptors to assess the sulfation abilities of the recombinant ASSTs. Most acceptors were chosen on their prior use in studies involving ASST enzymes. *p*NPS was selected as the sulfate donor, as it is commonly employed for this purpose in the literature (14,17,76). The experiment carried out first was a spectrophotometric monitoring which is based on the property of the desulfated donor, *p*NP, to absorb at 405 nm in basic conditions. The reactions were conducted at a constant temperature of 37°C and pH 8. The reactions details are given in Experimental procedures. In our study, we chose to compare three ASSTs, which are Ec_ASST_1, St_ASST_5 and Dh_ASST_6. For both Ec_ASST_1 and Dh_ASST_6, which are the most studied ASSTs to date, earlier studies have reported activities on various substrates (14,16,76), but to be able to directly compare activities with the enzyme St_ASST_5, focus of this study, it is important to perform the experiments in parallel and under same and comparable conditions.

The sulfotransferase activity strictly depends on the presence of an ASST and an acceptor (Figure 2a, b). However, in all cases without acceptor, a low hydrolysis activity with formation of *p*NP was observed, and for that reason, hydrolysis was subtracted from all results. The comparison allows us to conclude that the ASSTs from different clades display high but variable activity on different substrates. Notably, St_ASST_5 exhibits a clear preference for substrates bearing side chains, as observed with dopamine compared to resorcinol (Figure 2a). Although tyramine also contains a side chain, its reduced activity might be attributed to the absence of a second hydroxyl group on the aromatic ring, which possibly limits hydrogen bonding in the active site. But we cannot exclude that differences in activity are due to DMSO needed to solubilize this compound. In contrast, Ec_ASST_1 and Dh_ASST_6 display higher activity toward substrates with short or no side chains, such as resorcinol, and lower number of hydroxyl groups, consequently phenol or tyramine are preferred over dopamine (Figure 2). This suggests that Ec_ASST_1 and Dh_ASST_6 preferentially accommodate smaller substrates within their active site. In the following set of experiments, we further investigated reaction kinetics and specificity on the previously unreported arylsulfate sulfotransferase, focusing on St_ASST_5, to evaluate its enzymatic activity using phenol or dopamine as acceptor substrate, respectively.

**Figure 2.**
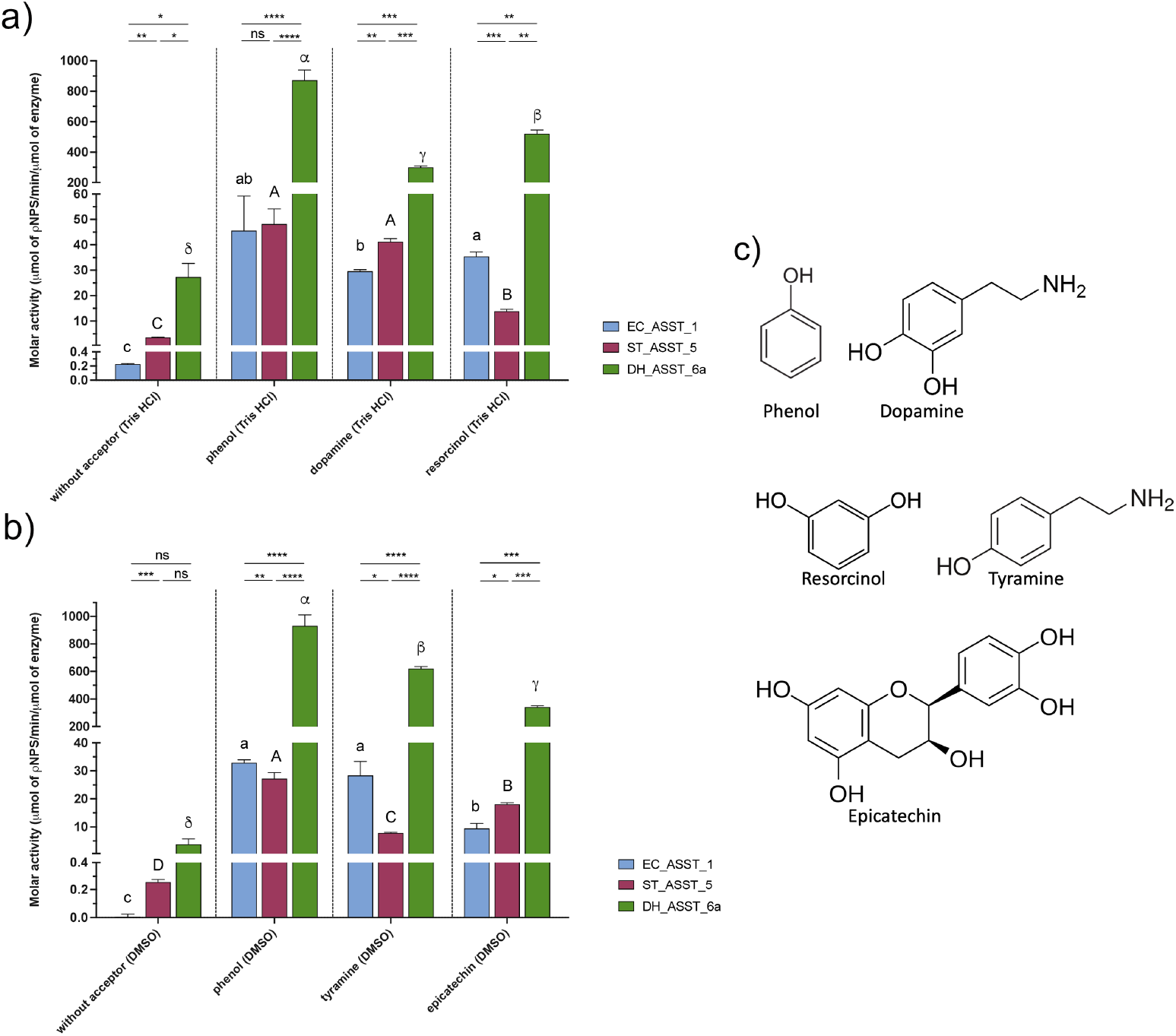
Molar activity of enzymes Ec_ASST_1, St_ASST_5, and Dh_ASST_6 in the presence of different acceptor substrates. The blue, pink, and green bars represent the enzymatic activity of Ec_ASST_1, St_ASST_5, and Dh_ASST_6, respectively. Enzymatic activity is expressed as μmol of *p*NPS·min⁻^1^·μmol⁻^1^ of enzyme. The “without acceptor” conditions correspond to activity measured in the absence of an acceptor substrate. Error bars indicate the standard deviation (SD) of independent replicates. Statistical analysis: a one-way Welch ANOVA test, followed by Dunnett T3 multiple comparison test, was performed for comparison of the three enzymes (Ec_ASST_1, St-ASST_5, and Dh_ASST_6) for the same substrate: (significance thresholds are indicated as follows: p < 0.0332 (*); p < 0.0021 (**); p < 0.0002 (***); p < 0.0001 (****)); a one-way Welch ANOVA test, followed by Dunnett T3 multiple comparison test with compact letter display, was conducted for comparison of different substrates for the same enzyme (bars that share at least one letter are not significantly different). (**a)** Acceptor substrates tested in Tris-HCl buffer: phenol, dopamine, and resorcinol. (**b)** Acceptor substrates tested in DMSO (10% v/v)/Tris-HCl buffer: phenol, tyramine, and epicatechin. (**c**) Chemical structures of the acceptor molecules used in the assay.

### Steady-State Kinetic Analysis Reveals a Ping–Pong Mechanism with Phenol-Dependent Inhibition

Steady-state kinetic experiments were conducted to elucidate the catalytic mechanism of St_ASST_5 (Figure 3). ASSTs are bireactant enzymes with two substrates and two products. To evaluate the kinetic behavior of the enzyme, a series of experiments were carried out varying the concentration of one substrate whilst keeping the other constant, with concentrations ranging from low to high concentrations for *p*NPS (50, 70, 100, 143 and 200 µM) and phenol (8, 10, 23, 32, 45, 70 and 180 mM). Reaction velocities increased with increasing phenol concentration up to approximately 45 mM, after which further increases in phenol led to a decline in activity, yielding a bell-shaped dependence characteristic of substrate inhibition (Figure 3a). Double-reciprocal (Lineweaver-Burk) plots of 1/v versus 1/[*p*NPS] resulted in a series of parallel lines, which is characteristic of a ping-pong (double-displacement) catalytic mechanism (Figure 3b) (77). Double-reciprocal plots of 1/v versus 1/[phenol] at different concentrations of *p*NPS (50, 70, 100, 143 and 200 µM) resulted in parallel lines, but only at non inhibiting phenol concentrations (up to approximately 45 mM) (Figure 3c). At higher phenol levels, the curves deviated and approached the y-axis asymptotically, further supporting the presence of phenol-dependent substrate inhibition (78).

**Figure 3.**
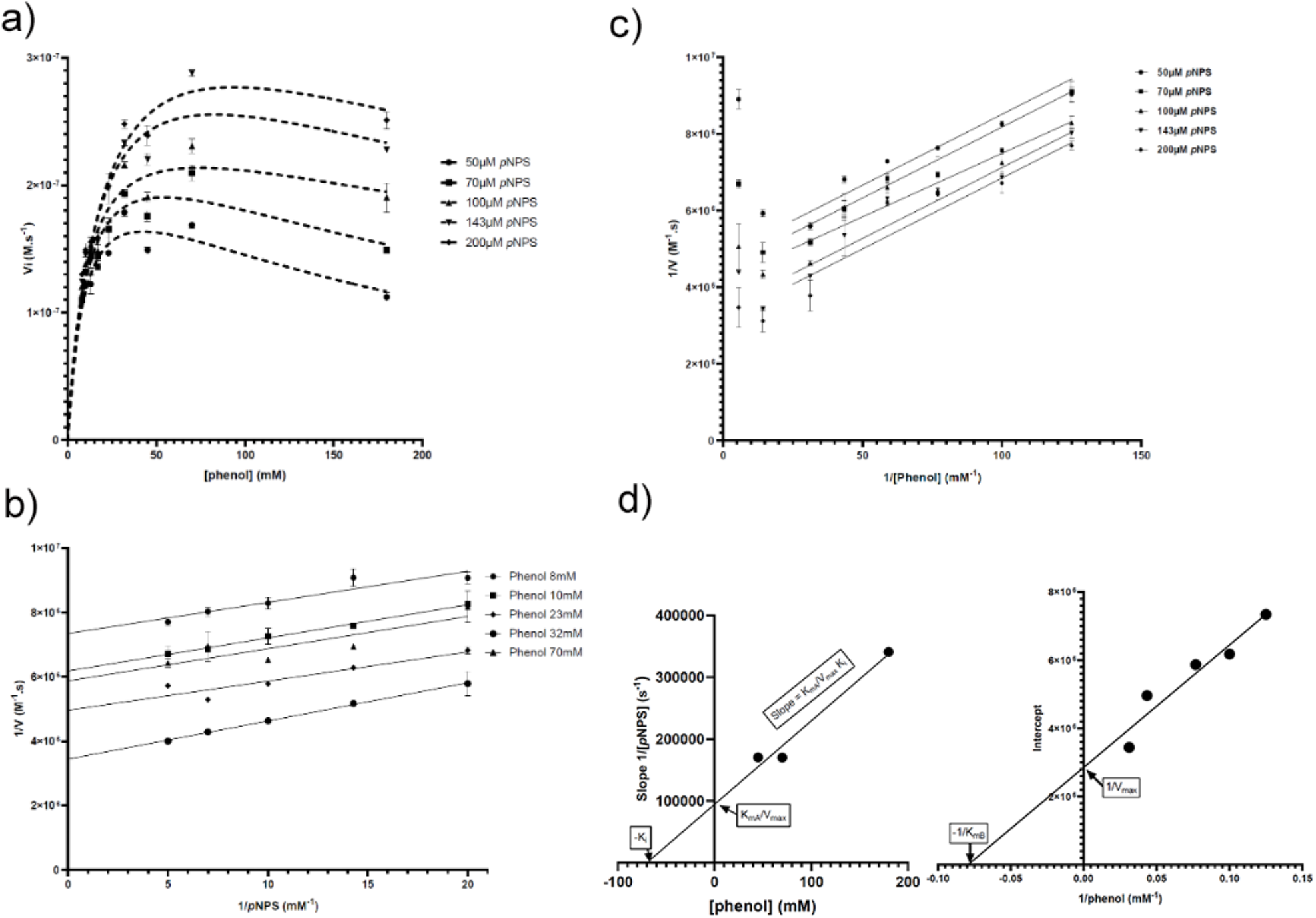
Kinetic analysis of St_ASST_5 catalyzed sulfation of phenol by *p*NPS. **a**) Initial reaction rates versus phenol concentration at different concentrations of *p*NPS. **b**) Same data set in double-reciprocal (Lineweaver-Burk) form using the same code of symbols. **c**) Double-reciprocal (Lineweaver-Burk) plot of initial reaction versus the concentration of *p*NPS at different phenol concentrations. **d**) Replot of slope of the double-reciprocal (Lineweaver-Burk) plot of (**c**) versus phenol concentration (left) and of the y-intercepts of the same double-reciprocal plot versus 1/[phenol] (right). The slope replot provides a y-intercept equal to *K*_mA_/V_max_ and an x-intercept corresponding to –*K*_i_. The intercept replot yields a y-intercept corresponding to 1/ V_max_ and an x-intercept corresponding to –1/*K*_m,phenol_.

Collectively, these results demonstrate that St_ASST_5 follows a ping-pong catalytic mechanism and is subject to phenol-dependent substrate inhibition, with quantitative kinetic parameters consistent with competitive interaction of phenol at elevated concentrations. These relationships are described by the ping-pong bi-bi rate equation incorporating substrate inhibition:

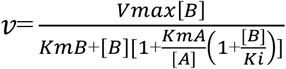

and its Lineweaver–Burk form:

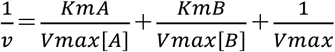

where [A] and [B] denote the concentrations of *p*NPS and phenol, respectively. To determine true kinetic constants, secondary replots derived from the Lineweaver-Burk analysis were evaluated by linear regression. Plotting the y-intercepts of the double-reciprocal lines as a function of 1/[phenol] at low phenol concentrations yielded a straight line, where the y-intercept corresponds to 1/V_max_ and the x-intercept to −1/*K*_m,phenol_ (Figure 3d right). In addition, plotting the slopes of the Lineweaver-Burk lines obtained under inhibitory conditions against phenol concentration produced a linear relationship in which the y-intercept represents K_mA_/V_max_ and the x-intercept corresponds to −*K*_i_, consistent with competitive inhibition (Figure 3d left) (79).

These analyses yielded the following parameters under the specified assay conditions: V_max_ = 0.35 µM·s⁻¹, *K*_m,*p*NPS_ = 60.9 µM, *K*_m,phenol_ = 12.61 mM, and *K*_i_,_phenol_ = 69.83 mM. The corresponding catalytic turnover number was *k*_cat_ = 9.8 s⁻¹.

#### Reaction product analysis by LC-MS and NMR

In the aim to define the regio-selectivity of the reaction on an acceptor bearing several hydroxyl and amine groups, in this case dopamine, sulfated products were analyzed by LC-MS after 24 h of reaction. The results demonstrate that St_ASST_5 is able to sulfate dopamine, leading to the formation of a sulfated product detected at *m/z* 232.0285 ([M-H]^-^) (Figure 4a, top right). Importantly, this ion was observed exclusively in the presence of both the enzyme and the donor *p*NPS, confirming that sulfation is enzyme-dependent. The extracted ion chromatograms revealed that dopamine (*m/z* 152.0717, [M-H]^-^) was detected at low intensity, likely due to suboptimal ionization efficiency. In contrast, the sulfated dopamine displayed a clear signal at a retention time of 1.84 min, with a significantly higher intensity (∼10⁷), supporting efficient conversion under the tested conditions. Notably, only a single sulfated product was observed.

**Figure 4:**
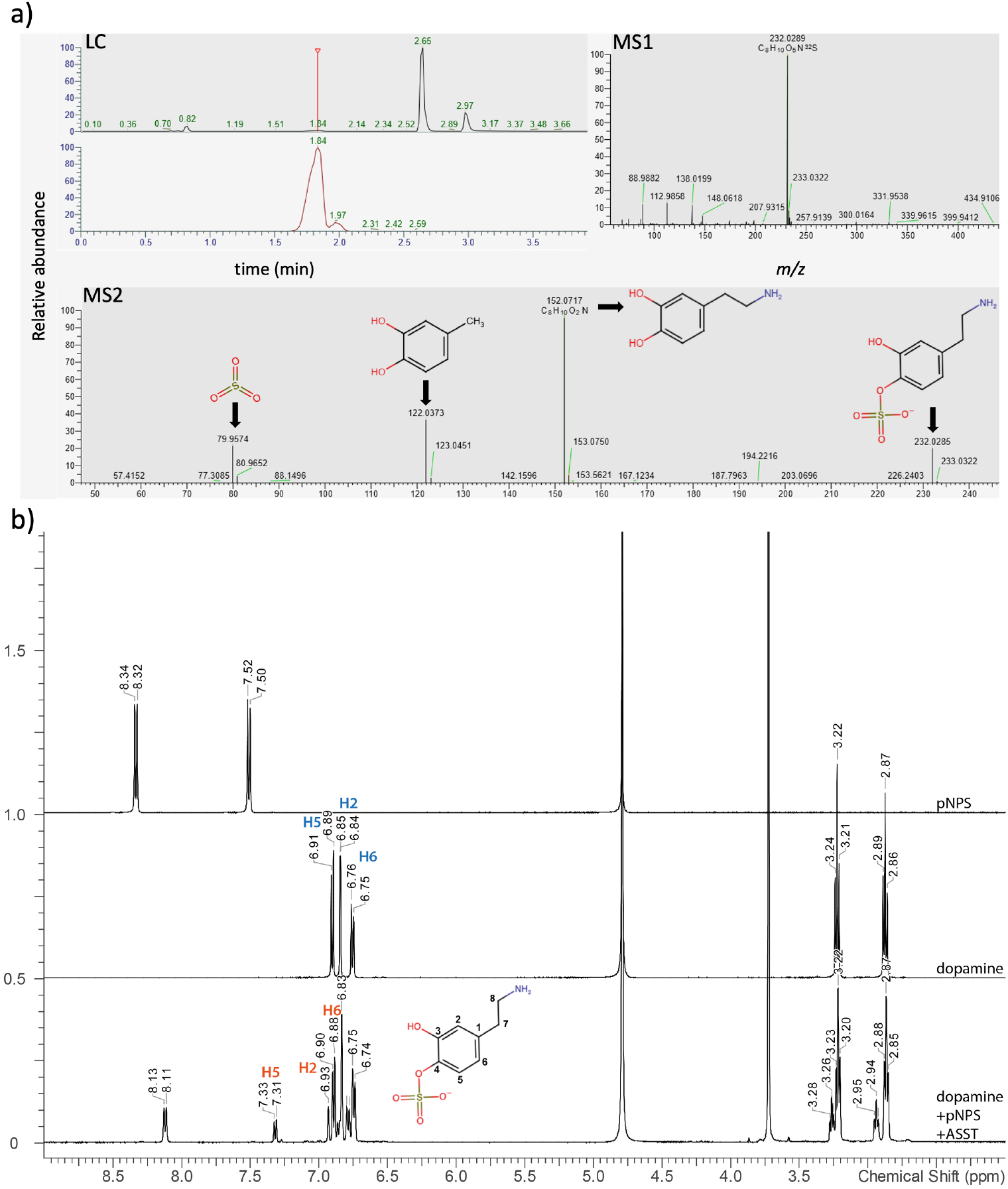
Detection of dopamine sulfated by St_ASST_5. Detection of phenolic substrates. **(a)** LC-MS/MS analysis of St_ASST_5 after 24 h of reaction. LC: liquid chromatography showing the relative abundance of the substrate as a function of retention time (min). MS1: detection of sulfated dopamine at *m/z* 232.0289. MS2: (collision energy 35%) yielded fragment ions at *m/z* 152.0717 (dopamine), *m/z* 79.9574 (sulfo group), and *m/z* 122.0373 (dopamine-derived fragment, likely corresponding to the loss of a C₁H₂N moiety). Note that dopamine alone was poorly detected under these conditions. **(b)** Determination of the sulfation position of dopamine by NMR. The chemical shift depends on their distance to the deshielding O-SO_3_^-^ group, allowing to assign the product being 4-O-sulfated dopamine, as described in (30,31). Dopamine-4-O-sulfate: δ 2.94 (t, J = 7.0 Hz, 2H, H-7), 3.26 (t, J = 7.1 Hz, 2H, H-8), 6.86 (dd, J = 8.2, Hz, 1H, H-6), 6.93 (d, J = not determined, 1H, H-2), 7.32 (d, J = 8.3 Hz, 1H, H-5). *p*-NP: δ 6.80 (d, J = 8.6 Hz, 1H, H-3), δ 8.12 (d, J = 8.8 Hz, 1H, H-2).

MS/MS fragmentation of the ion at *m/z* 232.0289 confirmed its identity as sulfated dopamine, yielding characteristic fragment ions at *m/z* 152.0717 (corresponding to dopamine), *m/z* 79.9573 (sulfo-group), and *m/z* 122.0373 likely corresponds to a dopamine-derived fragment resulting from the loss of C_1_H_2_N moiety (Figure 4a, bottom).

Analysis of the donor substrate showed the presence of *p*NPS (*m/z* 217.9768, [M-H]^-^), the signal of which decreased in the presence of the enzyme, consistent with its consumption during the reaction. The desulfated product *p*NP (*m/z* 138.0197) was also detected, with a strong increase in intensity in samples containing the enzyme, further supporting enzymatic turnover.

Overall, these results confirm that St_ASST_5 catalyzes the sulfation of dopamine, producing a single sulfated species. The position of sulfation was subsequently determined by NMR, which identified the 4-OH position as the site of modification (Figures 4b and S5).

#### Crystal structure of St_ASST_5

Well-diffracting crystals of St_ASST_5 were obtained in hanging drops, mixing 2 µL of protein with 2 µL of well-solution that contained 1.12-1.19 M AmSO_4_ and 1 M Na acetate at pH 4.6. The native form yielded crystals that diffracted between 1.6 Å and 1.7 Å resolution. In addition to the native structures, soaking experiments with the substrate donor *p*NPS (as powder) and the acceptor dopamine (dopamine, powder) resulted in complexed crystals that diffracted at 1.8 Å (St_ASST_5_*p*NPS) and 1.6 Å (St_ASST_5_dopamine), respectively. All crystals of St_ASST_5 crystallized with the space group P3_2_21; unit cell parameters are reported in Table 1.

**Table 1.** Data collection and refinement statistics *Values in parentheses are for the highest-resolution shell.

| Dataset name | St_ASST_5 | St_ASST_5_pNPS | St_ASST_5_dopamine |
| --- | --- | --- | --- |
| Data collection | PX2 (17/07/2024) | PX1 (24/05/2025) | PX1 (18/07/2025) |
| Space group | P3<sub>2</sub>21 | P3<sub>2</sub>21 | P3<sub>2</sub>21 |
| Unit cell parameters<br>(Å) | a=b=160.84, c=69.22 | a=b=160.41, c=69.27 | a=b=160.40, c=69.11 |
| Resolution (Å) | 46.43-1.86 (1.91-1.86) | 46.31-1.80 (1.85-1.80) | 48.99-1.60 (1.64-1.60) |
| $R_{pim}$ | 0.022 (0.364) | 0.021 (0.409) | 0.025 (0.332) |
| CC<sub>1/2</sub> | 1.000 (0.913) | 1.000 (0.781) | 0.999 (0.934) |
| $I/\sigma(I)$ | 22.6 (2.1) | 22.9 (1.8) | 17.9 (2.0) |
| Completeness (%) | 99.9 (98.5) | 99.8 (97.7) | 100.0 (99.9) |
| Redundancy | 20.6 (20.5) | 20.6 (19.2) | 20.3 (13.9) |
| Unique reflections | 85548 (6179) | 93998 (6754) | 134129 (9839) |
| **Refinement** |  |  |  |
| Resolution (Å) | 46.43-1.86 | 46.31-1.80 | 48.99-1.60 |
| No. atoms |  |  |  |
| Protein | 4491 | 4556 | 4503 |
| Ligand/ion | 10 | 86 | 37 |
| Water | 409 | 406 | 603 |
| *R*work/*R*free | 17.07/19.17 | 17.83/20.69 | 15.45/17.53 |
| B-factors |  |  |  |
| Protein (Å<sup>2</sup>) | 31.56 | 30.14 | 26.51 |
| Ligand/ion (Å<sup>2</sup>) | 60.12 | 48.02 | 40.04 |
| Water (Å<sup>2</sup>) | 36.80 | 37.21 | 36.93 |
| R.m.s. deviations |  |  |  |
| Bond lengths (Å) | 0.011 | 0.012 | 0.012 |
| Bond angles (°) | 1.697 | 1.745 | 1.787 |
| Ramachandran plot |  |  |  |
| Favored (%) | 95.55 | 95.35 | 95.73 |
| Allowed (%) | 4.09 | 4.29 | 3.56 |
| Outliers (%) | 0.36 | 0.36 | 0.71 |
| **PDB code** | 29YN | 29YP | 30JA |

**Table 2.**
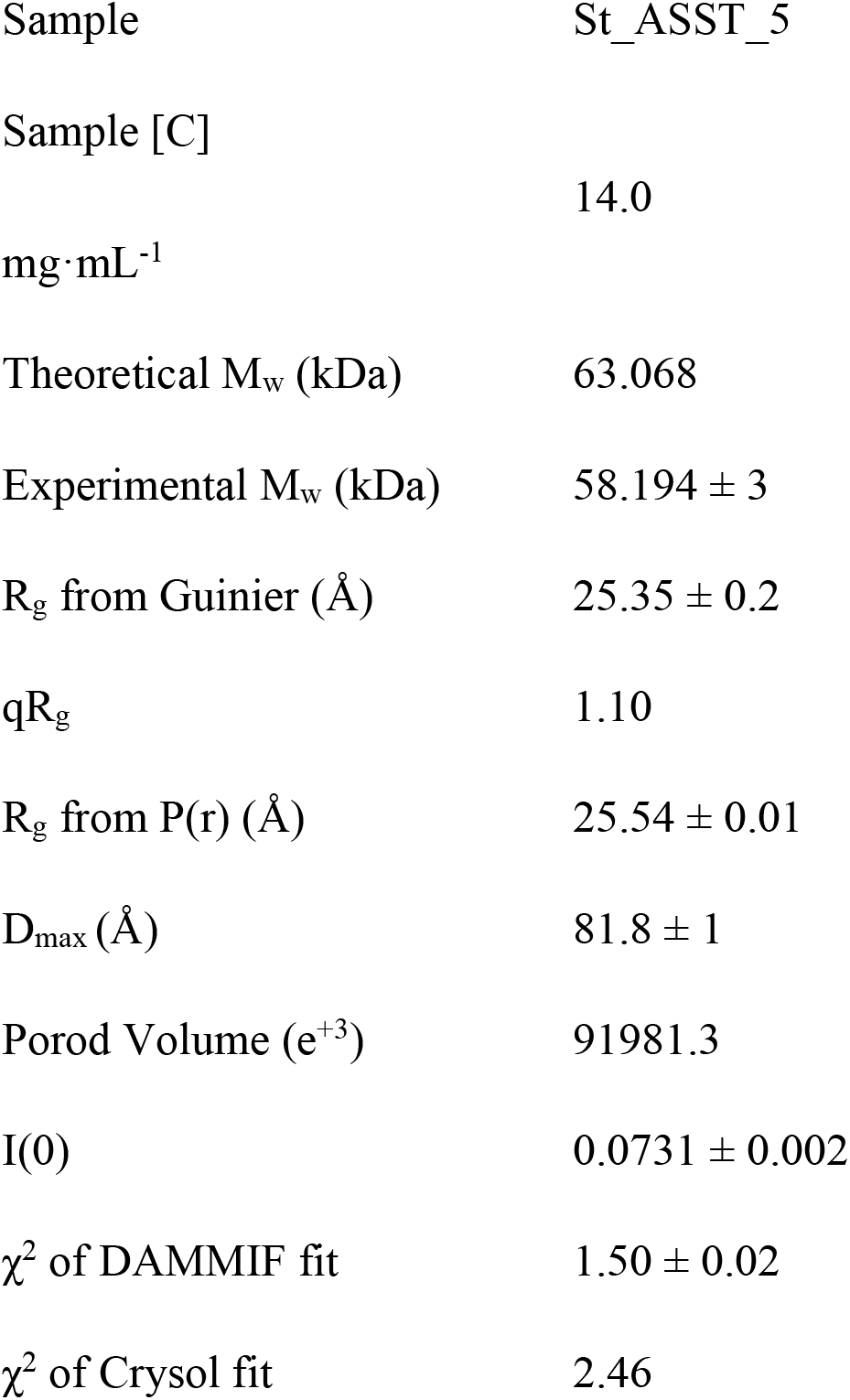
Experimental SAXS parameters derived from the scattering curves of St_ASST_5 in solution. The monomeric theoretical molecular weight is given as derived from the sequence.

The native-structure was solved by molecular replacement using the AlphaFold prediction as starting model (UniProt: A0A0D6HM77; moved to UniParc: UPI00000CCE18), while the substrate complexes were solved using St_ASST_5 as starting point. In all three St_ASST_5 structures, residues from 9 to 571 could be traced (Figure 5a-b) and are well defined by electron density, with the exception of a surface loop formed by residues 304-311 that was visible in the native structure, but for which no density was visible in the complexed structures. Notably, this disordered loop is located at the edge of a cavity in the center of the β-propeller, leading to the catalytic active site (Figure 5e).

**Figure 5.**
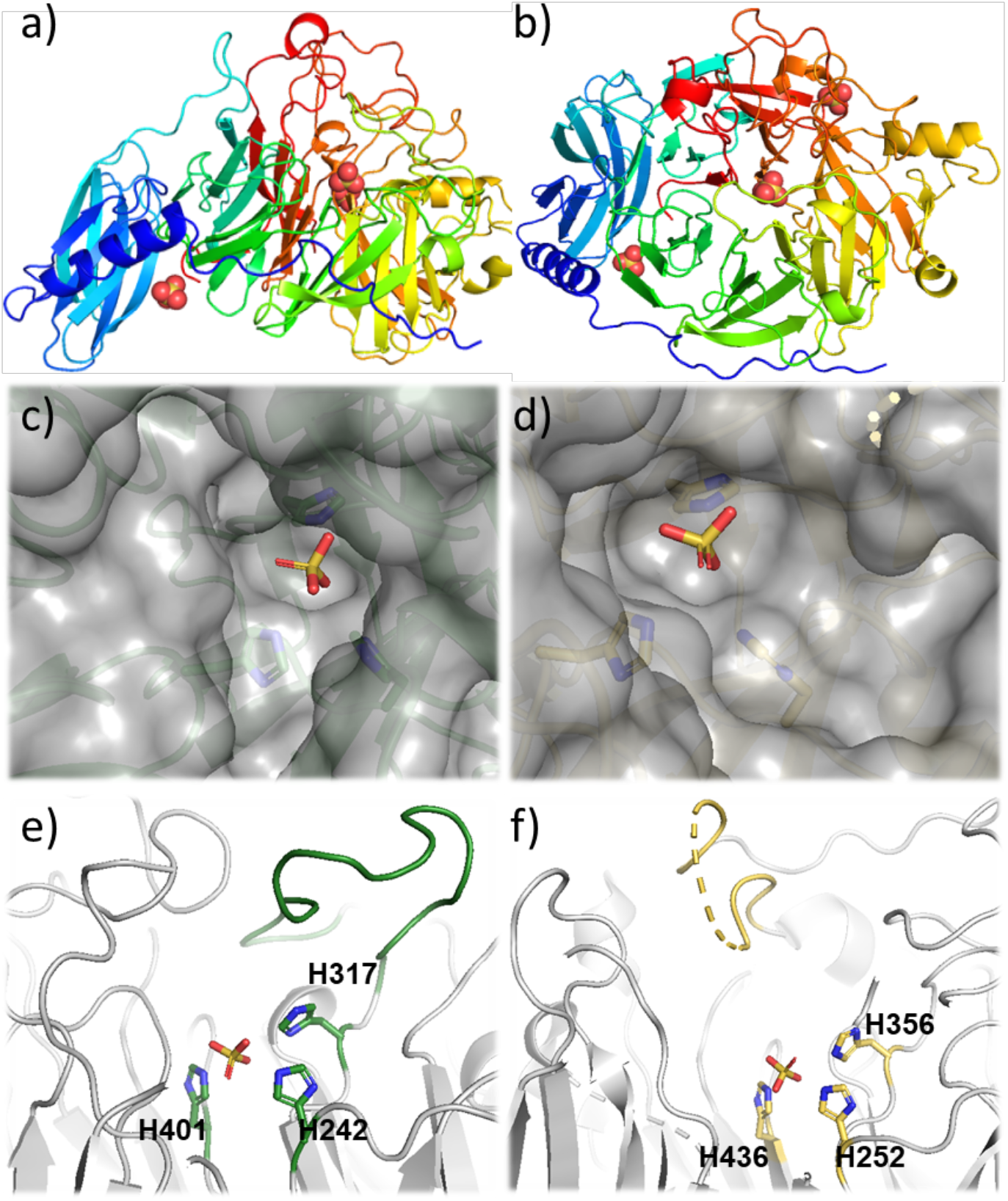
Comparison of the catalytic pockets of St_ASST_5 and Ec_ASST_1. **(a-b)** Overall structure of St_ASST_5. **(a)** Side view. **(b)** Top view of the catalytic site. **(c)** Surface representation of St_ASST_5 (green) showing a narrow and deep catalytic pocket. **(d)** Surface representation of Ec_ASST_1 (gray) highlighting a wider and shallower catalytic pocket. **(e-f)** Close-up view of the loop located above the catalytic site, previously described in Ec_ASST_1 as involved in catalysis, together with the three catalytic histidines and a sulfate ion. **(e)** St_ASST_5. **(f)** Ec_ASST_1. The Figure was drawn using PYMOL (13).

As expected, ASSTs are mainly composed of β-sheets and few α-helices. The native and complexed structures crystallized with one monomer in the asymmetric unit. The enzyme structure has the overall dimensions of 72.6 × 49 × 45 Å and is composed of two subunits, a C-terminal catalytic domain formed by residues 137-571 and an additional domain formed by residues 1-136. The catalytic domain has a typical β-propeller fold, consisting of six anti-parallel β-sheets and every β-sheet is composed of 4 β-strands. Moreover, additional helices are positioned after the third and fourth β-sheets, respectively. The additional domain has a typical β-sandwich fold consisting of seven anti-parallel β-strands and one long helix that connects the additional domain to the catalytic domain (Figure 5a-b). Notably, the precise function of this additional domain remains unknown to date.

#### Active site of St_ASST_5

The active site of St_ASST_5 is located in the central channel of the β-propeller, in coherence with the location observed in the structure of Ec_ASST_1 (12). The active site is accessible from the surface by a pocket with dimensions of 24 Å in height and a cross-section of 17 × 20 Å (Figure 5c). Three histidine residues at the bottom of the pocket, strictly conserved (except for very few sequences in clade 2), namely His242, 317 and 401 in St_ASST_5, form the catalytic active site (Figure 6a, b). These residues have been described as being involved in the catalytic mechanism in Ec_ASST_1 (12,14) (Figure 6d, f). Notably, in the St_ASST_5 and St_ASST_5_dopamine structures, a sulfate group is located close to His401 (3.5 Å) that is further hydrogen bonded to the other two histidines (His242 and His317), but also to two asparagines (Asn319 and Asn421) and one arginine (Arg335). In all crystal structures, additional sulfate groups have been identified, situated close to charged residues.

**Figure 6.**
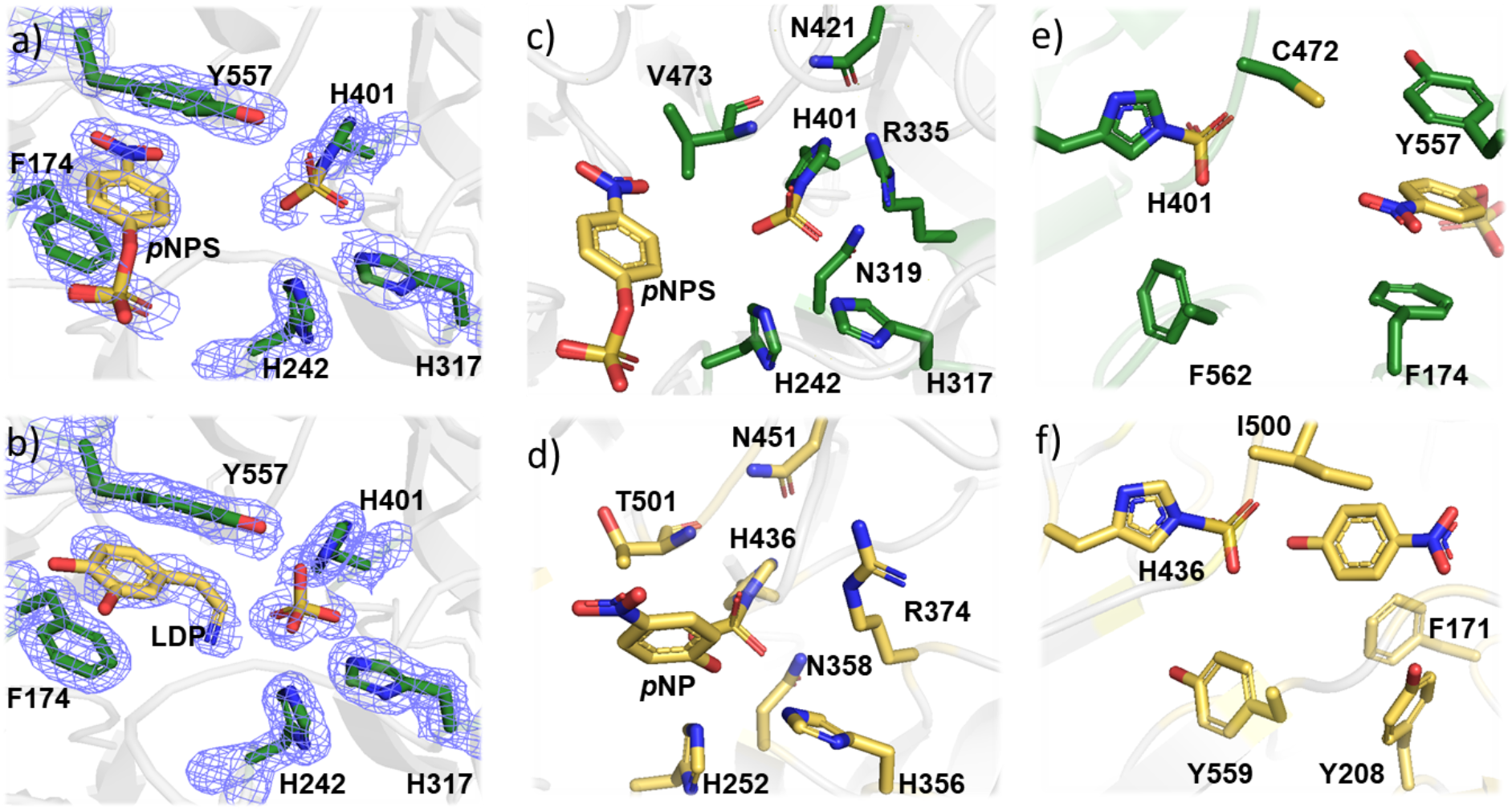
Crystal structure of St_ASST_5 alone and in complex with substrate molecules. **(a-b)** Catalytic site of St_ASST_5 showing the aromatic residues F-174 and Y-557 involved in substrate binding, together with electron density maps illustrating the accuracy of the models. **(a)** Complex with *p*NPS, well defined in the electron density. **(b)** Complex with dopamine, showing weaker electron density. **(c-d)** Analysis of residues involved in sulfate stabilization within the catalytic site. **(c)** St_ASST_5. **(d)** Ec_ASST_1. **(e-f)** Analysis of residues involved in substrate (*p*NP or *p*NPS) stabilization within the catalytic site. **(e)** St_ASST_5. **(f)** Ec_ASST_1.

Subsequently, the St_ASST_5_*p*NPS complex allowed to determine the structure of the sulfo-histidine catalytic intermediate. In the active site of the complex, the sulfate ion is covalently bound to N^e2^ of His401, demonstrating that it is the catalytic histidine (Figure 6a, c). The sulfo-histidine is stabilized by extensive hydrogen bonding to the side chains of the same residues that stabilized the ion sulfate in the native structure. Moreover, the loop spanning residues Pro301 to Thr311, ordered in the native structure, was disordered in all complex structures with no detectable electron density, and is located above the active site and corresponds to a region that was proposed to play a role in the recognition of natural substrates of Ec_ASST_1 (Figure 5f). These observations suggest that the substrate, even though artificial, used in this study may have induced local disorder, specifically affecting this loop. In addition, no reaction product, *p*NP, was detected in the catalytic site, nor in any of the structures we determined, in contrast to what was observed by Malojcic *et al.* (12,14). However, one molecule of the intact sulfated substrate donor, *p*NPS, was noticed near the catalytic site (Figure 6a, c). Surprisingly, the sulfo-group was oriented toward the outside of the active site. This donor molecule was roughly located at a distance of approximately 13.2 Å to the catalytic sulfated His401, and the sulfo-group was in hydrogen bonding distance with the main chain of Gly307. In our third structure of St_ASST_5, in complex with dopamine (Figure 6b), the acceptor molecule took the same position as *p*NPS, sandwiched between two aromatic residues, with the amino group orientated towards the catalytic active site, at approximately 3.5 Å from the sulfate ion near His401. Notably, the electron density was not perfectly defined, indicating some rotational disorder of the acceptor molecule, but indicating that the majority of the ethyl-amine extension pointed towards the active site (Figure 6b). None of the hydroxyl- or amine-groups formed additional hydrogen bonds with other residues, which may explain the partially disordered position of the substrate molecule in the structure.

Both the donor and acceptor substrates were wedged between two aromatic residues, Phe174 and Tyr557. Notably, the two aromatic residues are highly conserved within clade 5, at 99.8% for phenylalanine and 98% for tyrosine. The analyses of sequence alignments of the 22 clades revealed the absence of these two aromatic residues at homologous positions in all of the other clades (data not shown). This particular feature of clade 5 suggests a potential functional role in specific substrate recognition. Nonetheless, our crystal structures appear to have trapped unproductive configurations of donor and acceptor binding, since in both cases the reacting groups are not oriented as expected. This raises the question about binding site flexibility and necessary movements to allow substrates to be positioned such that the reaction can take place.

An additional difference between the St_ASST_5 and St_ASST_5_*p*NPS was observed on the surface of the two structures. A short loop next to *p*NPS was shifted by 8 Å between the two structures with a maximum distance observed at the position of Lys532 (Figure S7A).

Furthermore, another pair of aromatic amino acids (Phe458 and Tyr460) bind two *p*NPS molecules at the surface of the enzyme, far from the active site. Interestingly, these residues are conserved in more than 80 % in all 22 clades. Moreover, the analyses of the alignment of all representative sequences of the 22 clades (Barbeyron *et al*., in preparation) revealed additional residues conserved in more than 80% in ASSTs. Notably, these conserved residues reveal a tunnel in the catalytic domain of ASST structures that runs from below the active site into the center of the catalytic site (Figure S7B), suggesting the hypothesis that this tunnel could play the role of an additional substrate entry or leaving point into and from the active site. This remains to be demonstrated, especially since substrate molecules can access directly above the catalytic site, which is open and exposed.

#### Docking analyses in the active site and analyses of the structural “tunnel” in St_ASST_5

Docking was performed to generate suitable starting structures for the phenol- and dopamine-bound St_ASST_5 systems. Phenol could be docked readily because it contains only one reactive hydroxyl group and therefore has a clearly defined orientation for potential sulfonate-transfer. In contrast, dopamine was more difficult to place in a catalytically meaningful orientation because several functional groups could, in principle, participate in the reaction, including the two hydroxyl groups of the catechol-moiety and the amino group (See Table S4).

Although a crystal structure of St_ASST_5 with bound dopamine is available, dopamine adopts an orientation in which the amino group points toward the catalytically relevant histidines. This pose was considered unlikely to represent the productive binding mode for O-sulfonation, because the hydroxyl groups of the catechol-moiety are not ideally positioned towards the sulfonated His401 intermediate. Therefore, docking was used to explore alternative dopamine orientations and to identify a catalytically competent pose.

Docking poses were not selected based on docking score only, but according to geometric criteria relevant for sulfo-transfer (see section 4 in SI). Catalytically competent poses were defined by positioning of a substrate hydroxl-group toward the sulfur atom of the sulfonated His401 residue and by favorable orientation within the active-site histidine network (Figure S9 A). The final phenol and dopamine positions fulfilled these criteria and placed the reactive hydroxyl group in proximity to the sulfo-histidine intermediate (Figure 7a, b). The corresponding docking scores were -3.2345 kcal/mol and -4.3187 kcal/mol for phenol and dopamine, respectively.

**Figure 7.**
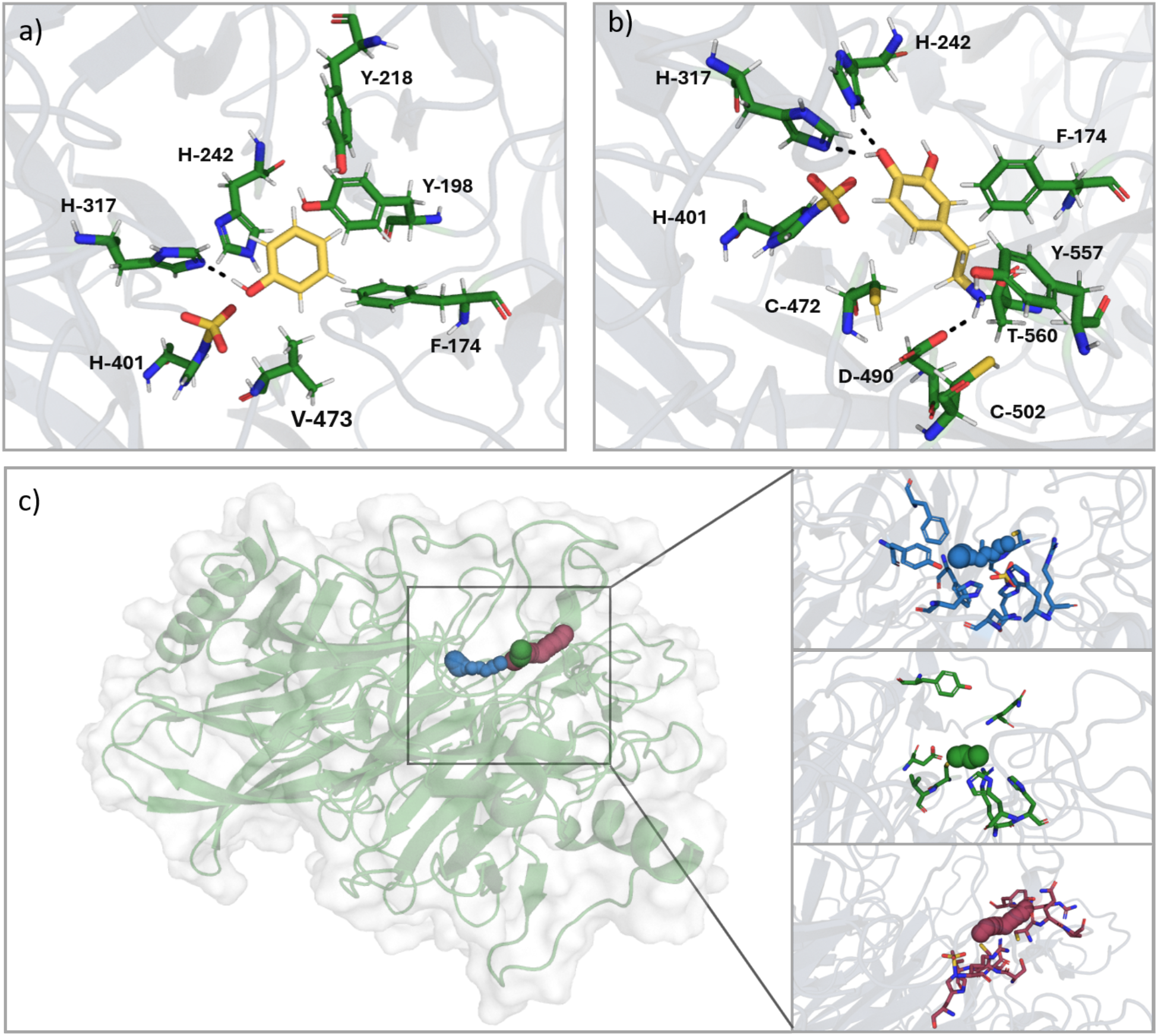
Docking poses and tunnel analysis of phenol and dopamine in St_ASST_5. **(a)** Final selected docking pose of phenol in the active site. The pose was chosen based on the orientation of the sulfated position toward the catalytically relevant histidines, using prior structural information from Dh_ASST_6. **(b)** Final selected docking pose of dopamine. The sulfated position is oriented toward the histidines of interest, representing a catalytically relevant binding mode. Key active-site and tunnel-lining residues are shown and labeled. **(c)** CAVER tunnel analysis of the selected phenol- and dopamine-bound structures using a minimum probe radius of 0.7 Å. The phenol tunnel is shown in blue, the first dopamine tunnel in green, and the second dopamine tunnel in magenta. The tunnel analysis shows one narrow access pathway for phenol, while dopamine displays two possible tunnel pathways.

For dopamine, the selected catalytically competent docking pose was also the best-scoring pose among the inspected docking solutions. In this orientation, the catechol hydroxyl group assigned as Position 1 (Figures S8, S9) is positioned between His317 and the sulfonated His401 intermediate, with the hydroxyl proton directed toward His317 and the oxygen of the catechol-moiety oriented toward the sulfate sulfur. This arrangement provides the required geometry for His317-mediated proton abstraction and subsequent S–O bond formation. The productive pose is further stabilized by the surrounding active-site environment. His242 and Asn319 contribute to the polar environment around the catechol/sulfo-group region, while Arg335 supports stabilization of the sulfonated intermediate. In addition, Cys472 is positioned close to the ring of the catechol-moiety and may contribute an S···π-type contact, whereas Val473 and Phe562 help define the hydrophobic pocket around the aromatic moiety (Figure S10). The aminoethyl substituent is oriented away from the catalytic His317/His401 region and toward the pocket entrance, where it can form a polar contact with Asp490. This interaction pattern stabilizes dopamine in a productive orientation while disfavoring nonproductive placement of the amino group in the catalytic region.

Tunnel analysis was carried out for the selected phenol- and dopamine-bound structures using CAVER (55) with a minimum probe radius of 0.6 Å (Table S6). For phenol, one main tunnel cluster was identified (Figure 7c). This tunnel had a throughput of 0.42, a length of 6.64 Å, a curvature of 1.28, and a bottleneck radius of 0.74 Å. Since the bottleneck radius is only slightly larger than the applied probe radius, this pathway represents a narrow access route for the phenol-binding site. Such a restricted pathway may help to confine phenol close to the sulfonated His401 intermediate and reduce the number of possible nonproductive orientations. The tunnel is lined by active-site and substrate-binding residues, including Phe174, Tyr198, His242, His317, Asn319, Arg335, His401, Cys472, and Phe562 (Table S7). Aromatic and hydrophobic residues form a compact environment around phenol, while polar and charged residues are positioned along the same pathway and may contribute to substrate orientation. This arrangement suggests that phenol binding is not only governed by substrate access, but also by local preorganization within the catalytic pocket.

For dopamine, two tunnel clusters were detected (Figure 7c). The main tunnel showed a higher throughput than the phenol tunnel (0.78 vs. 0.42), was shorter (3.29 Å vs. 6.64 Å), and has a larger bottleneck radius (0.86 Å vs. 0.74 Å). This indicates a more accessible entry route in the dopamine-bound structure (Table S6). Such increased accessibility may be important for accommodating dopamine, which is larger and more flexible than phenol and contains several potentially reactive groups, including two catechol hydroxyl groups and an amino group. At the same time, this flexibility may also increase the probability of nonproductive binding orientations, consistent with the experimentally observed dopamine-bound structure in which the amino group points toward the catalytic histidines. The main dopamine tunnel was lined by His317, Arg335, His336, His401, Cys472, Val473, Asp490, Gln503, and Tyr557 (Table S7). A second dopamine tunnel was also observed, but this pathway had a lower throughput of 0.26, a longer length of 8.66 Å, and a smaller bottleneck radius of 0.61 Å, indicating that it is less accessible than the main dopamine pathway. The presence of this additional, less favorable route nevertheless suggests that dopamine may sample a broader region of the active-site entrance than phenol. This could contribute to the broader substrate tolerance of St_ASST_5, but productive sulfonate-transfer still requires selection of a catalytically competent pose in which one catechol hydroxyl group is oriented toward the sulfonated His401 residue. More information can be found ins section 6 of SI.

The differences observed in the substrate access pathway suggest that St_ASST_5 can accommodate chemically distinct acceptor substrates through a flexible binding environment. While phenol is a small and rigid substrate that fits into a confined aromatic pocket, dopamine requires additional conformational freedom due to its larger size, flexible aminoethyl side chain, and multiple potentially reactive functional groups. The restricted phenol pathway may favor a defined binding orientation by limiting the number of possible conformations close to the sulfonated His401 intermediate. In contrast, the more accessible dopamine tunnel may facilitate the entry and accommodation of the larger catechol substrate but also allows multiple binding orientations, including nonproductive states as observed in the available dopamine-bound crystal structure. Similar relationships between tunnel architecture, active-site accessibility, and substrate selectivity have been described for other enzymes, where changes in access pathways influence ligand migration, positioning, and catalytic efficiency (80,81). That means, the additional dopamine access route may contribute to the broader substrate tolerance of St_ASST_5 while productive sulfo-transfer still depends on correct positioning of the acceptor hydroxyl group within the catalytic histidine network.

#### Analysis of genomic context of St_ASST_5

In order to investigate whether a hint towards the function of ST_ASST_5 in *Salmonella typhimurium* could be inferred from its genome, comparative genomic analysis was carried out using the MaGe (Magnifying Genomes) platform to explore its genomic context; based on the hypothesis that ASST genes are part of functional operons rather than isolated entities, we examined the genomic neighborhood of St_ASST_5. For this matter a 20 kb genomic window surrounding the St_ASST_5 gene was examined and revealed a conserved gene cluster potentially linked to sulfate metabolism and pH regulation (Figure S6). This cluster includes NhaR, a transcriptional activator of the sodium/proton antiporter NhaA, both showing high sequence similarity to their *E. coli* homologs (UniProt accession codes: P0A9G2, A9MYF8). The presence of these genes might suggest a connection between St_ASST_5 activity and ionic homeostasis.

In addition, the cluster contains a sulfatase maturation enzyme (SAM) with a predicted radical SAM domain, as well as a sulfatase of the S1_23 family (Uniprot: A0A3U5JH81, present in https://Abims.SulfAtlas). The co-localization of these genes with St_ASST_5 supports the existence of a functional operon dedicated to the synthesis, modification, and regulation of sulfated compounds. A second ASST gene, also classified within clade 5, and located next to St_ASST_5, further strengthens the hypothesis of a sulfate metabolism cluster possibly coupled to pH regulation mechanisms, but did not reveal further details allowing to better define a functional role.

#### Structure comparison of St_ASST_5 to other ASSTs

The structure of St_ASST_5 was compared with the only published ASST structure available to date, namely Ec_ASST_1 (3ELQ, 12) (Figure S11). As expected, both ASSTs share a similar overall fold, and each enzyme comprises two domains. The C-terminal catalytic domain adopts a β-propeller architecture, and superimposition gives a root mean square deviation (rmsd) of 2.4 Å and a Z-score of 7.1 between the two catalytic domains, reflecting a reasonable structural similarity (Figure S11a, b). The additional N-terminal domains of both enzymes exhibit a typical β-sandwich fold composed of seven antiparallel β-strands, with an additional α-helix in that of Ec_ASST_1. A major difference between the two structures is the presence of an extended α-helix connecting the two domains in St_ASST_5 that is absent in Ec_ASST_1 (Figure S11a, b). The rmsd between the structures of the additional domains is 2.7 Å. The precise function of the additional domains remains unclear, although its conservation suggests a possible role in substrate recognition or protein stability.

To investigate further potential functions of the additional domains, a fold seek was performed using DALI (51), but the structures of the best hits in the PDB do not add any valuable information. For example, the first hit (2QBW, rmsd of 2.2 and Z-score 8.6) is an adnectin domain that is involved in protein-protein interactions of a complex that controls plasma LDL-cholesterol levels (82). While this similarity might point towards the possibility that these additional domains are involved in protein-protein interactions, the target of these interactions remain unknown.

The catalytic active sites of Ec_ASST_1 and St_ASST_5 display similar overall architectures, both containing three conserved histidine residues at the center of the β-propeller domain and characteristic of ASST enzymes (Figure 5e, f; S11), further surrounded by several loops that are variable. In Ec_ASST_1, the side chains of Asn358, Arg374, and Thr501 are implicated in hydrogen-bond formation and in coordination of the sulfate group during catalysis. The corresponding residues in St_ASST_5 are Asn319 and Arg335, while Thr501 is replaced by Val473 (Figure 6c, d). In addition, an extra asparagine residue (Asn421) is present in St_ASST_5 and aligns with Asn451 in Ec_ASST_1, suggesting a possible role in hydrogen bonding described previously. Two hydrophobic residues (Phe171 and Ile500) that contribute to the catalytic pocket in Ec_ASST_1 are missing in St_ASST_5 (Figure 6e, f).

The major differences between the two structures are located in the loops connecting the different propeller blades, all positioned around and above the active site (Figure 5a and S11A). Notably, a flexible loop, Val321 to Leu327 in Ec_ASST_1 that was associated to catalytic activity (12), is also present in St_ASST_5, but involves a different sequence stretch and occupies a different region (Pro301 to Thr311). This change of location likely affects substrate orientation and accessibility and most likely contributes to the observed differences in substrate specificity between the two enzymes (Figure 5e, f).

In the Ec_ASST_1 structure complexed with *p*NPS, a de-sulfated donor (*p*NP) is positioned within the catalytic pocket, whereas in St_ASST_5, no *p*NP molecule is present. Instead, the intact *p*NPS molecule is located adjacent to the catalytic cleft, stacked between two aromatic residues, Phe174 and Tyr557, thus involved in substrate binding. At the equivalent position, Ec_ASST_1 contains Tyr208 but oriented outward from the pocket and lacks a residue corresponding to Tyr557 of St_ASST_5. Additionally, no residues involved in *p*NP substrate binding were described in Ec_ASST_1 (Figure 6 d, f).

The Protein Data Bank (PDB) does contain another crystal structure of a bacterial ASST belonging to clade 14 and originating from a *Streptomyces sp*., 9KF6 (released in November 2025), however for which no describing publication is reported so far. Nevertheless, the superimposition with St_ASST_5 (Figure S12) results in a rmsd of 3.13 Å, witnessing of major differences. While the relative positions of the three histidines is well conserved (Figure S12C), the major differences between the two structures are located in the loops above the active site, that are all but for two much shorter than in St_ASST_5 (Figure S12A, B). The absence of these long loops results in a much wider and open active site (Figure S12D). Interestingly, this ASST was characterized to be active on liponucleosides, and caprazamycin derivatives in particular, explaining the need for a large cavity above the active site to accommodate these large molecules as acceptors (83). The residues and detailed structural determinants leading to this specificity, however, are not characterized to date.

#### Analyses of substrate binding residues in St_ASST_5 by site directed mutagenesis

To investigate the roles of the catalytic histidine and the aromatic residues that are strictly conserved in St_ASST_5 and involved in donor and acceptor substrate binding, site-directed mutagenesis was performed. The catalytic histidine His401 was substituted by glutamine (H401Q), while the aromatic residues were replaced by alanine, F174A and Y557A, as well as the double mutant F174A/Y557A. The corresponding mutant enzymes were heterologously expressed as His-tagged proteins and purified in their soluble form. Enzymatic activity assays were performed following the same protocol as described for the spectrophotometric assay using *p*NPS as donor and phenol as acceptor. The pure hydrolytic activity of pNPS in absence of acceptor was also measured (Figure 8a) and subtracted from the total activity. These activity assays revealed that mutation of His401 completely abolished enzymatic activity, confirming its essential catalytic role. In contrast, mutation of the aromatic residues led to a decrease in activity: the F174A mutant showed a reduction of approximately 76 % while Y557A and the double mutant F174A/Y557A exhibited up to 90 % decrease in activity compared to the wild-type enzyme (Figure 8b). The results of these mutations suggest a key role for Tyr557 in substrate recognition, potentially providing an additional hydrogen bond through the terminal -OH group, although this conclusion remains to be confirmed using physiological substrates.

**Figure 8.**
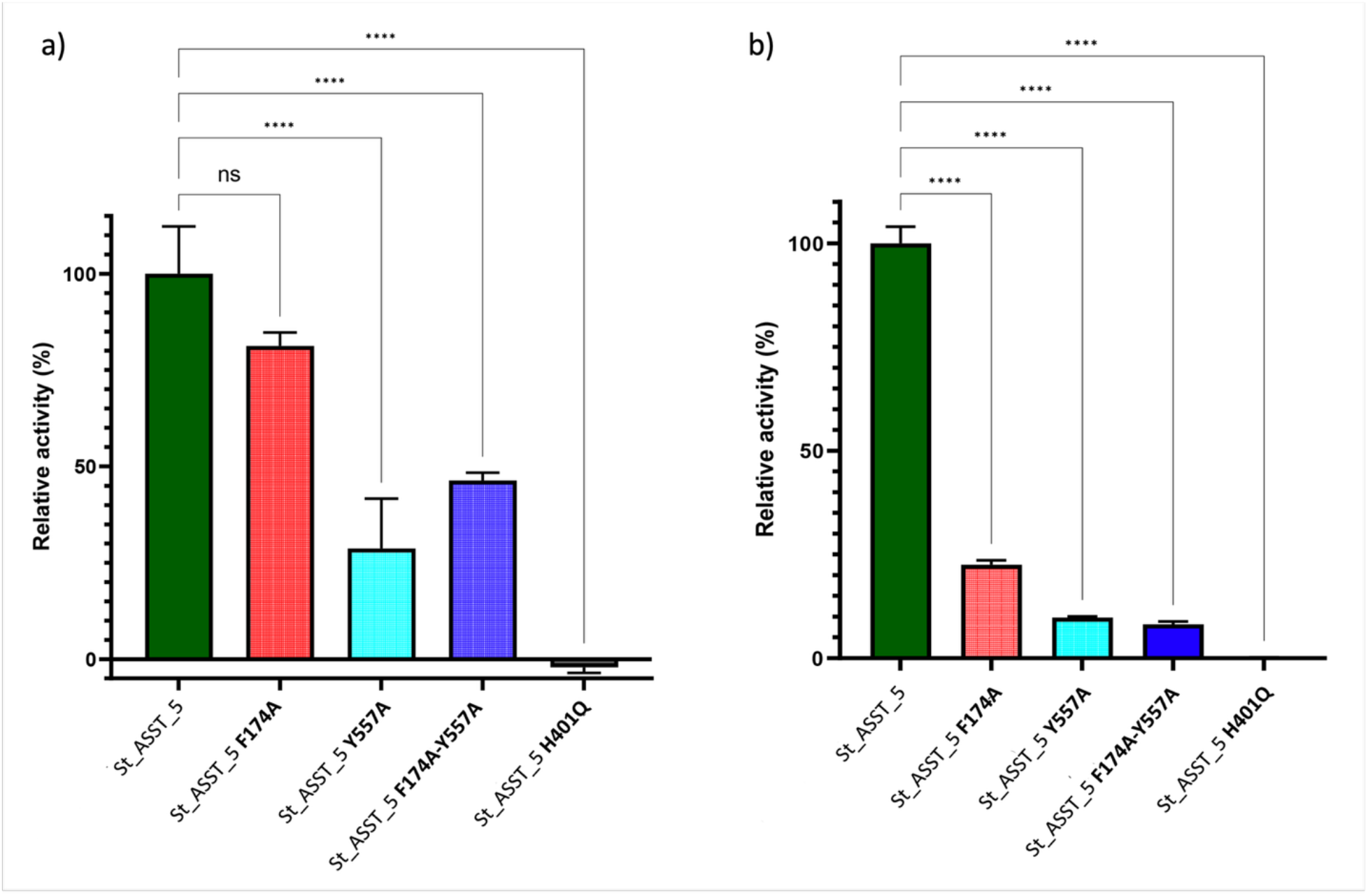
Activity measurements of active site mutants of St_ASST_5. (**a**) Relative hydrolytic activity (on *p*NPS without acceptor) of the mutants F174A, Y557A, H401Q and the double mutant F174A-Y557A of St_ASST5, as compared to the native enzyme. The release of *p*NP was measured in presence of the donor *p*NPS only. (**b**) Relative sulfate transfer activity of the mutants F174A, Y557A, H401Q and the double mutant F174A-Y557A as compared to the native enzyme. The activity was obtained by subtracting the hydrolytic activity from total activity in presence of both donor *p*NPS and phenol as acceptor.

#### Solution structure of St_ASST_5

To question further the role of the additional domains, we investigated their relative conformations in solution using small angle X-ray scattering (SAXS). The SAXS profile of St_ASST_5 displays the characteristic features of a mostly globular and compact monomeric protein (Figure 9a), with experimental Dmax and Rg values of 81.8 and 25.35 Å, respectively (Table 2). The overall shape derived from the scattering profile compares very well to the crystal structure (Crysol fit, χ^2^ of 2.46, Figure 9b), and the 10 *ab inito* models calculated by DAMMIF are highly similar (NSD of 1.0 calculated by DAMAVER), all of them also showing the presence of the additional domain (Figure 9b, c) and fitting the experimental curve with χ^2^ values ranging from 1.49 to 1.51. Overall, these results underpin that the structural arrangement of St_ASST_5 in the crystal structure and in solution are identical, and that the additional domain maintains a fixed position with respect to the catalytic β-propeller fold in solution, at least in absence of substrate.

**Figure 9.**
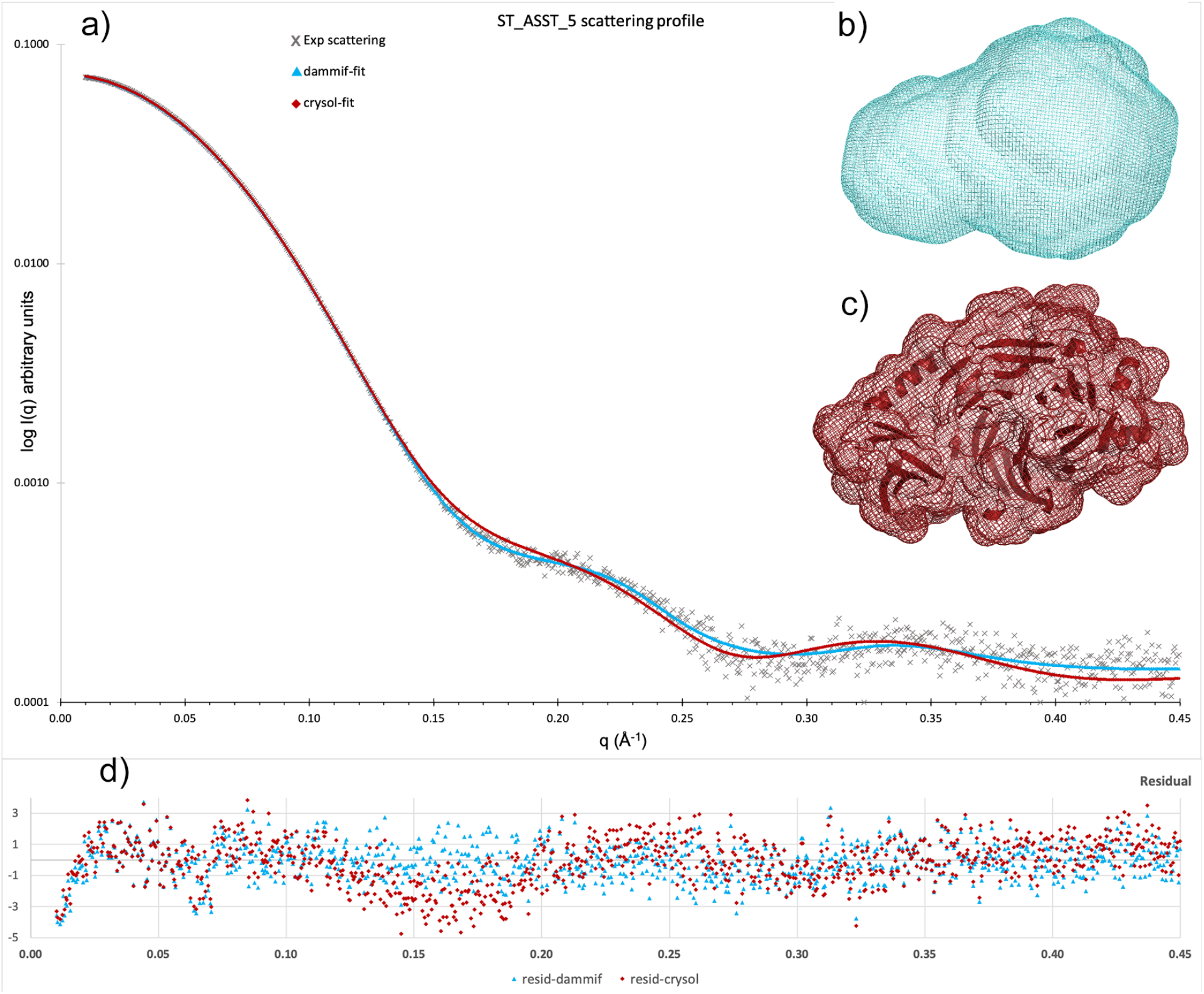
Overall shape of St_ASST_5 in solution measured by small angle X-ray scattering (SAXS). (**a)** Experimental scattering curve (grey crosses) fitted with DAMMIF (blue triangles) or Crysol (red diamonds). (**b)** Best representative shape among the 10 models calculated by DAMMIF giving a χ^2^ value of 1.5. (**c)** Surface representation of the crystal structure of St_ASST_5 that was fitted to the experimental curve using Crysol. (**d)** Residuals I(q)/sI(q) of the fitted curves shown in A) to the experimental data.

### 3QM/MM MD Free Energy Analysis of the second Sulfo Transfer step in St_ASST_5

To investigate the chemical step that determines acceptor sulfation in St_ASST_5, QM/MM umbrella sampling simulations were performed for phenol and dopamine. ASSTs catalyze a PAPS-independent sulfo-transfer reaction in which a sulfate group is transferred from an aryl sulfate donor to a phenolic acceptor. In a ping-pong bi-bi mechanism, the first half-reaction generates a covalently modified enzyme intermediate, while the second-half reaction transfers the group from this intermediate to the acceptor substrate and regenerates the enzyme. For St-ASST_5, the second step is therefore particularly important because it describes how the sulfonated His401 intermediate is resolved by phenolic acceptors and how different substrates can be transformed after formation of the sulfo-enzyme state. Similar sulfo-histidine intermediates and donor-bound states have been structurally characterized for bacterial ASSTs, but the detailed sequence of proton abstraction, S-O bond formation, and active-site stabilization during the acceptor sulfation step remains unclear. Prior QM/MM simulations, the systems underwent 3 independent MD runs. For more information refer to section 7 in SI.

The reaction was described using two distance-based collective variables (Figure S22). CV1 describes proton transfer from the substrate hydroxyl group to the catalytic histidine (His317) nitrogen, while CV2 describes formation of the S–O bond between the substrate oxygen and the sulfur atom of the sulfate group in His401. These coordinates describe a general base assisted sulfo-transfer from the sulfonated histidine intermediate to the acceptor hydroxyl group. In chemical term, the reaction corresponds to an asynchronous proton coupled sulfo-transfer, in which His317 activates the acceptor oxygen before or during nucleophilic attack at the sulfate sulfur. For more information, refer to section 8 in SI.

#### Phenol as Reference Substrate

For phenol, projection of the two-dimensional free-energy surface onto the sampled reaction pathway shows a continuous low-energy pathway from the reactant region toward the product region (Figure 10a). The reaction proceeds from the reactant state trough a shallow intermediate region before reaching the principal transition-state region, with an activation free energy of ΔG^‡^ = 10.45 kcal mol⁻¹. Analysis of the underlying CVs and representative structures shows that proton abstraction from the phenolic hydroxyl group by His317 occurs early along the reaction coordinate, whereas substantial shortening of the acceptor O-S distance occurs later. The shallow intermediate therefore represents a preorganized near-attack configuration rather than necessarily a stable chemical intermediate. The phenolic oxygen has already been activated by His317, while the S-O distance to the sulfated His401 donor remains comparatively long (Tabel S12 and S13).

**Figure 10:**
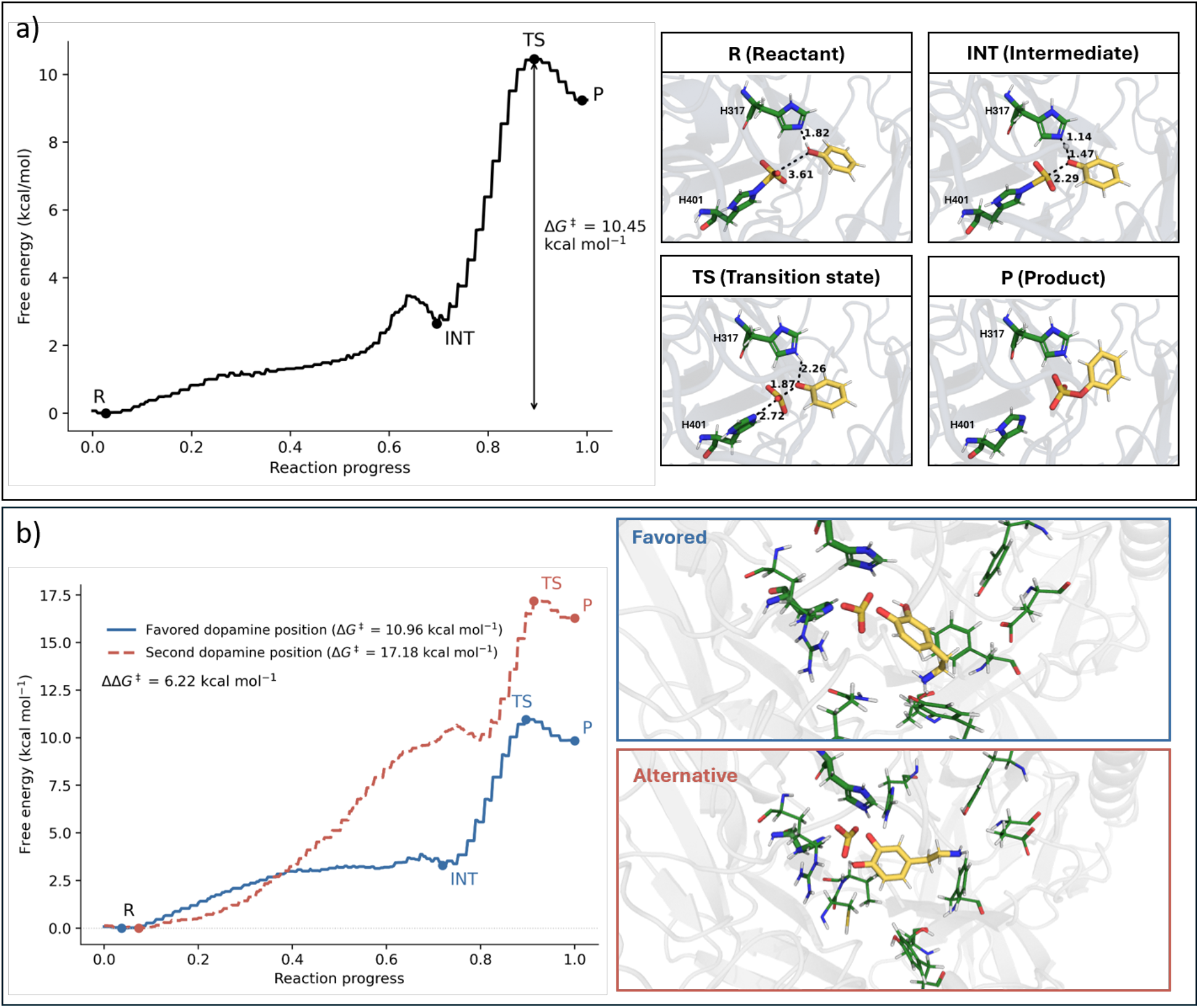
QM/MM free-energy profiles and representative structures for phenol and dopamine sulfo-transfer in St_ASST_5. **(a)** Reaction free energy profile for phenol along the sampled reaction path, with the reactant state (R), a shallow intermediate (INT), the transition state (TS), and the product state (P). The calculated activation free-energy barrier is ΔG^‡^ = 10.45 kcal mol⁻¹. Representative active-site structures corresponding to the states along the phenol pathway are shown on the right. **(b)** Comparison of the two possible dopamine sulfo-transfer pathways. The favored dopamine hydroxyl position (blue) displays a lower activations barrier of ΔG^‡^ = 10.96 kcal mol⁻¹, whereas reaction through the alternative hydroxyl position (red, dashed) requires ΔG^‡^ = 17.18 kcal mol⁻¹, yielding to ΔΔG^‡^ = 6.22 kcal mol⁻¹. Representative structures of the favors and alternative TS dopamine configurations are shown on the right.

The dominant free energy barrier within the modeled second half-reaction is associated primarily with S-O bond formation and concomitant reorganization of the sulfo group. The representative structures in Figure 10a show this progression. Phenol is already positioned in the catalytic pocket in the reactant state, His317 promotes early deprotonation of the acceptor hydroxyl group, and the transition-state region is reached only upon approach of the activated oxygen toward the sulfur center of sulfonated His401. These results support an asynchronous mechanism in which acceptor activation precedes the principal bond-forming event.

This mechanistic ordering is consistent with the general catalytic logic of sulfotransferases, in which activation of the acceptor hydroxyl group and an approximately in-line nucleophilic attack at sulfur facilitate sulfo-group transfer (12,83,84). St_ASST_5 differs fundamentally from canonical PAPS-dependent sulfotransferases, however, because the sulfo donor is a covalent sulfo-histidine intermediate generated during the first half-reaction of the ping-pong bi-bi catalytic cycle. Covalent histidine sulfation and ping-pong catalysis have previously been established for bacterial ASSTs (12,14), whereas the molecular sequence of acceptor activation and sulfo transfer during the second half-reaction has remained less well resolved. The present QM/MM umbrella-sampling calculations extend this structural framework by indicating that resolution of the sulfated His401 intermediate proceeds through early His317-mediated acceptor activation followed by the principal S-O bond-forming barrier.

A similar qualitative ordering was observed for Dh_ASST_6, where productive sulfo transfer likewise required formation of a near-attack configuration before crossing the principal bond-forming barrier. Thus, despite differences in enzyme scaffold, substrate environment, and computational sampling, both ASST systems support a central role for active-site preorganization and transition-state stabilization in PAPS-independent sulfo transfer (85–87). Further details of the two-dimensional free energy surface, collective-variable evolution, and representative states are provided in Section 9 of the Supporting Information.

#### Dopamine as Alternative Acceptor Substrate

The favored dopamine OH position follows the same chemical overall sequence as phenol. Proton transfer to His317 precedes extensive S-O bond formation, and the dominant free energy barrier is associated with sulfo-group transfer from sulfated His401 to the activated catechol group. The corresponding activation free energy is ΔG^‡^= 10.96 kcal mol⁻¹, only 0.51 kcal mol⁻¹ above the value obtained for phenol. This small difference should not be interpreted as a quantitatively exact prediction of relative reaction rates, rather, it indicates that once dopamine adopts a catalytically competent orientation, the energetic cost of the chemical transformation is comparable to that of phenol sulfation.

Importantly, the calculated potential of mean force describes only the second, acceptor-dependent half-reaction of the ping-pong mechanism, beginning from the preformed sulfo-His401 intermediate. It therefore does not represent the overall free energy change of sulfuryl transfer from the donor substrate to dopamine. Consequently, the product state lying above the reactant basin should not be interpreted as evidence that the complete catalytic reaction is thermodynamically unfavorable. Instead, the positive product-side free energy reflects the relative stability of the enzyme-bound states sampled along this isolated half-reaction and the chosen collective variables. The overall reaction thermodynamics would additionally include formation of the sulfoenzyme during the first half-reaction, as well as the substrate binding and product release.

To determine whether the experimentally observed regioselectivity could be rationalized at the level of the chemical free energy landscape, sulfo transfer through the second catechol hydroxyl group was evaluated independently. This alternative pathway exhibits a substantially higher activation free energy of ΔG^‡^= 17.18 kcal mol⁻¹, corresponding to a ΔΔG^‡^ of 6.22 kcal mol⁻¹ relative to the favored pathway (Figure 10b). Thus, the favored hydroxyl position is strongly preferred kinetically within the QM/MM model. The product-side free energy difference shows the same qualitative preference. The favored pathway reaches a product state at +9.85 kcal mol⁻¹ relative to its corresponding reactant basin, whereas the alternative pathway remains at +16.28 kcal mol⁻¹. The favored reaction is therefore also thermodynamically less unfavorable within the independently referenced PMFs.

The representative transition-state structures provide additional insight into the origin of the regioselectivity (Figure 10b). In both pathways, proton transfer to His317 is essentially complete in the transition-state configuration, with H···N distance of approximately 1.06–1.07 Å. Moreover, both configurations retain an approximately in-line geometry for sulfo transfer, with O-S-N(His401) angles of ∼173–174°. The energetic discrimination between the two catechol positions therefore does not appear to arise simply from loss of the canonical in-line attack geometry or failure of His317 to activate the acceptor.

Instead, switching the reacting catechol oxygen changes the orientation of dopamine within the active site and the degree of progression along the sulfo-transfer coordinate. In the favored transition-state structure, the forming O-S distance is 1.89 Å while the departing S···N(His401) distance has increased to 3.02 Å. In the alternative transition state, the corresponding distances are 1.96 and 2.79 Å, respectively. Thus, the favored transition state is somewhat further advanced toward formation of the product S-O bond and cleavage of the sulfo-His401 linkage. At the same time, the different catechol orientation reorganizes the surrounding interaction network. In the alternative configuration, the reacting oxygen is positioned close not only to His317 but also to the second active-site histidine, while the nonreacting catechol oxygen is redirected toward other polar residues of the pocket. The alternative pathway therefor requires a distinct redistribution of substrate-protein interactions to bring the second hydroxyl group into the same catalytic axis.

Differences in the intrinsic or enzyme-modulated acidity of the two catechol hydroxyl groups may additionally contribute to this regioselectivity. Dopamine possesses multiple coupled protonation equilibria, and its phenolic hydroxyl groups are not necessarily equivalent with respect to microscopic deprotonation behavior (88,89). More generally, studies of catechol-processing enzymes have shown that phenolic acidity can contribute to regioselectivity together with substrate orientation and active-site interactions.(90,91). Such an effect would be mechanistically plausible here because deprotonation of the reacting hydroxyl group by His317 precedes sulfo transfer.

These structural differences are consistent with the calculated free-energy profiles. The favored orientation permits dopamine to undergo His317-mediated deprotonation followed by sulfuryl transfer with a phenol-like activation barrier, whereas use of the alternative hydroxyl group requires substantially greater energetic investment to reorganize the substrate and active-site interaction network before transfer can proceed. The two catechol hydroxyl groups are therefore sterically accessible but not catalytically equivalent. Further details of the two-dimensional free-energy surfaces, collective-variable evolution, and representative states are provided in Section 10 of the Supporting Information.

Comparison of phenol and dopamine therefore indicates that St_ASST_5 uses the same fundamental His317/His401-mediated chemical strategy for both acceptors, while the additional regioselectivity calculation shows that not all dopamine hydroxyl orientations are chemically equivalent. The favored dopamine position retains a phenol-like barrier, whereas the second catechol hydroxyl position is separated by a substantially higher barrier. This supports a model in which substrate specificity and regioselectivity arise from formation and stabilization of a productive near-attack geometry rather than from substrate binding affinity alone.

## CONCLUSION

The crystallization of a PAPS-independent aryl-sulfate sulfotransferase, St_ASST_5, in presence of donor and acceptor substrates has provided valuable structural insight into the enzyme’s catalytic intermediate states. We report three distinct high-resolution structures representing the free enzyme at 1.9, the sulfonated enzyme intermediate at 1.8 and the enzyme-dopamine complex at 1.6 Å resolution that collectively capture pictures along the catalytic pathway. Combined with kinetics, biochemical characterization of the reaction product, site directed mutagenesis, MD simulations and QM/MM calculations this study sheds light on important residues, His401 and His317 in St_ASST_5, that mediate the sulfo-transfer. The study also highlights the importance of residues Phe174 and Tyr557, only present in St_ASST_5, that stabilize aromatic moieties of donor and acceptor molecules, and that thus appear to play a role in acceptor specificity through the stabilization of sulfo-transfer transition states in ASST’s belonging to this clade. The results show that dopamine, the preferred acceptor among those examined in this study and which has multiple functional groups, is selectively sulfonated at the 4-OH position. They also underpin the hypothesis that different clades within the large family of ASST’s coincide with different substrate specificities, at least for those characterized to date. All together the results advance our understanding of the structural determinants that govern the catalytic mechanism and substrate specificity in particular, and lay a foundation for future enzyme-engineering aimed at tailoring ASSTs to transfer sulfonate on acceptors of choice.

## ASSOCIATED CONTENT

### Supporting Information

**One PDF File (NimbonaNetal_SupportingInformation.pdf) that contains:**

Additional tables and figures of biochemical analyses; Additional figures on structural comparison; Computational methods; Additional tables, figures and discussion on docking, QM/MM calculations and umbrella sampling

## AUTHOR CONTRIBUTIONS

**Conceptualization:** M.C., F.D., M.D.D.

**Experimental methodology and investigation:** N.N, H.O., D.B., L.C., M.D.D., F.D., M.C.

**Computational methodology and software:** M.D.D., D.B.

**Data curation and validation:** N.N., L.C. (Biochemistry, Enzymology, Crystallography), H.O., A.J.D. (Enzymology), D.B. (Computational Investigation)

**Formal analysis:** N.N. (Biochemistry, Enzymology, Crystallography), H.O. A.J.D. (Enzymology), D.B. (Computational Investigation)

**Visualization:** N.N., H.O., A.J.D., D.B., M.C

**Writing – original draft:** N.N., M.C

**Writing – review & editing:** All authors

**Supervision:** M.C., L.C., M.D.D., F.D.

**Funding acquisition:** F.D., M.C., M.D.D.

## Notes

The authors declare that they have no known competing financial interests or personal relationships that would influence the work reported in this paper.

## ACKNOWLEDGMENT

This research was funded, in whole or in part, by the Agence Nationale de la Recherche (ANR), grant ANR-22-CE92-0060-01 and by the Région Pays de la Loire, grant Trajectoire Europe, and by the DFG (German Research Foundation), Project Number 505682627.

We are grateful to the Platform Metabomer and Station biologique de Roscoff (SBR), which is supported by EMBRC-France and the state funds provided by the French Ministry (MESR) for continued support (PLF172). We also thank Cédric Leroux for his valuable assistance and support throughout this work.

We are grateful to the NMR-EPR Platform in Brest and the Biogenouest platforms CrystalO at the Station Biologique de Roscoff for access and providing measurements. We also wish to thank Dr. Tristan Barbeyron and Dr. Diane Jouanneau for valuable scientific discussions.

We acknowledge SOLEIL for provision of synchrotron radiation facilities through MX BAGs 20201190 and 20220933 and we wish to warmly thank Pierre Legrand and Serena Sirigu for their technical assistance on the beamlines PX1, PX2 and Aurélien Thureau for assistance with SAXS measurements.

