## Supplemental Information, tables and figures for "Exploring the substrate specificity and catalytic mechanism in a PAPS-independent ASST from *Salmonella typhimurium*"

**ABBREVIATIONS**

ASST : arylsulfate sulfotransferase

CDD Search : conserved domain database

CV : collective variables

LC-MS/MS : Liquid Chromatography coupled to tandem Mass Spectrometry

MaGe : magnifying genomes

MES : 2-(N-morpholino)ethanesulfonic acid

NMR : Nuclear Magnetic Resonance

PAPS : 3'-phosphoadenosine 5'-phosphosulfate

PCS : *p*-cresyl sulfate

PDB : protein data bank

*p*NPS : *p*-nitrophenyl sulfate

*p*NP : *p*-nitrophenol

RMSD : root mean square deviation

SD : standard deviation

SDS-PAGE : Sodium Dodecyl Sulfate PolyAcrylamide Gel Electrophoresis

QM/MM : Quantum Mechanics/Molecular Mechanics

**Supplementary data**

1.Production of recombinant ASSTs

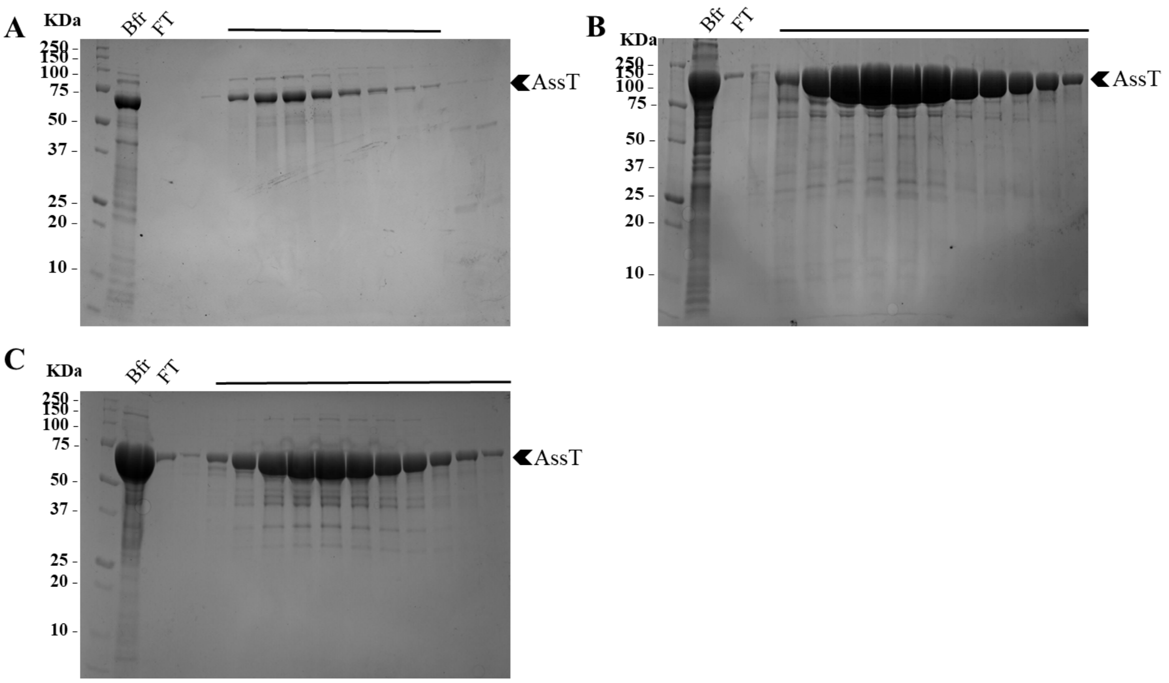


**Figure S1: SDS-PAGE analysis of recombinant ASSTs.** The figures are gels after a Size Exclusion Chromatography, (A) Ec_ASST_1 (Mw = 63.7 kDa), (B) St_ASST_5 (Mw = 63.8 kDa), and (C) Dh_ASST_6 (Mw = 74.3 kDa). The line above the gel indicates the fractions that were pooled.

| Construct | | Sequence |
|---|---|---|
| Ec_ASST_1 (Uniprot code Q8FDI4) | DNA | GCAGGTTTTAAACCGGCACCGCCTGCAGGTCAGCTGGGTGCAGTTATTGTTGATCCGT<br>ATGGTAATGCACCGCTGACCGCACTGGTTGATCTGGATAGCCATGTTATTAGTGATGTT<br>AAAGTTACGGTGCATGGCAAAGGTGAAAAAGGTGTTGAAATTAGCTATCCGGTTGGTCA<br>AGAAAGCCTGAAAACCTATGATGGTGTTCCGATTTTTGGCCTGTATCAGAAATTTGCCAA<br>TAAGGTTACCGTGGAATGGAAAGAAAATGGCAAAGTGATGAAAGATGATTACGTTGTTC<br>ATACCAGCGCCATCGTGAATAACTATATGGATAATCGTAGCATCAGCGATCTGCAGCAG<br>ACCAAAGTTATTAAAGTTGCACCGGGTTTTGAAGATCGTCTGTATCTGGTTAATACCCAT<br>ACCTTTACCGCACAGGGTAGTGATCTGCATTGGCATGGCGAAAAAGATAAAAATGCAGG<br>TATTCTGGATGCAGGTCCGGCAACCGGTGCACTGCCGTTTGATATTGCACCGTTTACCT<br>TTATTGTGGATACCGAAGGTGAATATCGTTGGTGGCTGGATCAGGATACCTTTTATGAT<br>GGTCGTGATCGCGATATTAACAAACGTGGTTATCTGATGGGTATTCGTGAAACACCGCG<br>TGGTACATTTACCGCAGTTCAGGGTCAGCATTGGTATGAATTTGATATGATGGGTCAAG<br>TGCTGGAAGATCATAAACTGCCTCGTGGTTTTGCAGATGCAACCCATGAAAGCATTGAA<br>ACCCCGAATGGCACCGTTCTGCTGCGTGTTGGTAAAAGCAATTATCGTCGTGATGATGG<br>CGTTCATGTTACCACCATTCGTGATCATATTCTGGAAGTTGATAAAAGCGGTCGTGTTGT<br>TGATGTTTGGGATCTGACCAAAATTCTGGATCCGAAACGTGATGCACTGCTGGGTGCCC<br>TGGATGCCGGTGCAGTTTGTGTTAATGTGGATCTGGCACATGCCGGTCAGCAGGCAAA<br>ACTGGAACCGGATACACCGTTTGGTGATGCATTAGGTGTTGGTCCGGGTCGTAATTGG<br>GCACATGTTAATAGCATTGCATATGATGCCAAAGACGATAGCATTATTCTGAGCAGCCG<br>TCATCAGGGTGTTGTTAAAATTGGTCGCGATAAACAGGTGAAATGGATTCTGGCACCGA<br>GCAAAGGTTGGGAAAAACCGCTGGCAAGCAAACTGCTGAAACCGGTTGATGCAAATGG<br>TAAACCGATTACCTGTAATGAAAATGGCCTGTGCGAAAACAGCGATTTCGATTTTACCTA<br>TACACAGCATACCGCATGGATTAGCAGCAAAGGCACCCTGACCATTTTTGATAATGGTG<br>ATGGTCGCCATCTGGAACAGCCAGCACTGCCGACCATGAAATATAGCCGTTTTGTGGAA<br>TACAAGATCGACGAAAAGAAAGGTACAGTTCAGCAGGTTTGGGAATATGGTAAAGAACG<br>CGGTTATGATTTCTATAGCCCGATTACCAGCATCATTGAATATCAGGCAGATCGCAATAC<br>CATGTTTGGTTTTGGTGGTAGCATCCACCTGTTTGATGTTGGTCAGCCGACCGTTGGTA<br>AACTGAATGAAATTGATTACAAAACGAAAGAAGTGAAAGTCGAGATTGATGTGCTGAGC<br>GATAAACCGAATCAGACCCATTATCGTGCCCTGCTGGTTCGTCCGCAGCAGATGTTTAA<br>A |
| | Protein | AGFKPAPPAGQLGAVIVDPYGNAPLTALVDLDSHVISDVKVTVHGKGEKGVEISYPVGQESL<br>KTYDGVPIFGLYQKFANKVTVEWKENGKVMKDDYVVHTSAIVNNYMDNRSISDLQQTKVIK<br>VAPGFEDRLYLVNTHTFTAQGXDLHWHGEKDKNAGILDAGPATGALPFDIAPFTFIVDTEGE<br>YRWWLDQDTFYDGRDRDINKRGYLMGIRETPRGTFTAVQGQHWYEFDMMGQVLEDHKLP<br>RGFADATHESIETPNGTVLLRVGKSNYRRDDGVHVTTIRDHILEVDKSGRVVDVWDLTKILD<br>PKRDALLGALDAGAVCVNVDLAHAGQQAKLEPDTPFGDALGVGPGRNWAHVNSIAYDAKD<br>DSIILSSRHQGVVKIGRDKQVKWILAPSKGWEKPLASKLLKPVDANGKPITCNENGLCENSD<br>FDFTYTQHTAWISSKGTLTIFDNGDGRHLEQPALPTMKYSRFVEYKIDEKKGTVQQVWEYG<br>KERGYDFYSPITSIIEYQADRNTMFGFGGSIHLFDVGQPTVGKLNEIDYKTKEVKVEIDVLSD<br>KPNQTHYRALLVRPQQMFK |
| St_ASST_5 (Uniprot code A0A0D6HM77) | DNA | ATGAATACACTAACGACAACTTCAGTTGTACTGCCGGCACCGCGTCCGGCGATAAACCA<br>AGGTATTGACATCAACAACGAGATGGTGTTAAACCACACCGCGATCTATGAGAACTGCC<br>TGGCGCAGGTTACGCAAGAGAATACCGTCGAGAACGCTCTGATGCTGCTCGACCCGTA<br>TGGCACCGCGCCACTGTCTGCCTATGCCGGTGTGTGGAGCCTTGAGCCGGCTGAAATC<br>ATCGTTACTGTTCAGGATGCGGCAAAGACTGCTATGCCTGTTGAGCACCTGTATACCTT<br>AACCGCTGGCGCGAACCTGTTGCCGGTTCTCGGTCTGGTAGCGGACACCGAAAATCGT<br>ATTGTGTTCAGCCAGGCGGACACGCCGCTCGCGGTGTACACGCTGATTACTCAACCGC<br>TGCCGCCGGTGGACAGTGCGGAAGTTGTGCTGGGTTTTCCGATTATCAACGTGACTCA<br>GCCTGCTACCGATGCAAATAAAATGGCACCGGGTTTCTACTTCATTACCCACTTCGACC<br>GTTATAACTACGCTCTGGACCAAAACGGCTTAGTACGCTGGTACGTTACCCAAGATTAC<br>CCGTCCTATAATTTTGTCCGTATCGATAACGGCCACTTTCTGACCACCAGCGAGGCGAA<br>GAATACCTATCTGGATATGTATGAATTCGATATGATGGGCCGTCTGCACACCTTCTACAA<br>CTTGGACAACCAATTTCATCATAGCATTTGGCCGTGGGATTCTAATACCATTGTGGCCC<br>CGAGCGAGTATACGTCCGGCCGTCCGGACGACCTGAAAACCAATGAGGATGGTGTTTC<br>CGTGGTGGATTTGACCACGGGTCTGGAAACCGCATACTATGATATGGCTAAGGTTTTGG<br>ATACCACGCGTGTTAGCAGACCAAGCGGTACTGCTCCGGGTGAGGACCCGACGGTGA<br>AAGATTGGCTGCATATTAACCAATCCTATGTGAACGAAACCAACCAGCTCCTGATCGCG<br>TCCGGCCGTCATCAGTCTGCGGTTTTTGGTGTGGACTTGCAGACCCAAGCGCTGCGCT<br>TCATTCTATCGACCCACGAAGACTGGGATGACGCGTACCAGCCGTACTTGTTGACCCC<br>GGTCGATTCTGAGGGTGTTGCGCTGTACGACTTTAGCAAGCAGGAAGACATCGATGCC<br>GCGGACCGCGATTTCTGGACCTGGGGTCAACATAATGTTGTAGAGATCGCAAATAACAC<br>GCCGGGTATTGTCGAGTTCATGGTTTTCGACAATGGCAACTACCGCTCACGCGATGATT<br>CGAAGAGCCTGCTTCCACCGGATAACTACAGCAGAATTGTGCACTTCGTTGTCAATATG<br>AATGAAATGACCGTTATGCGTCCGTTTGAATACGGTAAAGAACTGGGTGCGCGTGGTTA<br>CAGCAGCTGTGTTAGCGCGAAGGCGATCCAGCAGAACGGCAATATTGTGGTGCACTTT<br>GCAGATTGCACCTTTGACGAAAACGGCCGTGCCATCTCTTGCCAGCCGGGCGAAAGCG<br>ATATCATCGACCCGCAGGCAGGTAGCGAAGCGATGGGTTTGCTGATCCTGCAAGAGAT<br>CGCGCCGACAGAGAAAACCGTGTTGTTTGAGGCCACCATGACCAGCGGCTACTATAAA<br>AACGCCGAAACCAACGGTGAGGGCTATCGTTACGACATCACGTCGTTCCGCGTCTACA<br>AAATGGATTTGTACGCATGA |

| | | |
|---|---|---|
| | Protein | MNTLTTTSVVLPAPRPAINQGIDINNEMVLNHTAIYENCLAQVTQENTVENALMLLDPYGTAP<br>LSAYAGVWSLEPAEIIVTVQDAAKTAMPVEHLYTLTAGANLLPVLGLVADTENRIVFSQADTP<br>LAVYTLITQPLPPVDSAEVVLGFPIINVTQPATDANKMAPGFYFITHFDRYNYALDQNGLVRW<br>YVTQDYPSYNFVRIDNGHFLTTSEAKNTYLDMYEFDMMGRLHTFYNLDNQFHHSIWPWDS<br>NTIVAPSEYTSGRPDDLKTNEDGVSVVDLTTGLETAYYDMAKVLDTTRVSRPSGTAPGEDP<br>TVKDWLHINQSYVNETNQLLIASGRHQSAVFGVDLQTQALRFILSTHEDWDDAYQPYLLTPV<br>DSEGVALYDFSKQEDIDAADRDFWTWGQHNVVEIANNTPGIVEFMVFDNGNYRSRDDSKS<br>LLPPDNYSRIVHFVVNMNEMTVMRPFEYGKELGARGYSSCVSAKAIQQNGNIVVHFADCTF<br>DENGRAISCQPGESDIIDPQAGSEAMGLLILQEIAPTEKTVLFEATMTSGYYKNAETNGEGY<br>RYDITSFRVYKMDLYA |
| Dh_ASST_6 (Uniprot code B8FZF6) | DNA | ATGCGCACCTATCTGAATACCGAAAAACATCTGATTACCCTGCAGGCAGAAAGCGAAGA<br>ACGTTTTCTGGCAGAACTGCGTGCAGGTAATTATACCGCAGAAAGTCCGCTGGTTGTTA<br>AAAACCCGTATATCATTAATCCGCTGGCAGCCGTTATTTGCTTTAATACCGATGAAGAAA<br>CCACCGCTGAAATTACCGTTAAAGGTAAAGCAATTGAAGGCGATCTGAGCCATACCTTT<br>GCAGCAGCAAAAGAACATGTGCTGCCGGTTTATGGTCTGTATGATGATTATGTTAACAC<br>CGTGGTGATCAAACTGAGCAATGGTAAAACCAGCGAAGTGAAAATTGAAGTCGAAGAAC<br>TGAATGTGAACAAAGCCCTGTATTGTCGTACCACACCGGAATATTTTGGCAAAGATTTTA<br>TGCTGATTAGCACCACCACACCGCTGATTGAAAGCGCACGTACCGCAGGTTTTGATTAT<br>GCCGGTGATCTGCGTTGGTGTATTACCAATCTGCAGAGCTGGGATATTAAGAAACTGGA<br>AAATGGTCGTCTGCTGTATACCAGCCATCGTACCGTTCAGAAACCGTATTATAACGTTG<br>GTGTGATGGAAATGGATTTCTGCGGCAAAATCTATAAAGAATATCGTCTGCCTGGTGGC<br>TATCATCACGATGCAGTTGAATTAGAAAATGGCAATATTCTGGCAGCCAGCGATAACGA<br>TTTTAATGATAGCGTTGAAGATTTTGTGGTCGAAATTGAACGTGCAACCGGTGCAGTGAT<br>TAAAAGCTGGGATCTGCAGAAAATTCTGCCTCGTGGTCAGGGTAAAGCCGGTGATTGG<br>AATCATCATGATTGGTTTCATAATAACGCCGTGTGGTATGATAAACCGACCAATAGCATT<br>ACCATGAGCGGTCGTCATATGGATGCAGTGATCAATTTTGATTACGATAGCGGTGCACT<br>GAATTGGATTCTGGGTGATCCGGAAGGTTGGAGCGAAGAATGGCAGAAATACTTTTTCA<br>AAAACGTGACCAAAGGCGATTTCGATTGGCAGTATGAACAGCATGCAGCACGTATTCTG<br>CCGAATGGTGATGTGTTTCTGTTTGATAATGGCACCTATCGCAGCAAAAATGAAGCAAC<br>CCGTGTGGATCCGGAACAGAATTTTAGCCGTGGTGTTATTTATCGCATCGACACCGATA<br>AAATGGAAATCGAACAGGTTTGGCAGTACGGTAAAGAACGTGGTGCAGAATTCTATAGC<br>CCGTATATTTGCAACGTGGATTATTATGGTGAAGGCCATTATATGGTTCATAGCGGTGGT<br>ATTGCAACCTATCGTGGTAAACATACCGATGGTCTGGGTGCAATGCTGCTGAATAAATA<br>CAAAGATGAACACATCCATCTGACCCTGGAAAGCATTACCGTTGAAGTTCAGAATGATC<br>AGCTGAAATATGAGCTGAAAGTTCAAGGTGGTAACTATTATCGTGCACGTCGTGTGAGC<br>CCGTATGATGAGAAAACCAATCTGGTGTTAGGTAAAGGTGAACTGTTAGGTGGTTTTGG<br>TGTTACTCCGGAATTTATGAAGGTCAACTTTAAAGATGCAGAAACCGAACTGAGCGAAA<br>AACACAATCTGAATGTTATCCTGGAAGAAGATCGCCTGGCAATTCGTGCAAGCTTTCGT<br>GAAGGTAGCCAGGTTTTTCTGGAACTGAAAGGTGCAGAACAGAGCAAATTTTACAATAT<br>TCCGACCGAAGTTCATGATGTTACCGCAGCATGTGTTAGCTTTGAAGAACAGAACGATA<br>ACGACTTCCAGTTTTATGTTAGCCGTGAAGGTCTGAGCGGTGAATTTGGTATTTATCTGA<br>ACATTGACAGCAAACGCTATGATACCCATCTGAGCGTTAAACTG |
| | Protein | MRTYLNTEKHLITLQAESEERFLAELRAGNYTAESPLVVKNPYIINPLAAVICFNTDEETTAEIT<br>VKGKAIEGDLSHTFAAAKEHVLPVYGLYDDYVNTVVIKLSNGKTSEVKIEVEELNVNKALYCR<br>TTPEYFGKDFMLISTTTPLIESARTAGFDYAGDLRWCITNLQSWDIKKLENGRLLYTSHRTVQ<br>KPYYNVGVMEMDFCGKIYKEYRLPGGYHHDAVELENGNILAASDNDFNDSVEDFVVEIERA<br>TGAVIKSWDLQKILPRGQGKAGDWNHHDWFHNNAVWYDKPTNSITMSGRHMDAVINFDY<br>DSGALNWILGDPEGWSEEWQKYFFKNVTKGDFDWQYEQHAARILPNGDVFLFDNGTYRS<br>KNEATRVDPEQNFSRGVIYRIDTDKMEIEQVWQYGKERGAEFYSPYICNVDYYGEGHYMV<br>HSGGIATYRGKHTDGLGAMLLNKYKDEHIHLTLESITVEVQNDQLKYELKVQGGNYYRARR<br>VSPYDEKTNLVLGKGELLGGFGVTPEFMKVNFKDAETELSEKHNLNVILEEDRLAIRASFRE<br>GSQVFLELKGAEQSKFYNIPTEVHDVTAACVSFEEQNDNDFQFYVSREGLSGEFGIYLNIDS<br>KRYDTHLSVKL |

**Table S1 : DNA and protein sequences of constructs synthesized by Thermo Fisher (GenArt) and used in this study: Ec_ASST_1, St_ASST_5, and Dh_ASST_6.**

| | |
|---|---|
| ASST5_F174A_s | GTTTCTACTTCATTACCCACGCCGACCGTTATAACTACGCTC |
| ASST5_F174A_as | GAGCGTAGTTATAACGGTCGGCGTGGGTAATGAAGTAGAAAC |
| ASST5_Y557A_s | CAACGGTGAGGGCTATCGTGCCGACATCACGTCGTTC |
| ASST5_Y557A_as | GAACGACGTGATGTCGGCACGATAGCCCTCACCGTTG |
| ASST5_H401Q | CGATTTCTGGACCTGGGGTCAAGAGAATGTTGTAGAGATCGC |
| ASST5_H401Q_as | GCGATCTCTACAACATTCTCTTGACCCCAGGTCCAGAAATCG |

**Table S2 : Oligonucleotide primers used for site-directed mutagenesis of ASST enzymes.**

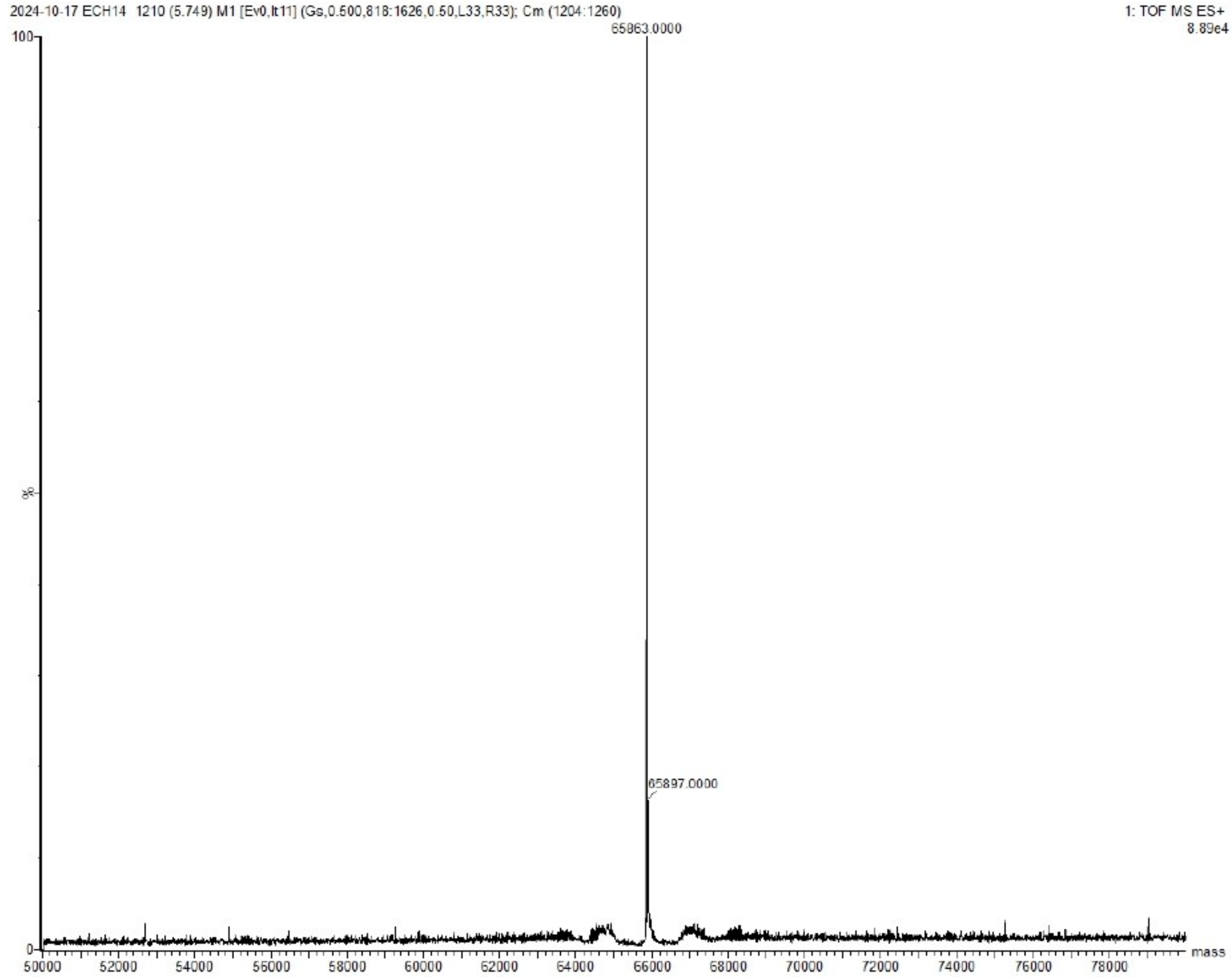


**Figure S2: UPLC–HRMS analysis of purified St_ASST_5 using an ESI(+) Q-TOF Synapt™ G2 system (Waters) after reversed-phase C18 separation**. The intact molecular mass was determined by spectral deconvolution (MaxEnt1). Theoretical mass: 65,994 Da; experimental mass: 65,863 Da. The observed 131 Da difference is consistent with the loss of the N-terminal methionine.

2. Biochemical characterization of St_ASST_5

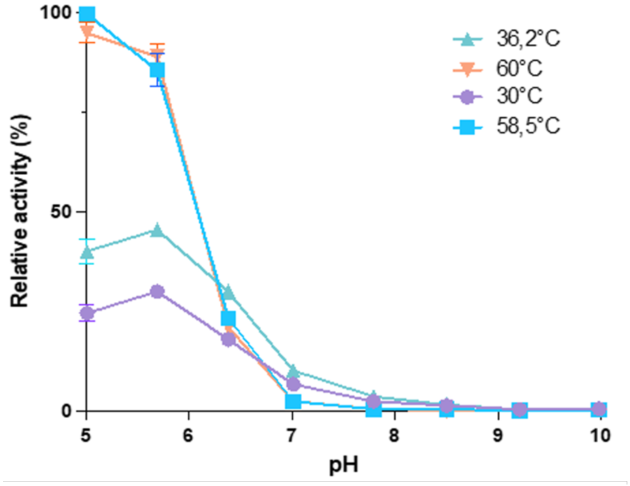


**Figure S3: Relative activity of St_ASST_5 as a function of the reaction pH at different temperatures (left) or of temperature at different pH (right)**. Reactions were carried out in universal Britton-Robinson buffer (75 mM) using p-nitrophenyl sulfate (10 mM) as the donor substrate and phenol (20 mM) as the acceptor substrate.

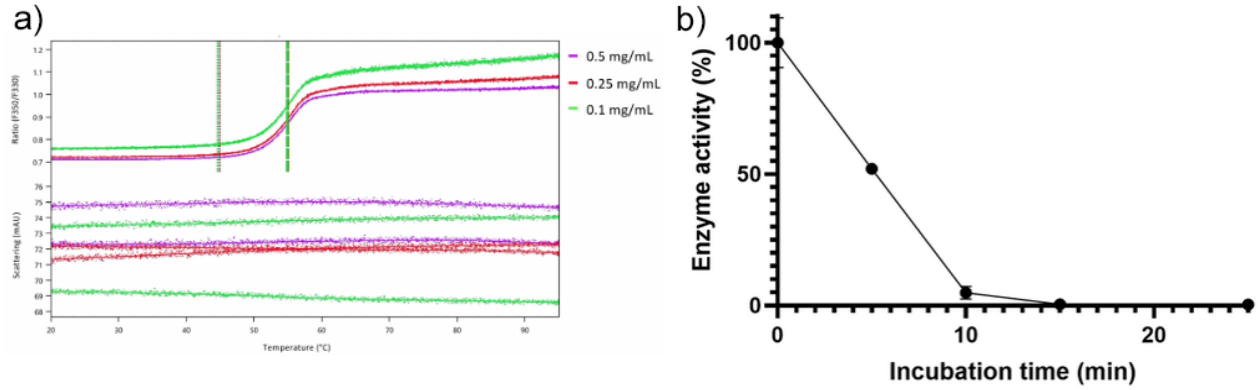


**Figure S4: Thermal stability of St_ASST_5 analyzed by NanoDSF** (**a**): purified St_ASST_5 was loaded into glass capillaries and heated from 20 to 90°C at 1°C/min. Fluorescence ratios ($F_{350}/F_{330}$) were monitored to determine the melting temperature (Tm) onset (for ratio) of denaturation, static light scattering was recorded to detect aggregation. Experiments were performed at three concentrations (0.1, 0.5 and 1.0 mg/mL); residual enzymatic activity of St_ASST_5 at 58°C (**b**): the enzymatic activity was monitored over time after pre-incubation at 58°C in 50 mM MES buffer (pH 5.5). Activity is expressed relative to a non-heated control (100% activity).

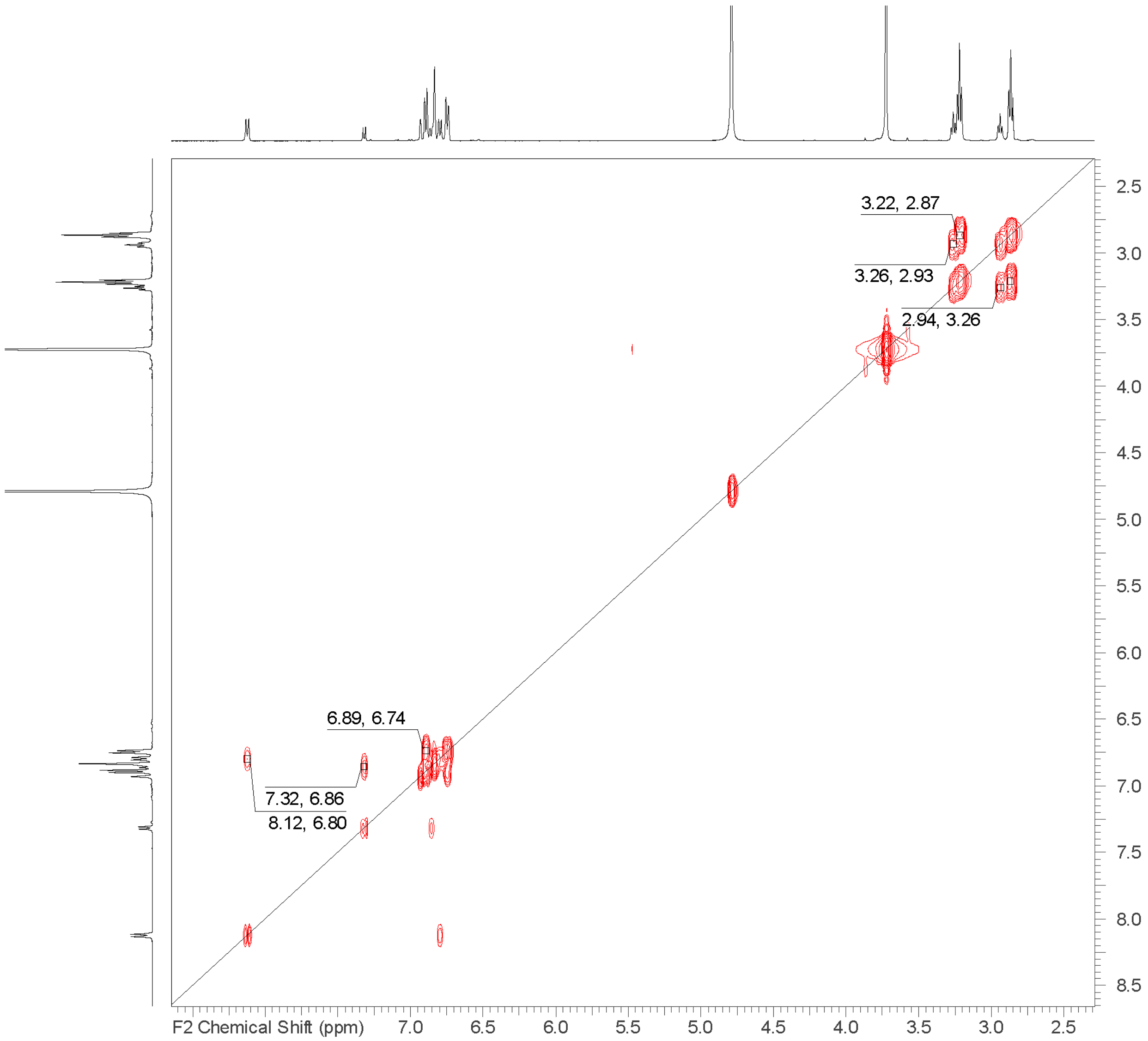


**Fig S5. 2D-NMR spectrum of reaction product of St_ASST_5**. $^{1}H$-$^{1}H$ COSY spectrum of the reaction products after activity of St_ASST_5 on *p*NPS and dopamine, showing the signals of the aromatic protons in the region 6.7-8.2 ppm. The chemical shift depends on their distance to the deshielding O-$SO_3^-$ group, allowing to assign the product being 4O sulfated dopamine.

The COSY spectrum shows correlation between signals at 8.12 and 6.80 ppm, allowing the attribution of signals from p-NP. It also shows correlation between signals at 7.32 and 6.86 ppm, allowing the attribution of aromatic protons from dopamine-sulfate. The deshielding of H5 being more pronounced than H2, it clearly shows the sulfatation has occurred on position 4 and not 3.

3. Comparative genomics

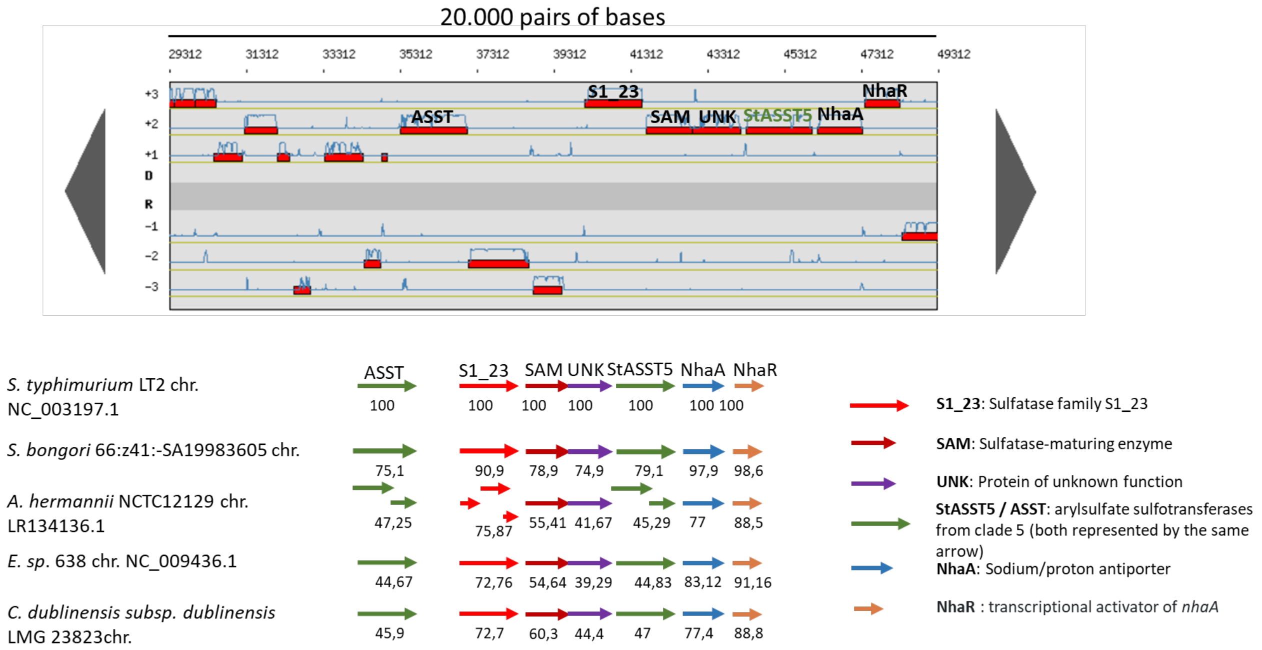


**Figure S6: Genomic organization of the ST_ASST_5 gene cluster viewed on MaGe.** Comparative genomic analysis of a ~20 kb region surrounding the St_ASST_5 gene in *Salmonella enterica* subsp. *enterica* serovar Typhimurium 14028S and other selected bacterial species. The *Salmonella typhimurium* LT2 chromosome is included as an additional reference to assess the conservation of the gene cluster across closely related strains and more distant species. Genes are represented as arrows indicating transcriptional orientation. Conserved gene clusters include genes encoding a sulfatase (S1_23), a sulfatase-maturating enzyme (SAM), and arylsulfate sulfotransferases (StASST5 / ASST). The latter correspond to clade 5 ASSTs and are represented by the same arrow. Additional annotated genes include UNK (protein of unknown function), NhaA (sodium/proton antiporter), and NhaR (transcriptional activator of *nhaA*). Numbers below each gene indicate the percentage of sequence identity relative to the corresponding gene in *S. enterica* serovar Typhimurium 14028S.

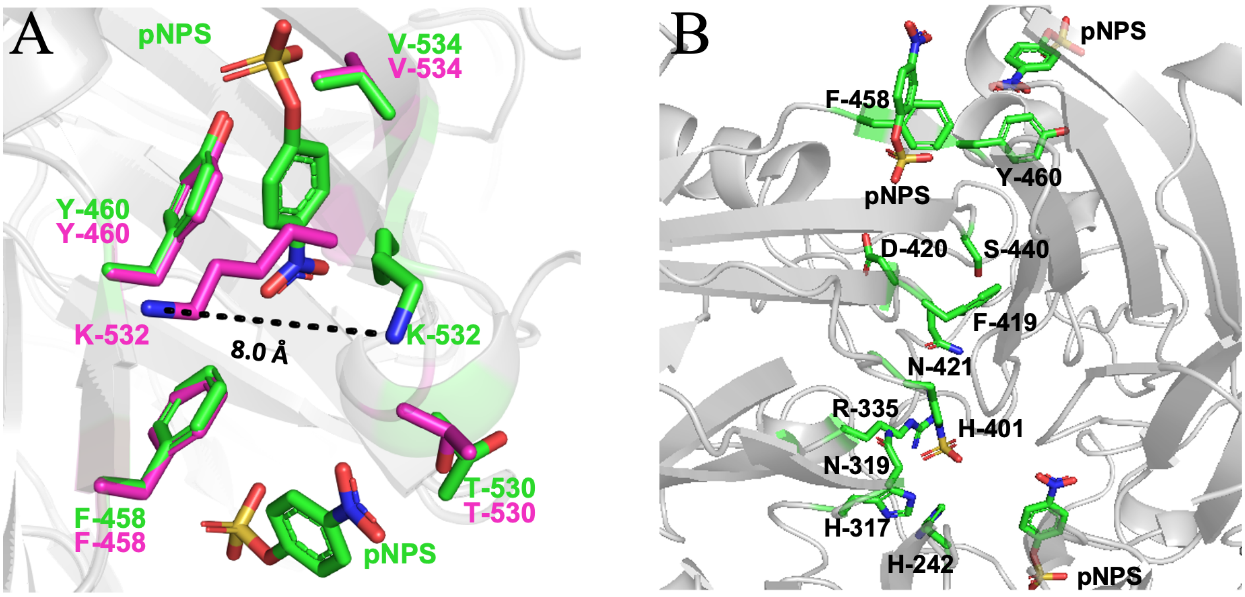


**Figure S7: Structural differences between St_ASST_5_nat and St_ASST_5_pNPS and identification of a catalytic tunnel**. A: Comparison between the native (pink) and complex with pNPS (green) structures of St_ASST_5 showing a loop displacement spanning residues Val-534 to Thr-530, with the largest shift observed at Lys-532. B: Two pNPS molecules, stabilized by interactions with aromatic residues (Phe-458 and Tyr-460), reveal the presence of a tunnel formed by residues conserved (showed in green sticks) in more than 80% of ASSTs.

4. Docking pose selection for phenol and dopamine in St_ASST_5

Docking was used to generate catalytically competent starting structures for the phenol- and dopamine-bound St_ASST_5 systems used in subsequent MD and QM/MM simulations using MOE (1). The docking pocket was defined around the active-site residues His242, His317, Asn319, Arg335, His401, Cys472, Val473, and Phe562. In total, 3000 docking poses were generated for each substrate. Docking scores are reported in kcal·mol$^{-1}$. Final pose selection was based primarily on catalytic geometry rather than docking score alone. A pose was considered catalytically competent when the acceptor hydroxyl group was oriented toward the sulfate sulfur of the sulfonated His401 intermediate and the hydroxyl proton was positioned toward His317, the proposed catalytic base, according to the sulfo-transfer mechanism previously described for the homologous Dh_ASST_6 system (2).

| Simulation/MOE residue | Manuscript residue | Role |
|---|---|---|
| **HIE235** | His242 | Active-site histidine |
| **HID310** | His317 | Catalytic base |

| | | |
|---|---|---|
| **ASN312** | Asn319 | Active-site polar residue |
| **ARG328** | Arg335 | Active-site charged residue |
| **HIX/HIS394** | His401 | Sulfonated catalytic histidine |
| **CYS465** | Cys472 | Active-site residue |
| **VAL466** | Val473 | Active-site residue |
| **PHE555** | Phe562 | Aromatic pocket residue |

**Table S3. Residue numbering used for docking analysis.**

Phenol docking was straightforward because phenol contains only one reactive hydroxyl group. The selected pose placed the phenolic hydroxyl group between His317 and the sulfonated His401 intermediate. The phenolic oxygen was oriented toward the sulfate sulfur, while the hydroxyl proton pointed toward His317. This pose was therefore used as the catalytically competent starting structure for phenol MD and QM/MM simulations.

Dopamine docking was more ambiguous because dopamine contains two catechol hydroxyl groups and an amino group. Therefore, several docking poses were inspected manually for their catalytic relevance.

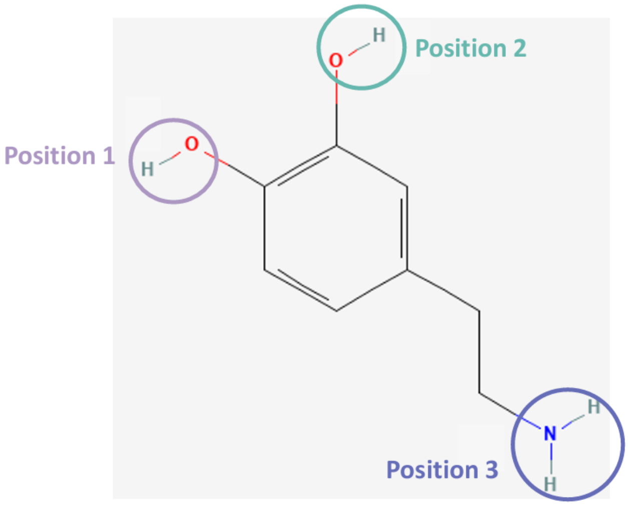


**Figure S8. Chemical structure of dopamine highlighting the three possible sulfation sites considered for regioselective sulfo transfer.** Position 1 corresponds to the catechol hydroxyl group that adopted the catalytically competent orientation. Position 2 indicates the second catechol hydroxyl group, whereas Position 3 corresponds to the terminal amino group.

Poses in which the amino group was oriented toward the catalytic histidines, or in which both catechol hydroxyl groups were directed away from the sulfate sulfur, were considered nonproductive for O-sulfation. The selected dopamine pose instead placed one catechol hydroxyl group (Position 1 according to Figure S8) toward the sulfate sulfur of the sulfonated His401 intermediate and positioned the corresponding hydroxyl proton toward His317.

| **Substrate/ pose** | **Docking Score kcal·mol⁻¹** | $O_{acceptor}$–$S_{sulfate}$ **Å** | $H_{acceptor}$–$N_{His317}$ **Å** | **Interpretation** |
|---|---|---|---|---|
| **Phenol Selected pose** | -3.2345 | 3.563 | 2.123 | Catalytically competent |
| **Dopamine selected pose** | -4.3187 | 3.549 | 2.681 | Catalytically competent O-sulfation pose |
| **Dopamine alternative pose 1** | -4.2332 | 3.8702 | 2.7237 | Catechol hydroxyl not optimally aligned |
| **Dopamine alternative pose 2** | -4.0494 | 4.052 | 3.083 | Amino group oriented toward catalytic histidine |

**Table S4. Geometric criteria for selected and alternative docking poses.**

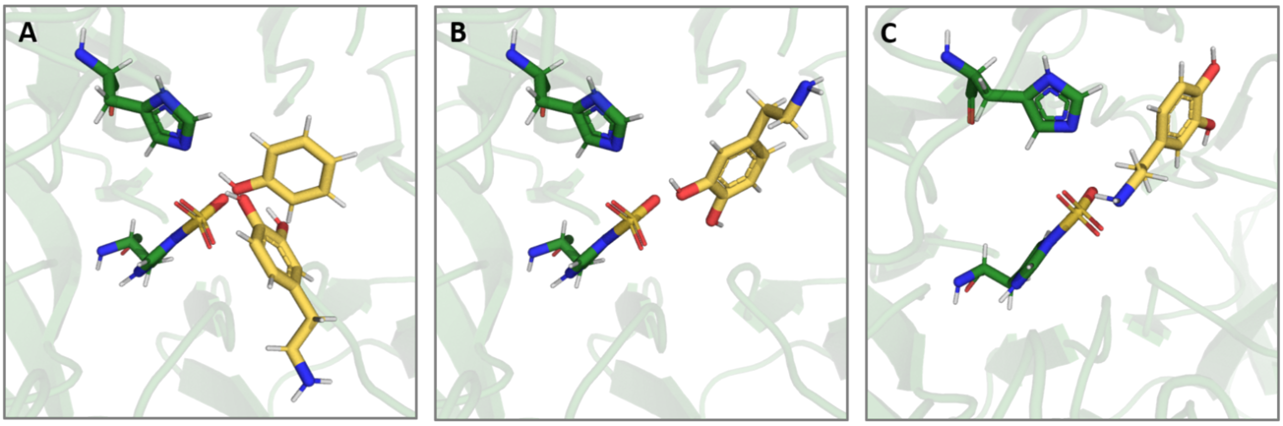


**Figure S9. Representative docking poses of dopamine in the active site of St_ASST_5.** (a) Selected catalytically competent dopamine docking pose used for subsequent simulations, shown together with the final phenol docking pose for comparison. In this orientation, the catechol hydroxyl group at position 1 is directed toward the His401-bound sulfo-group and the catalytic His317 residue. (b) Alternative dopamine docking pose in which the second catechol hydroxyl group is oriented toward the catalytic region. (c) Alternative dopamine docking pose in which the terminal amino group points toward the His401/His317 catalytic region.

The productive dopamine docking pose, was selected because it combined the most favorable docking score with the required catalytic geometry. The catechol hydroxyl group assigned as Position 1 was positioned between His317 and the sulfonated His401 intermediate. The corresponding catechol oxygen was oriented toward the sulfate sulfur, while the hydroxyl proton pointed toward the catalytic nitrogen of His317, fulfilling the geometric requirements for His317-mediated deprotonation and subsequent S–O bond formation. In addition to these catalytic contacts, the pose was stabilized by the surrounding active-site environment. His242 was positioned close to the catechol moiety and may contribute to substrate alignment. Cys472, Val473, and Phe174 form a confined pocket around the aromatic ring and help maintain the catechol group in the productive orientation. The aminoethyl substituent is directed away from the catalytic His317/His401 region and is stabilized toward the pocket entrance, including a polar contact with Asp490. This interaction pattern anchors dopamine without allowing the amino group to compete with the reactive catechol hydroxyl group for the catalytic region.

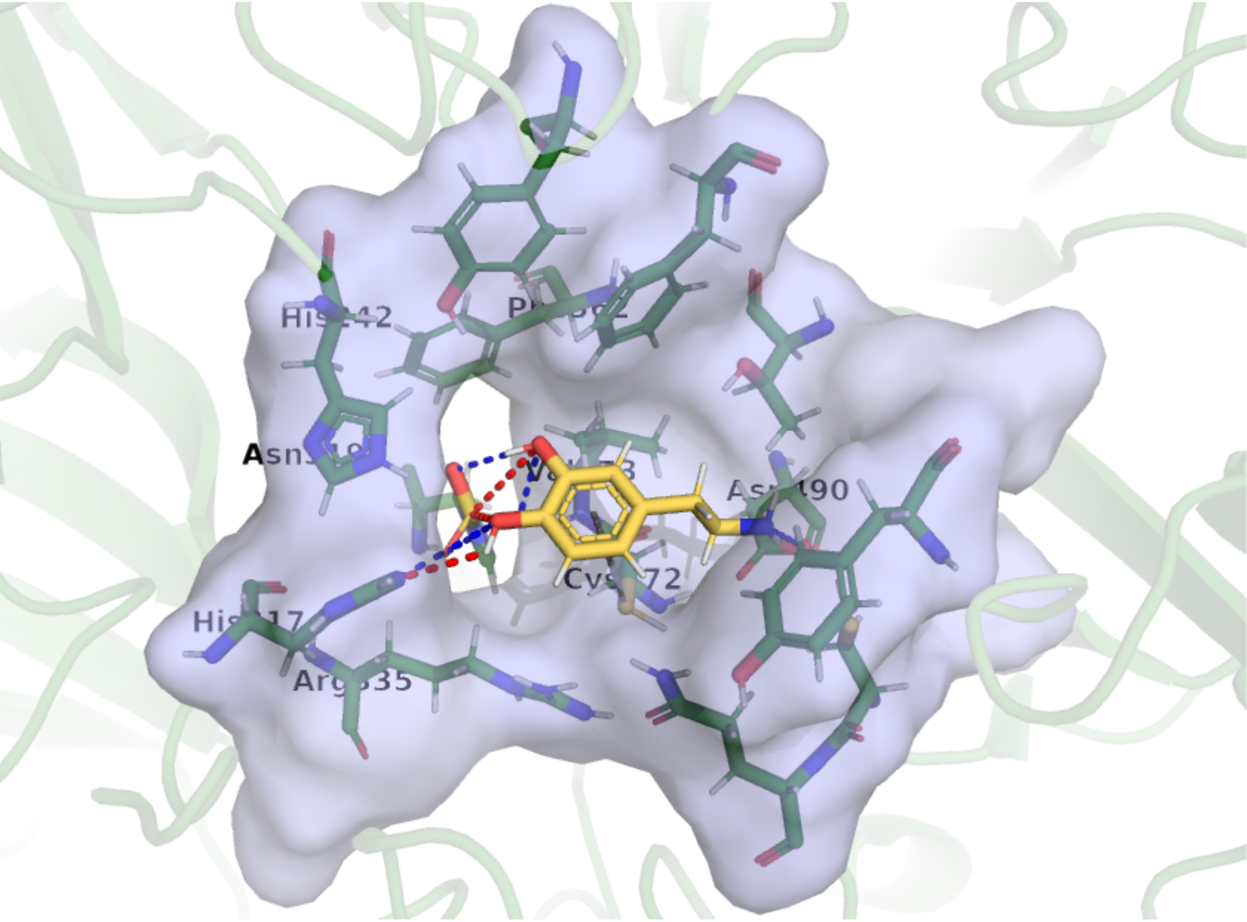

**Figure S10. Stabilization of the catalytically competent dopamine docking pose in the St_ASST_5 active-site pocket.** The selected best-scoring dopamine docking pose is shown in yellow sticks within the active-site pocket of St_ASST_5. The pocket surface is shown in transparent light blue, and surrounding active-site residues are shown as sticks. Dashed lines indicate key stabilizing contacts between dopamine and the pocket, including the catalytic positioning of the reactive catechol hydroxyl group toward the sulfonated His401 intermediate and His317, as well as additional interactions with surrounding residues. Cys472 is positioned close to the catechol ring and may contribute to stabilization through an S···π-type interaction.

The alternative pose places the second catechol hydroxyl group toward the catalytic region and therefore represents the alternative regioisomeric orientation. Although this pose retains active-site contacts, it does not correspond to the selected productive Position 1 hydroxyl group. The position, where the amino group is oriented toward the His401/His317 catalytic region and forms favorable polar contacts with the sulfonated His401/Arg335 region. However, in this orientation the catechol hydroxyl groups are displaced from the sulfate sulfur and His317, making the pose nonproductive for O-sulfonation. Thus, the comparison of dopamine docking poses shows that favorable binding interactions alone are insufficient; productive catalysis requires simultaneous stabilization of the catechol ring, anchoring of the aminoethyl substituent away from the catalytic center, and correct positioning of the reactive catechol hydroxyl group between His317 and the His401-bound sulfo-group.

| **Pose** | **Orientation** | **Stabilizing contacts** | **Interpretation** |
|---|---|---|---|
| **Position 1** | Position 1 catechol OH toward His401/His317 | His401 sulfo group, His317, His242, Cys472, Val473, Phe174, Asp490 | Best-scoring and catalytically competent |
| **Position 2** | Alternative catechol OH toward catalytic region | Active-site contacts retained, but wrong catechol OH selected | Alternative regioisomeric orientation |
| **Position 3** | Amino group toward catalytic region | Sulfo-His401/Arg335/His317 contacts stabilize amino group | Stable but nonproductive for fation |

**Table S5: Docking-pose comparison of dopamine binding orientations and stabilizing active-site interactions in St_ASST_5.**

6. CAVER analysis of substrate access pathways in St_ASST_5

Substrate access pathways in the selected phenol- and dopamine-bound St_ASST_5 structures were analyzed with the CAVER 3.0 (3) plugin in PyMOL (4). Calculations were performed with a probe radius of 0.6 Å, shell radius of 3.0 Å, shell depth of 4.0 Å, average-link clustering, and a clustering threshold of 3.5. Tunnel-lining residues were defined as residues within 3.0 Å of the tunnel. For dopamine, four detected tunnels were clustered into two final tunnel clusters, and the cheapest tunnel per cluster was retained for analysis.

Residues are reported using manuscript numbering. Thus, residue 394 in the CAVER output corresponds to His401 in the manuscript.

| Substrate | Tunnel | Throughput | Bottleneck radius / Å | Length / Å | Curvature |
|---|---|---|---|---|---|
| **Phenol** | Main tunnel | 0.42 | 0.74 | 6.64 | 1.28 |
| **Dopamine** | Cluster 1 | 0.775 | 0.859 | 3.295 | 1.103 |
| **Dopamine** | Cluster 2 | 0.260 | 0.609 | 8.661 | 1.153 |

**Table S6. Tunnel characteristics of phenol and dopamine access pathways.**

The dopamine CAVER output identifies cluster 1 as the more accessible pathway, with a higher throughput, larger bottleneck radius, and shorter length than cluster 2.

| Substrate | Tunnel | Tunnel-lining residues |
|---|---|---|
| **Phenol** | Main tunnel | Phe174, Tyr198, His242, His317, Asn319, Arg335, His401, Cys472, Val473, Phe562, phenol |
| **Dopamine** | Cluster 1 | His317, Arg335, His336, His401, Cys472, Val473, Asp490, Gln503, Tyr557, dopamine |
| **Dopamine** | Cluster 2 | Arg335, His401, Ser471, Cys472, Val473, Asp490, Cys502, Gln503, Pro504, Arg556, Tyr557, dopamine |

**Table S7. Tunnel-lining residues**

For phenol, the single main pathway is narrow and lined by residues from the catalytic pocket, consistent with a confined binding environment. In contrast, dopamine shows two possible access pathways. The main dopamine tunnel is shorter and wider than the phenol tunnel, whereas the second dopamine tunnel is longer and narrower. This supports the interpretation that dopamine can occupy a more flexible active-site access environment, while productive catalysis still requires correct positioning of one catechol hydroxyl group toward the sulfonated His401 intermediate.

5. Structure comparisons

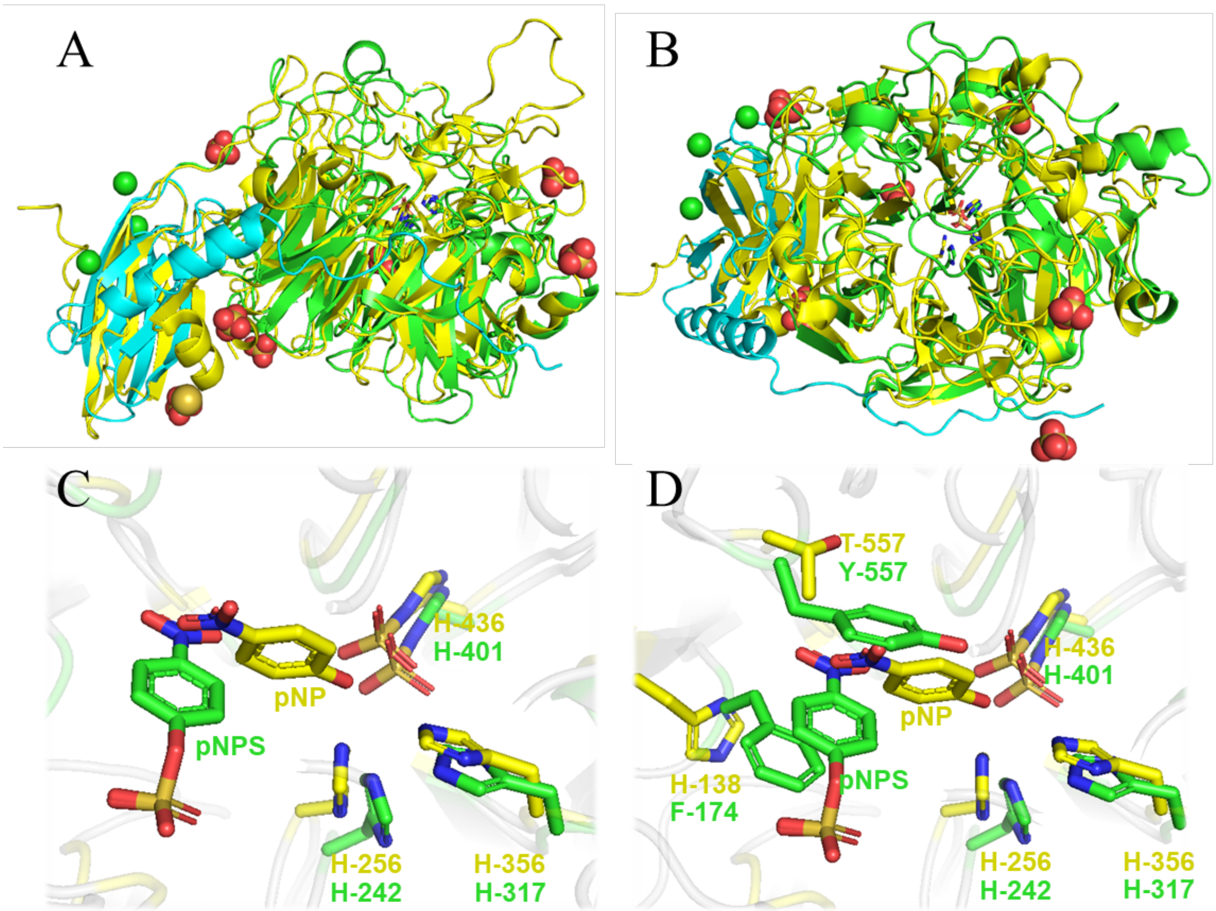


**Figure S11: Structural comparison between St_ASST_5 and Ec_ASST_1**. St_ASST_5 is shown in green and EC_ASST_1 in yellow. (**a**) Side view of the superposition of St_ASST_5 and Ec_ASST_1. (**b**) View from the top, into the catalytic site of the superposition of St_ASST_5 and Ec_ASST_1. (**c**) Enzyme-sulfate bound intermediate structures of the active site showing *p*NPS in the ST_ASST_5 and *p*NP in the EC_ASST_1. (**d**) Close-up view of residues involved in substrate binding. In St_ASST_5, Phe174 (F-174) and Tyr557 (Y-557) participate in substrate recognition, while in Ec_ASST_1 residues His138 (H-138) and Thr557 (T-557) occupy approximately the same positions next to the catalytic site.

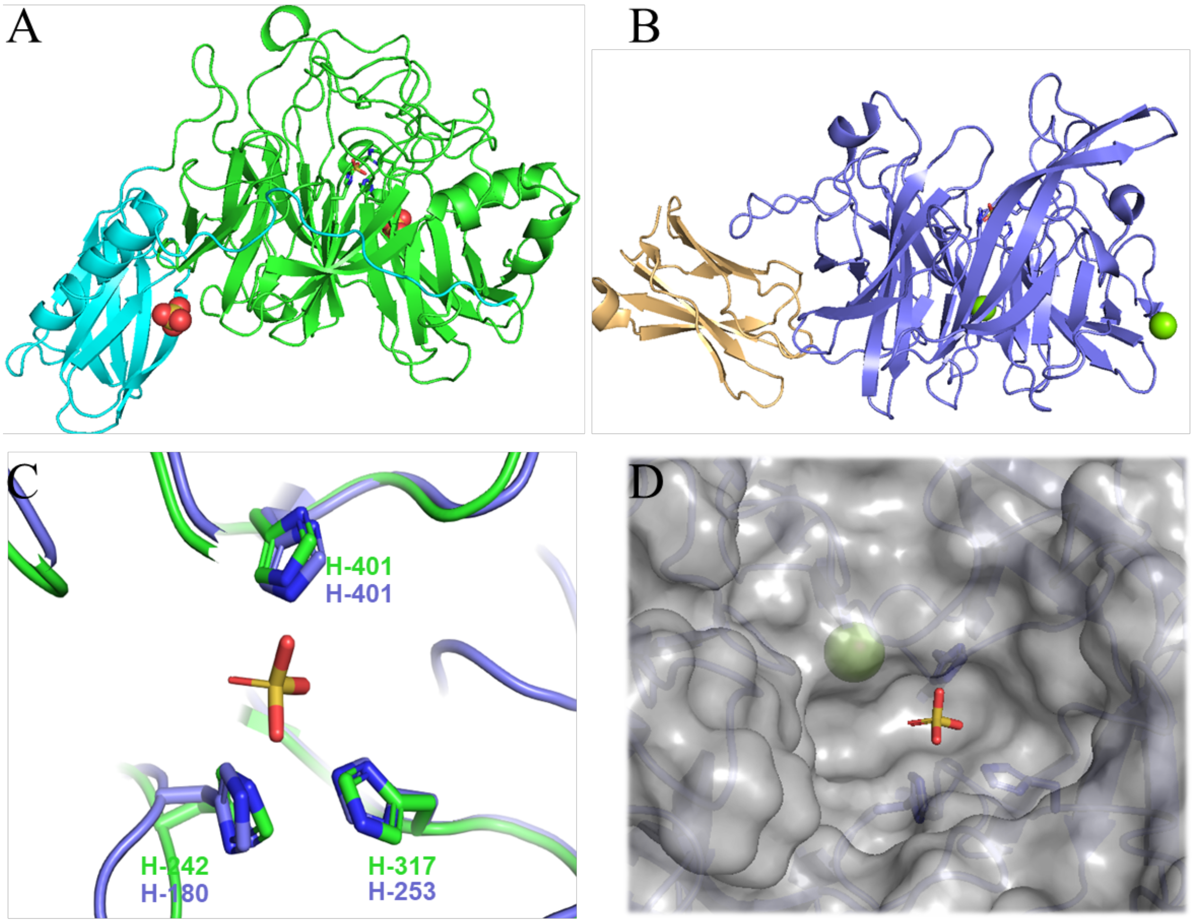


**Figure S12: Structural comparison between St_ASST_5 and CPZ4**. (A) Overall structure of St_ASST_5 shown in green the catalytic site and in blue the additional domain. (B) Structure of CPZ4 (accession code 9KF6) the catalytic domain shown in purple and the additional domain in wheat. (C) Superposition of the active sites highlighting conserved histidine residues involved in catalysis. Residues from St_ASST_5 are shown in green and those from CPZ4 in purple. (D) Surface representation of the CPZ4 active site pocket illustrating how wide and open it is (more than 20 Å wide).

#### 7. Classical MD analysis of phenol- and dopamine-bound St_ASST_5

Classical MD simulations performed with AMBER22 using the ff19SB force field were analyzed with cpptraj and MDAnalysis to assess the stability of the selected phenol- and dopamine-bound St_ASST_5 complexes and to identify catalytically relevant conformations for subsequent QM/MM simulations (5-7). For each substrate, three independent trajectories were combined after backbone alignment, giving 6000 sampled frames per system.

##### 7.1 Backbone stability

The backbone RMSD values (determined with 8) showed that both substrate-bound systems remained structurally stable during the simulations. For the phenol-bound system, the backbone RMSD fluctuated mainly between approximately 1.2 and 2.1 Å after initial relaxation. The dopamine-bound system showed slightly lower overall backbone deviations, with RMSD values mainly between approximately 0.9 and 1.7 Å. No major global rearrangement or unfolding of St_ASST_5 was observed in either system.

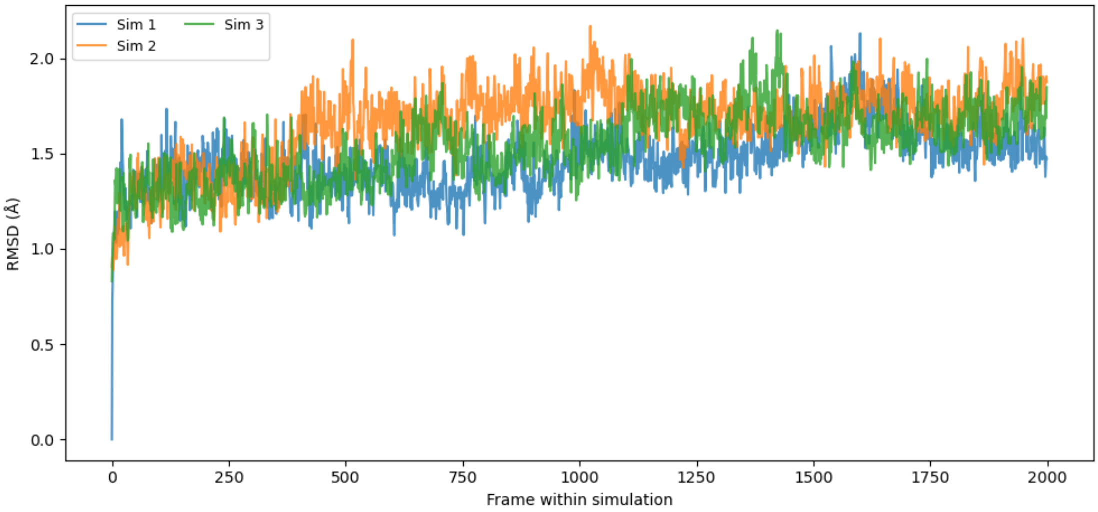


**Figure S13. Backbone RMSD of the phenol-bound St_ASST_5 simulations**. Backbone RMSD values for the three independent phenol-bound MD simulations after alignment to the protein backbone. First run is shown in blue, second run is shown in orange, third run is shown in green.

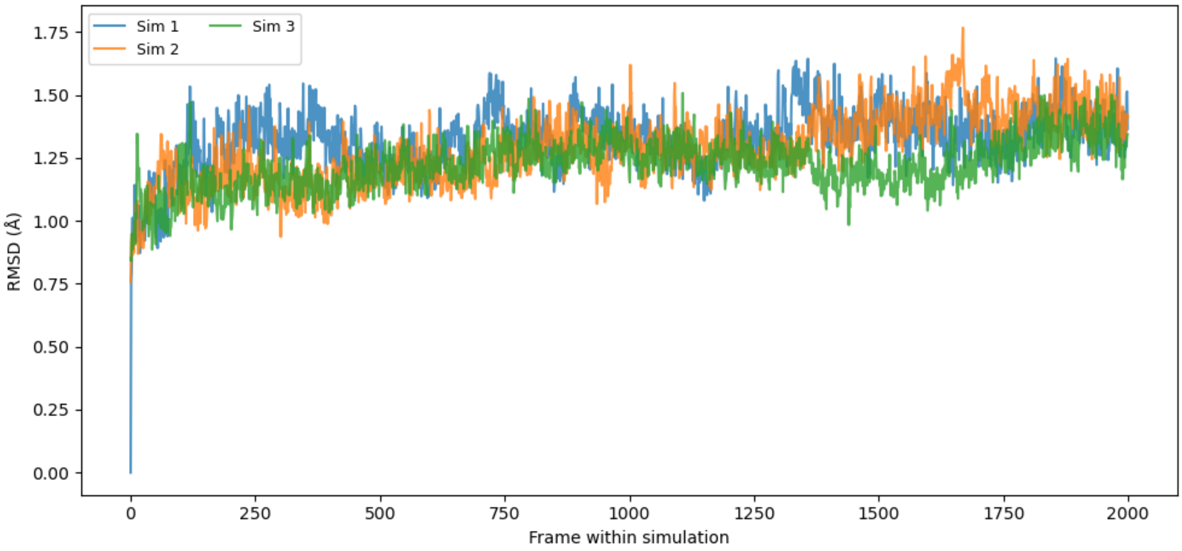


**Figure S14. Backbone RMSD of the dopamine-bound St_ASST_5 simulations**. Backbone RMSD values for the three independent dopamine-bound MD simulations after alignment to the protein backbone. First run is shown in blue, second run is shown in orange, third run is shown in green.

#### 7.2 Conformational clustering

The combined aligned trajectories were clustered based on protein backbone coordinates to characterize the conformational variability sampled during the MD simulations (8). For phenol, five clusters were identified, whereas four clusters were identified for dopamine. The all-versus-all RMSD heatmaps show defined cluster blocks for both systems, indicating that the trajectories sample several related backbone conformational states. For phenol, the RMSD values between cluster representatives ranged from 1.15 to 1.45 Å. For dopamine, the representative structures were slightly more similar, with inter-cluster RMSD values ranging from 1.11 to 1.34 Å. Thus, both systems retained the same overall fold, while sampling moderate conformational variability around the active site.

| System | Cluster | Representative frame | Cluster size |
|---|---|---|---|
| **Phenol** | C1 | 5889 | 245 |
| **Phenol** | C2 | 5056 | 1618 |
| **Phenol** | C3 | 3358 | 1484 |
| **Phenol** | C4 | 1819 | 1082 |
| **Phenol** | C5 | 2137 | 1571 |
| **Dopamine** | C1 | 3987 | 486 |
| **Dopamine** | C2 | 1563 | 772 |
| **Dopamine** | C3 | 2928 | 816 |
| **Dopamine** | C4 | 4398 | 3926 |

**Table S8. Cluster representatives from the combined MD trajectories.**

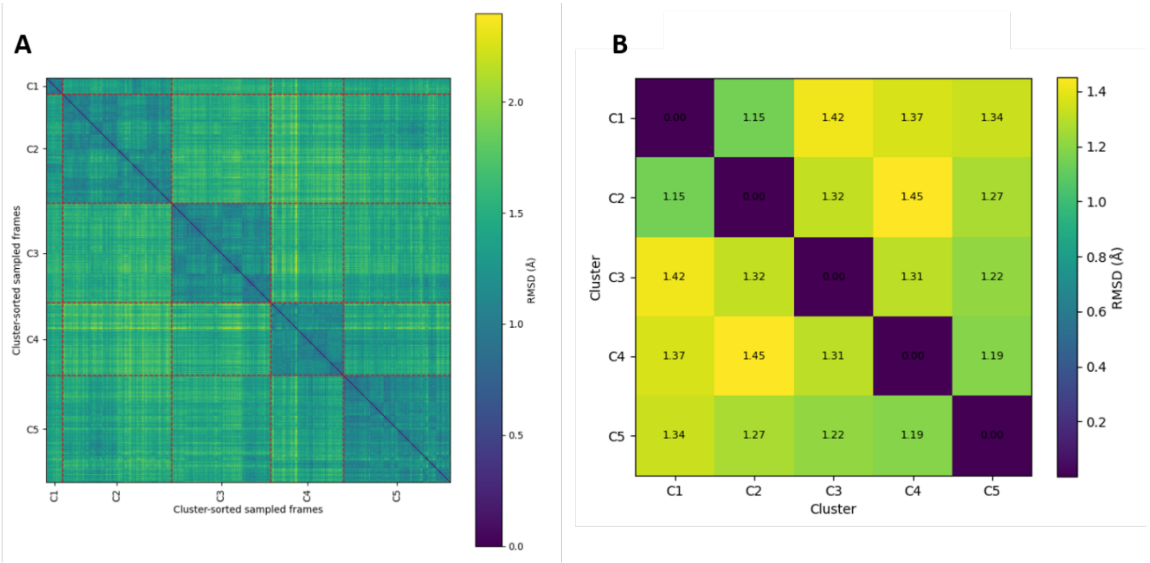


**Figure S15. RMSD Heatmaps of phenol-bound simulations.** (A) All-versus-all backbone RMSD heatmap of the phenol-bound trajectory. Frames were sorted by cluster assignment. Red dashed lines indicate cluster boundaries. (B) Backbone RMSD values between representative structures of the five phenol-bound clusters.

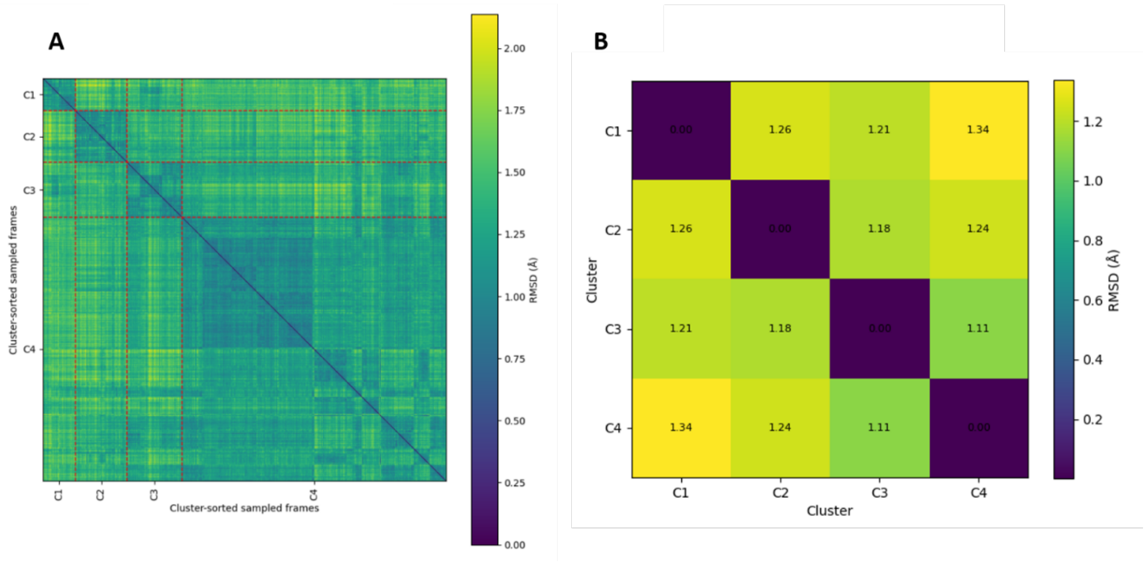


**Figure S16. RMSD Heatmaps of dopamine-bound simulations.** (A) All-versus-all backbone RMSD heatmap of the dopamine-bound trajectory. Frames were sorted by cluster assignment. Red dashed lines indicate cluster boundaries. (B) Backbone RMSD values between representative structures of the four dopamine-bound clusters.

##### 7.3 PCA of the combined trajectories

Principal component analysis further supported the clustering results (8). For phenol, the first two principal components explained 15.0% and 12.1% of the variance, respectively. For dopamine, PC1 and PC2 explained 15.2% and 9.3% of the variance. In both systems, the cluster representatives occupy distinct regions of the projected conformational space, indicating that the MD simulations sample multiple related conformational substates.

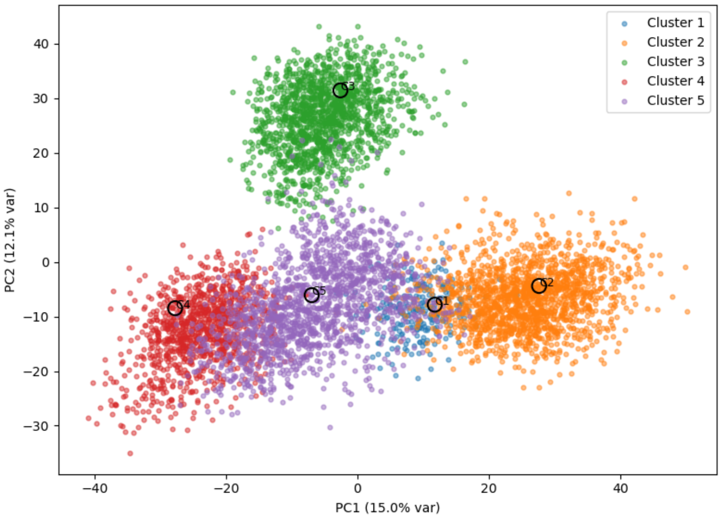


**Figure S17. PCA projection of the phenol-bound trajectory**. Projection of the combined phenol-bound MD trajectory onto the first two principal components. Cluster representatives are shown as black circles.

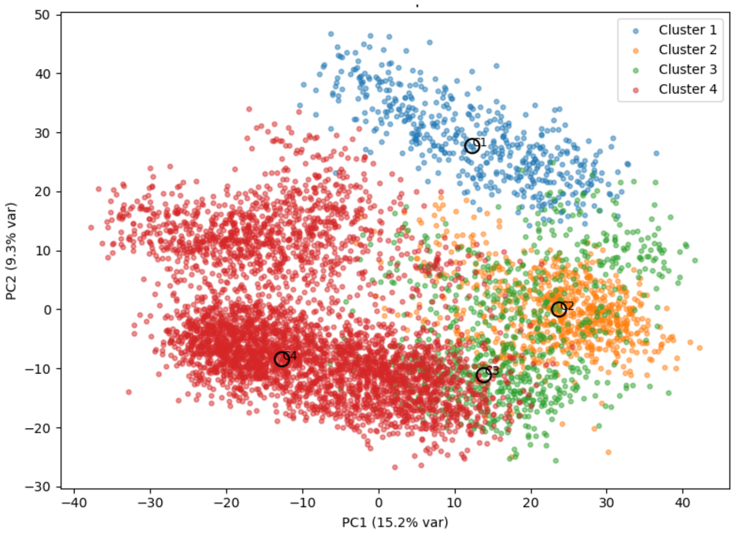


**Figure S18. PCA projection of the dopamine-bound trajectory**. Projection of the combined dopamine-bound MD trajectory onto the first two principal components. Cluster representatives are shown as black circles.

7.4 Catalytic distance analysis

To identify catalytically relevant frames, two distances were monitored for each substrate: the distance between the substrate hydroxyl proton and His317, and the distance between the substrate oxygen and the sulfate sulfur of the sulfonated His401 intermediate.

| **System** | **Distance** | **Atom pair** | **Interpretation** |
| --- | --- | --- | --- |
| **Phenol** | $H_{phenol}–N_{His317}$ | 8685–4765 | Proton transfer geometry |

| | | | |
|---|---|---|---|
| **Phenol** | $O_{phenol}$–$S_{sulfate}$ | 8684–6074 | S–O bond-forming geometry |
| **Dopamine** | $H_{dopamine}$–$N_{His317}$ | 8701–4765 | Proton transfer geometry |
| **Dopamine** | $O_{dopamine}$–$S_{sulfate}$ | 8700–6074 | S–O bond-forming geometry |

**Table S9. Atom pairs used for catalytic distance monitoring**

For phenol, frames were selected when both conditions were fulfilled: $H_{phenol}$–$N_{His317}$< 4 Å and $O_{phenol}$–$S_{sulfate}$< 6 Å. Using these criteria, 433 of 6000 frames fulfilled both conditions, corresponding to 7.22% of the phenol ensemble.

For dopamine, frames were selected when both conditions were fulfilled: $H_{dopamine}$–$N_{His317}$ < 6 Å and $O_{dopamine}$–$S_{sulfate}$ < 4 Å. Using these criteria, 1367 of 6000 frames fulfilled both conditions, corresponding to 22.78% of the dopamine ensemble.

| System | H–N cutoff / Å | O–S cutoff / Å | Frames satisfying both criteria | Fraction of ensemble |
|---|---|---|---|---|
| **Phenol** | < 4 | < 6 | 433 / 6000 | 7.22% |
| **Dopamine** | < 6 | < 4 | 1367 / 6000 | 22.78% |

**Table S10. Catalytically relevant frames identified from MD**

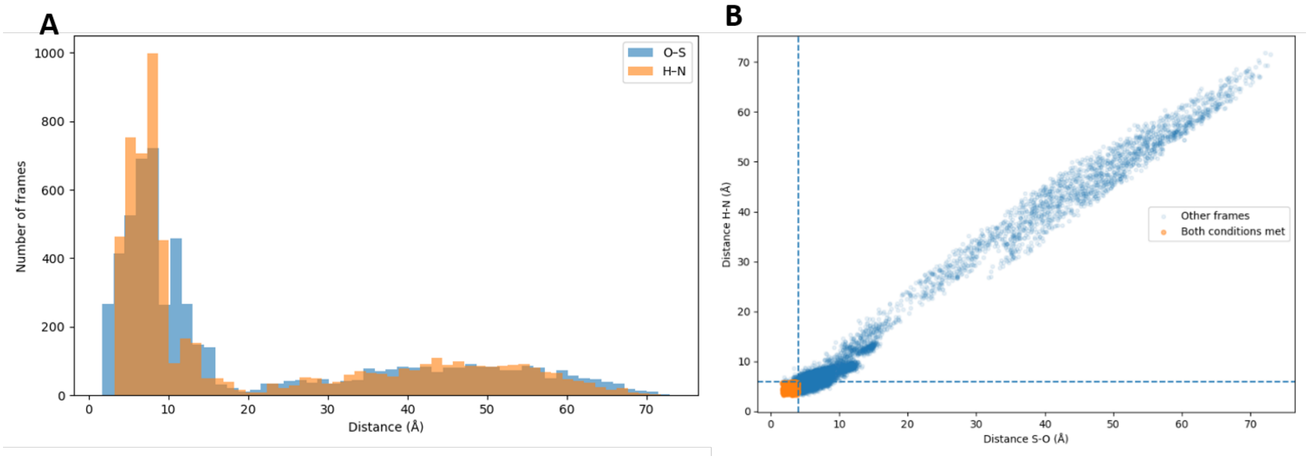


**Figure S19. Distribution and selection of catalytic relevant distances in the phenol-bound MD simulations.** (A) Distributions of the $H_{phenol}$–$N_{His317}$ and $O_{phenol}$–$S_{sulfate}$ distances across the combined phenol trajectory. (B) Scatter plot of the $H_{phenol}$–$N_{His317}$ and $O_{phenol}$–$S_{sulfate}$ distances. Frames satisfying both distance criteria are highlighted in orange.

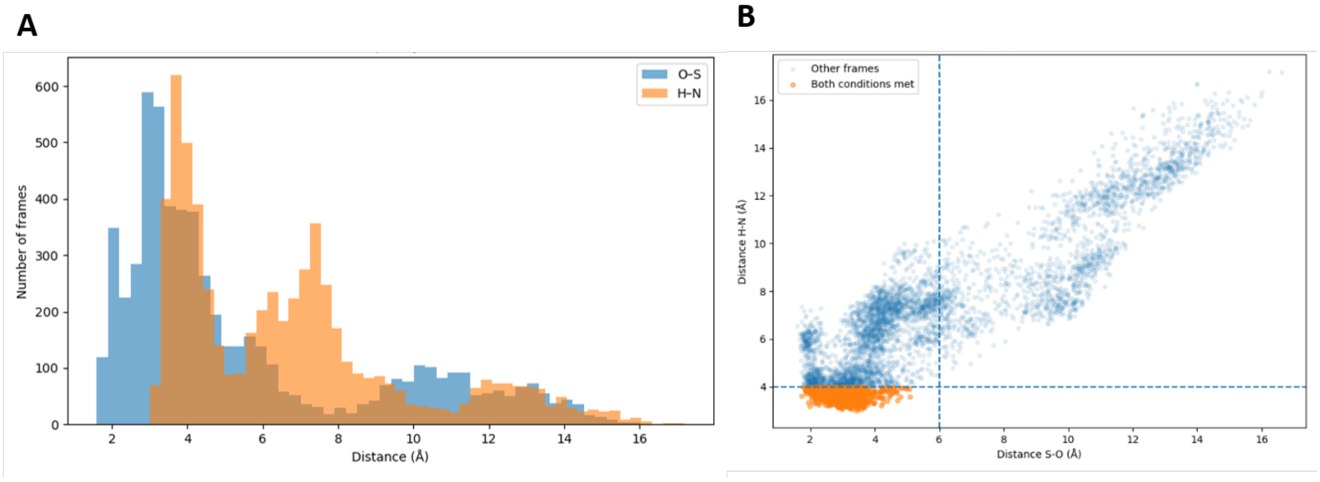


**Figure S20. Distribution and selection of catalytic relevant distances in the dopamine-bound MD simulations**. (A) Distributions of the $H_{dopamine}$–$N_{His317}$ and $O_{dopamine}$–$S_{sulfate}$ distances across the combined dopamine trajectory. (B) Scatter plot of the $H_{dopamine}$–$N_{His317}$ and $O_{dopamine}$–$S_{sulfate}$ distances. Frames satisfying both distance criteria are highlighted in orange.

The MD simulations show that both phenol- and dopamine-bound St_ASST_5 remain structurally stable and sample multiple related backbone conformations. The catalytic distance analysis further shows that only a subset of frames maintains a geometry compatible with both His317-mediated proton abstraction and S–O bond formation with the sulfonated His401 intermediate. These catalytically relevant frames were therefore used to guide the selection of starting conformations for subsequent QM/MM calculations. Dopamine showed a larger fraction of frames fulfilling the selected distance criteria than phenol. This suggests that, once a productive dopamine pose is formed, the reactive catechol hydroxyl group can remain positioned near the His317/His401 catalytic arrangement during MD sampling. In contrast, phenol sampled fewer frames satisfying both criteria, indicating a more restricted set of productive geometries.

#### 8. Umbrella Sampling

##### 8.1 General QM/MM setup

QM/MM umbrella sampling (9) simulations were performed to characterize the second catalytic step of St_ASST_5, in which the sulfate group is transferred from the sulfonated His401 intermediate to the acceptor substrate. Separate simulations were carried out for phenol and dopamine starting from catalytically competent substrate-bound sulfo-enzyme structures.

All QM/MM simulations were performed with Amber22 (5-7) using the sander QM/MM implementation (10). The QM region was treated with DFTB3 (11), using the 3OB parameter set (12,13). The simulations were carried out at 300 K with a time step of 0.0002 ps. Each umbrella window was propagated for 10,000 MD steps, corresponding to 2 ps per window. Coordinates were saved every 100 steps. The nonbonded cutoff and QM/MM cutoff were both set to 10.0 Å. A Langevin thermostat was used with a collision frequency of 1.0 $ps^{-1}$. The total charge of the QM region was +1.

The residue numbering in the simulation topology differs from the manuscript numbering. In the manuscript, the catalytic sulfo-histidine is referred to as His401. In the Amber topology, this residue corresponds to residue 394. All residue names in the text below are therefore given using the manuscript numbering.

| Amber residue index | Manuscript residue | Role in QM region |
|---|---|---|
| **394** | His401 | Sulfonated catalytic histidine / sulfo donor |
| **310** | His317 | Catalytic base for acceptor deprotonation |
| **328** | Arg335 | Active-site charged residue |
| **235** | His242 | Active-site histidine |
| **555** | Phe562 | Aromatic active-site residue |
| **312** | Asn319 | Polar active-site residue |
| **465** | Cys472 | Active-site residue near substrate pocket |
| **466** | Val473 | Active-site residue near substrate pocket |
| **ligand residue** | Phenol or dopamine | Acceptor substrate |

**Table S11. Mapping between Amber topology numbering and manuscript numbering for the QM region.**

The QM region included the acceptor substrate, the sulfonated His401 residue, His317, and selected active-site residues surrounding the reacting groups. For both phenol and dopamine, the protein part of the QM region was kept identical to allow comparison between both acceptor substrates. The ligand was included fully in the QM region. Where covalent bonds between the QM and MM regions were cut (14), link atoms were introduced automatically by Amber (5).

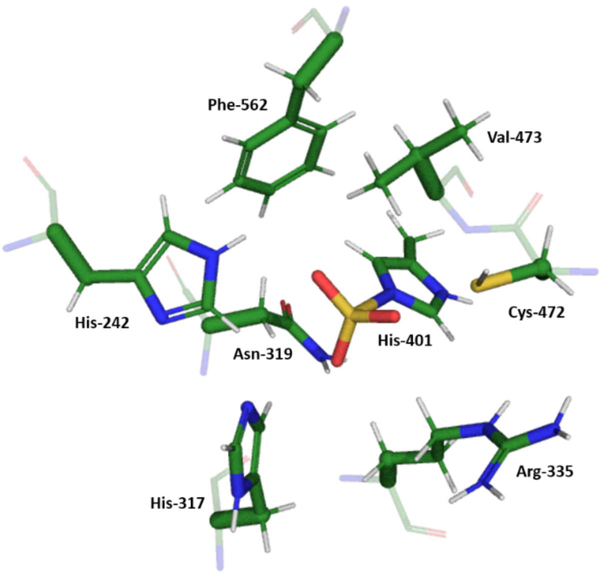


**Figure S21. QM region used for QM/MM simulations of phenol and dopamine sulfo transfer in St_ASST_5.** The atoms included in the QM region are shown as sticks and comprise the the sulfonated His401 intermediate, His317, and selected active-site side chains surrounding the reaction center. Protein backbone atoms, which were not included in the QM region, are shown transparently for structural context. Residue labels follow the manuscript numbering.

8.2 Collective variables and free-energy reconstruction

The reaction was described using two distance-based collective variables. CV1 monitors proton transfer from the substrate hydroxyl group to the catalytic His317 nitrogen, while CV2 monitors formation of the S–O bond between the substrate oxygen and the sulfate sulfur of the sulfonated His401 intermediate:

$$CV1 = d(H_{\mathrm{sub}}\text{-}N_{His317}) \quad (1)$$

$$CV1 = d\left(O_{\mathrm{sub}}\text{-}S_{sulfate}\right) \quad (2)$$

Thus, CV1 follows acceptor deprotonation, whereas CV2 follows sulfuryl transfer. These two coordinates describe the coupled proton-transfer and S–O bond-forming process during conversion of the sulfo-enzyme intermediate into the sulfated product.

The umbrella windows were generated along a coupled path between the reactant-like and product-like geometries. Harmonic distance restraints were applied to both collective variables in each window using Amber NMR-style restraints (10). The production setup used rk2 = rk3 = 100 kcal·mol$^{-1}$ Å$^{-2}$ for both distance restraints in Amber notation. The biased trajectories were analyzed by extracting CV1 and CV2 from each umbrella window. The two-dimensional free-energy surface was reconstructed from the sampled CV distributions using an MBAR/KDE-based analysis (15,16). The free-energy surface was shifted so that the minimum free-energy value was set to 0 kcal·mol$^{-1}$. A one-dimensional PMF was then obtained by projecting the two-dimensional free-energy surface onto the sampled reaction path. The activation free energy was determined as the difference between the reactant minimum and the highest point along this path.

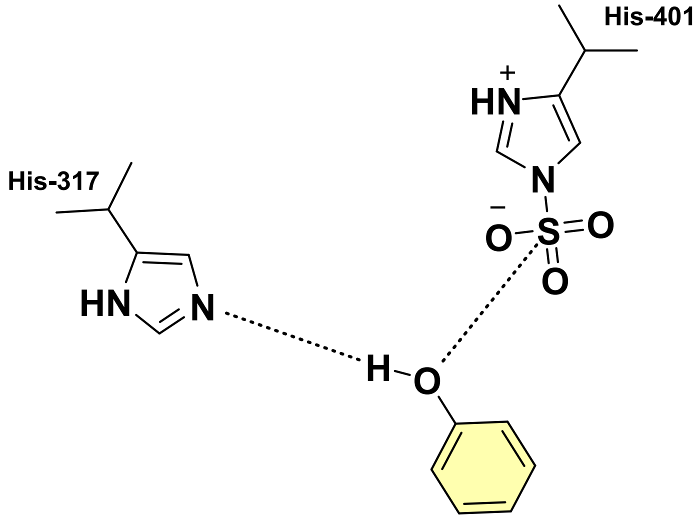


**Figure S22. Definition of the collective variables used for QM/MM umbrella sampling.** CV1 corresponds to the $H_{\text{sub}}$–$N_{His317}$ distance and monitors proton transfer from the substrate hydroxyl group to His317. CV2 corresponds to the $O_{\text{sub}}$–$S_{sulfate}$ distance and monitors formation of the S–O bond between the acceptor oxygen and the sulfate sulfur of the sulfonated His401 intermediate.

#### 9. Phenol QM/MM umbrella sampling

##### 9.1 Umbrella window setup for phenol

For phenol, 25 umbrella windows were generated along the coupled proton-transfer and S–O bond-formation pathway. CV1 was defined as the distance between the phenolic hydroxyl proton and the His317 nitrogen. CV2 was defined as the distance between the phenolic oxygen and the sulfate sulfur of the sulfonated His401 intermediate.

For the phenol system, CV1 used atoms 8685 and 4765, and CV2 used atoms 8684 and 6074. The CV1 target distance was changed from 1.8266 Å in the reactant-like region to 1.0000 Å in the product-like region. The CV2 target distance was changed from 3.6647 Å to 1.6000 Å. The 25 umbrella windows were generated by linear interpolation between these endpoint values.

| Parameter | Value |
|---|---|
| **Number of umbrella windows** | 25 |
| **Sampling per window** | 2 ps |
| **MD steps per window** | 10,000 |
| **Time step** | 0.0002 ps |
| **Saved frames per window** | 100 |
| **CV1** | $H_{phenol}$–$N_{His317}$ |
| **CV2** | $O_{phenol}$–$S_{Sulfate}$ |
| **CV1 atom pair** | 8685–4765 |
| **CV2 atom pair** | 8684–6074 |
| **CV1 target range** | 1.8266 → 1.0000 Å |
| **CV2 target range** | 3.6647 → 1.6000 Å |
| **Restraint force constant** | rk2 = rk3 = 100 kcal·mol$^{-1}$ Å$^{-2}$ |

**Table S12. Umbrella sampling setup for phenol.**

9.2 Sampling overlap and free energy reconstruction for phenol

The sampled CV distributions followed the intended reaction pathway from the reactant-like region to the product-like region. Neighboring windows showed continuous sampling along both CV1 and CV2, supporting reconstruction of the free-energy profile along the coupled reaction coordinate.

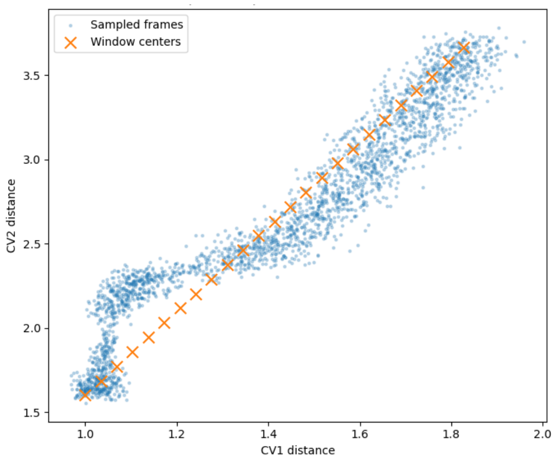


**Figure S23. Sampled CV space for phenol umbrella sampling.** Scatter plot of all sampled CV1 and CV2 values from the phenol umbrella windows. The umbrella window centers are shown as markers. The sampled points follow a continuous path from the reactant region to the product region.

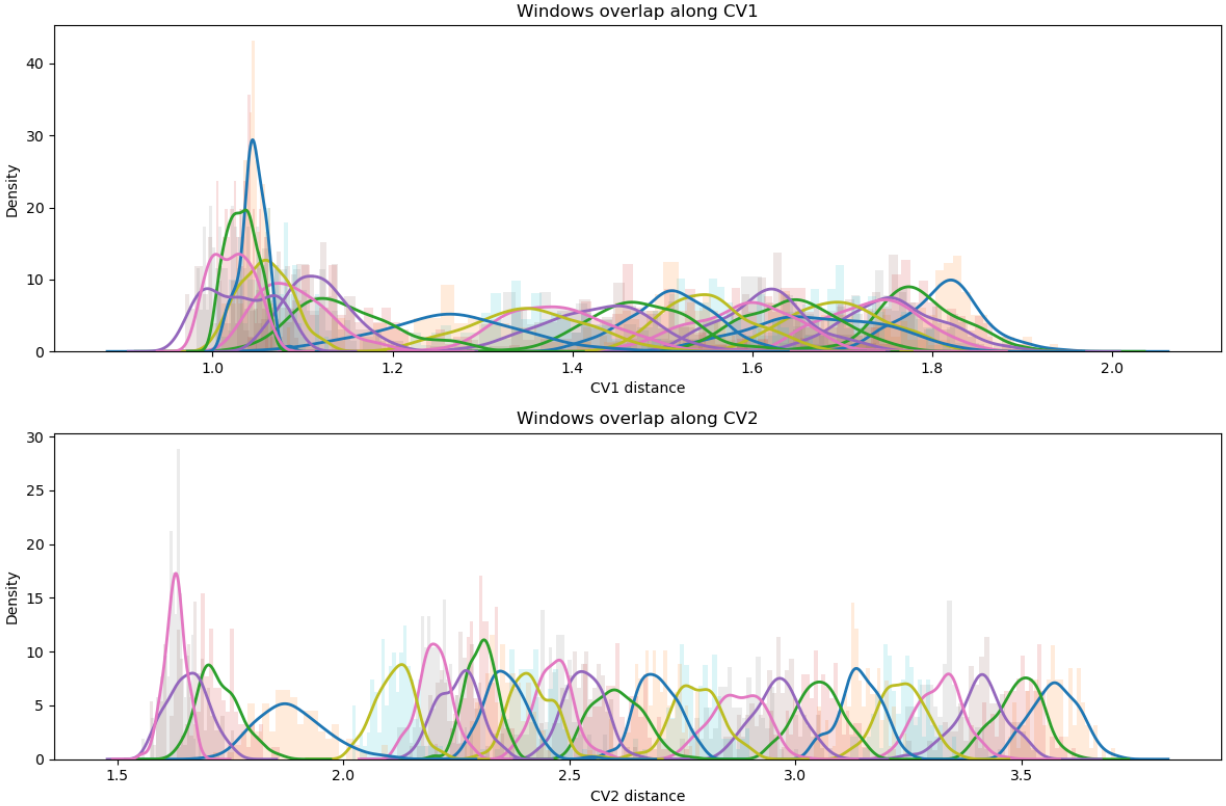


**Figure S24. Umbrella-window overlap for phenol QM/MM umbrella sampling.** Distribution of sampled configurations in the phenol umbrella sampling simulations along CV1 and CV2. CV1 describes the proton-transfer coordinate between the phenolic hydrogen atom and the catalytic nitrogen atom of His317, whereas CV2 describes formation of the bond between the phenolic oxygen atom and the sulfur atom of the His401-bound sulfo group. The histograms and density curves show the sampling distribution of individual umbrella windows. The observed overlap between neighboring windows supports reconstruction of the one- and two-dimensional PMF profiles for the phenol sulfo-transfer reaction.

#### 9.3 Representative phenol structures

Representative structures were extracted from the sampled pathway to illustrate the progression of the reaction. These structures correspond to the reactant region, a preorganized near-attack configuration, the transition-state region, and the product region. The key distances are the $H_{phenol}$–$N_{His317}$ distance and the $O_{phenol}$–$S_{sulfate}$ distance.

| State | Description | CV1 / Å | CV2 / Å | Free energy / kcal·mol⁻¹ | Window/ |
|---|---|---|---|---|---|
| **I** | Reactant-like state | 1.820 | 3.608 | 0.00 | 1 |
| **II** | Preorganized near-attack configuration | 1.144 | 2.30 | 2.648 | 17 |
| **III** | Transition-state region | 1.044 | 1.871 | 10.451 | 22 |
| **IV** | Product-like state | 1.023 | 1.650 | 9.237 | 25 |

**Table S13. Representative structures along the phenol sulfation pathway.**

### 10. Dopamine QM/MM umbrella sampling

#### 10.1 Umbrella window setup for dopamine

The same QM/MM umbrella sampling protocol was applied to dopamine. Dopamine contains two catechol hydroxyl groups and an amino group; therefore, the simulations were performed for the catalytically competent O-sulfation pose in which one catechol hydroxyl group was oriented toward the sulfonated His401 intermediate and His317. For the dopamine system, CV1 was defined as the distance between the reactive dopamine hydroxyl proton and the His317 nitrogen. CV2 was defined as the distance between the corresponding dopamine oxygen and the sulfate sulfur of the sulfonated His401 intermediate. CV1 used atoms 8701 and 4765, and CV2 used atoms 8700 and 6074. The CV1 target distance was changed from 1.8483 Å in the reactant-like region to 1.1000 Å in the product-like region. The CV2 target distance was changed from 3.8528 Å to 1.6000 Å. The initial dopamine setup contained 25 umbrella windows generated by linear interpolation between these endpoint values.

| Parameter | Value |
|---|---|
| **Initial number of umbrella windows** | 25 |
| **Additional refined windows** | 3 |
| **Sampling per window** | 2 ps |
| **MD steps per window** | 10,000 |
| **Time step** | 0.0002 ps |
| **Saved frames per window** | 100 |
| **CV1** | $H_{dopamine}$–$N_{His317}$ |
| **CV2** | $O_{dopamine}$–$S_{Sulfate}$ |
| **CV1 atom pair** | 8701–4765 |
| **CV2 atom pair** | 8700–6074 |
| **CV1 target range** | 1.8483 → 1.1000 Å |
| **CV2 target range** | 3.8528 → 1.6000 Å |
| **Restraint force constant** | rk2 = rk3 = 100 kcal $mol^{-1}$ $Å^{-2}$ |

**Table S14. Umbrella sampling setup for dopamine.**

For dopamine, the initial window distribution showed less continuous overlap in the transition-state region than observed for phenol. Additional refined umbrella windows were therefore introduced in this region to improve overlap between neighboring windows and support a more reliable PMF reconstruction. The refined windows were placed between neighboring windows in the barrier region using midpoint CV values derived from the sampled CV distributions. This refinement was introduced to improve sampling continuity and window overlap, not simply to smooth the PMF.

| Refined window | Window counting | CV1 target / Å | CV2 target / Å |
|---|---|---|---|
| **us_21** | 21 | 1.076 | 2.059 |
| **us_0213** | 22 | 1.061 | 1.971 |
| **us_0214** | 23 | 1.057 | 1.847 |
| **us_0215** | 24 | 1.060 | 1.792 |

| | | | |
|---|---|---|---|
| **us_22** | 25 | 1.051 | 1.741 |

**Table S15. Refined dopamine umbrella windows.**

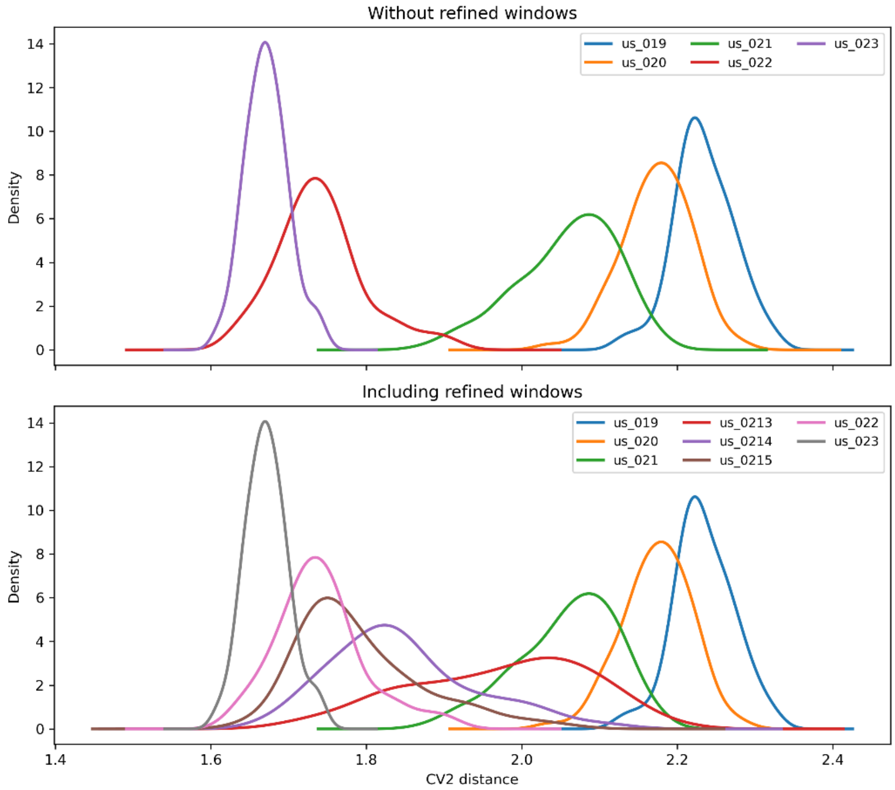


**Figure S25. Refinement of dopamine umbrella-window overlap in the transition-state region.** Comparison of the sampled CV2 distributions for dopamine QM/MM umbrella sampling before and after introducing additional refined windows. The upper panel shows the initial overlap between windows us_019–us_023, where discontinuous sampling is observed in the barrier region. The lower panel shows the improved CV2 overlap after inclusion of additional refined windows placed between neighboring umbrella windows using midpoint CV values derived from the sampled distributions. This refinement increases sampling continuity in the transition-state region and supports a more reliable PMF reconstruction.

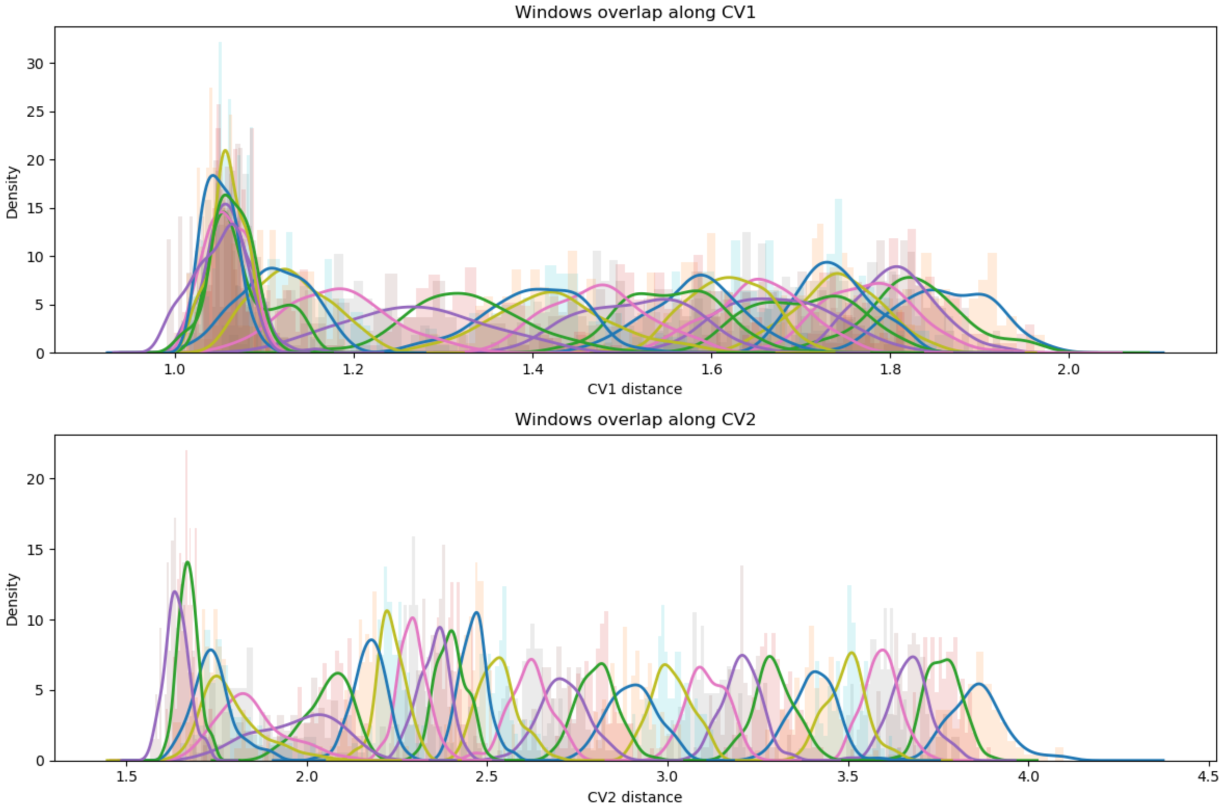


**Figure S26. Umbrella-window overlap for dopamine QM/MM umbrella sampling**. Distribution of sampled configurations in the dopamine umbrella sampling simulations along CV1 and CV2. CV1 describes the proton-transfer coordinate between the phenolic hydrogen atom and the catalytic nitrogen atom of His317, whereas CV2 describes formation of the bond between the phenolic oxygen atom and the sulfur atom of the His401-bound sulfo-group. The histograms and density curves show the sampling distribution of individual umbrella windows. The observed overlap between neighboring windows supports reconstruction of the one- and two-dimensional PMF profiles for the phenol sulfo-transfer reaction.

#### 10.2 Representative dopamine structures

Representative structures were extracted from the dopamine pathway to illustrate the same reaction sequence as observed for phenol: early proton transfer to His317 followed by S–O bond formation from the sulfonated His401 intermediate to the dopamine oxygen.

| State | Description | CV1 / Å | CV2 / Å | Free energy / kcal mol$^{-1}$ | Window |
|---|---|---|---|---|---|
| **I** | Reactant-like state | 1.832 | 3.771 | 0.00 | 2 |

| | | | | | |
|---|---|---|---|---|---|
| **II** | Preorganized near-attack c uration | 1.182 | 2.292 | 3.292 | 20 |
| **III** | Transition-state region | 1.058 | 1.889 | 10.96 | 22 |
| **IV** | Product-like state | 1.047 | 1.641 | 9.847 | 28 |

**Table S16. Representative structures along the dopamine sulfation pathway.**

6.4 Comparison of phenol and dopamine sampling

Phenol and dopamine were analyzed using the same QM/MM protocol, the same protein QM region, and the same two collective variables. This allows direct qualitative comparison of the chemical step once a catalytically competent substrate pose has been reached. Both substrates show early proton transfer to His317 followed by S–O bond formation from the sulfonated His401 intermediate to the acceptor oxygen. The calculated barriers are similar, with $\Delta G^{\ddagger}$ = 10.45 kcal·mol$^{-1}$ for phenol and $\Delta G^{\ddagger}$ = 10.96 kcal·mol$^{-1}$ for dopamine.

| Substrate | CV1 definition | CV2 definition | ΔG‡kcal·mol$^{-1}$ | Mechanistic interpretation |
|---|---|---|---|---|
| **Phenol** | $H_{phenol}$–$N_{His317}$ | $O_{phenol}$–$S_{sulfate}$ | 10.45 | Early proton abstraction followed by S–O bond formation |
| **Dopamine** | $H_{dopamine}$–$N_{His317}$ | $O_{dopamine}$–$S_{sulfate}$ | 10.96 | Same chemical sequence for productive dopamine O-sulfation |

**Table S17. Summary of QM/MM umbrella sampling results.**

The similar barriers indicate that dopamine sulfation is not limited by a substantially higher intrinsic chemical barrier once the catalytically competent catechol orientation is reached. Instead, the main difference between phenol and dopamine is likely associated with substrate positioning and the availability of productive versus nonproductive binding modes.
